# A scalable approach to resolving variants of uncertain significance

**DOI:** 10.64898/2026.02.14.705848

**Authors:** Malvika Tejura, Yile Chen, Abbye E. McEwen, Ross Stewart, Yuriy Sverchkov, Florent Laval, Ivan Woo, Daniel Zeiberg, Runxi Shen, Shawn Fayer, Jeremy Stone, Nahum Smith, Silvia Casadei, Ziyu R. Wang, Matthew W. Snyder, Benjamin J. Capodanno, Pankhuri Gupta, Mariam Benazouz, Shantanu Jain, Sarah Heidl, Lara Muffley, Shengcheng Dong, Benjamin C. Hitz, Idan Gabdank, Khine Lin, Estelle Y. Da, Sabrina Best, Sally Grindstaff, David Reinhart, Leslie Rodriguez-Salas, Obsa Seid, Allyssa J. Vandi, Cameron Wenman, Melinda K. Wheelock, Sriram Pendyala, Dan Holmes, Alicia Xu, Airi Hosokai, Maxime Tixhon, Chloe Reno, Jessica D. Ewald, Kerstin Spirohn-Fitzgerald, Tanisha Teelucksingh, Tong Hao, Zitong S. Chen, Marzieh Haghighi, Ahmad Kamal Hamid, Esteban A. Miglietta, Erin Weisbart, Georges Coppin, Luke Lambourne, Marinella Gebbia, Atina G. Coté, Warren van Loggerenberg, Kirby M. Fawcett, Robert D. Steiner, Jill M. Johnsen, Andrew B. Stergachis, Lilia M. Iakoucheva, Shantanu Singh, Beth A. Cimini, Frederick P. Roth, Richard G. James, the IGVF Coding Variants Focus Group, Marc Vidal, Mikko Taipale, Anne E. Carpenter, Michael A. Calderwood, Mark Craven, Vikas Pejaver, Alan F. Rubin, Predrag Radivojac, Douglas M. Fowler, Lea M. Starita

## Abstract

Over 90% of missense variants across ∼4,000 disease-associated genes are variants of uncertain significance (VUS). Experimental variant effect measurements provide critical evidence about pathogenicity and inform disease biology, but most variants lack data and clinical translation has been limited. The Impact of Genomic Variation on Function Consortium generated experimental data for 62,215 variants across ten genes using multiplexed assays and 1,407 variants across 163 genes using arrayed assays, curated 193,139 additional community-generated variant effect measurements across 30 additional genes, and developed automated calibration methods for translating experimental data and variant effect predictions into clinical evidence. To reduce current VUS, we developed a scalable workflow using only experimental and predictive evidence, enabling reclassification of 75% of the 16,115 VUS in these genes as pathogenic or benign with <1% error. To minimize future VUS, we analyzed >90,000 unobserved variants; 62% had enough evidence to be “preclassified” as pathogenic or benign. We validated our data, evidence and classifications using All of Us and created interactive resources to enable clinical use of the calibrated data. Thus, for 40 genes, representing 1% of the clinical genome, we resolve most existing VUS and future variants, illustrating how systematic use of scalable evidence can empower genomic medicine.

## Main

Genetic testing has identified millions of unique single nucleotide variants (SNVs) in genes associated with disease^1,2^. However, most of these variants have never been functionally characterized, and over 90% of missense and near-intronic variants are classified as variants of uncertain significance (VUS)^3^. VUS cause confusion and distress, and cannot be used in diagnosis or treatment, therefore representing a major barrier to genomic medicine. Moreover, individuals from ancestries historically underrepresented in genomic medicine and research bear a disproportionately large proportion of VUS, perpetuating inequities^4–8^. In 2020, the NHGRI predicted that by 2030 the term’VUS’ would become obsolete and that individuals from ancestrally diverse backgrounds would benefit equitably from advances in human genomics^9^. Making this prediction a reality requires understanding whether variants affect gene and protein function and knowing how variants impact function, including by altering splicing, disrupting protein folding, destabilizing complexes, perturbing molecular interactions, or rewiring cellular networks.

Experimental assays and computational variant effect predictions are the most scalable sources of knowledge about how variants affect gene and protein function. In particular, Multiplexed Assays of Variant Effect (MAVEs) leverage high-throughput DNA sequencing to enable measurement of the impact of nearly all single nucleotide or amino acid substitutions^10–12^. MAVEs for *BRCA1*^13^, *MSH2*^14^, *TP53* ^15,16^ and other clinically relevant genes have demonstrated the power of experimental data to reclassify VUS and reveal mechanisms of disease^3^. MAVE-derived experimental data is comprehensive, and therefore benefits individuals with VUS regardless of ancestry^17^. In parallel, variant effect predictors can estimate the impact of most variants in the genome, and advances in machine learning have increased their accuracy^18–23^. However, use of experimental and predictive data in the clinic has been limited; experimental data is lacking for most genes^24^, both data sources are difficult to translate into clinical evidence^25^, and there is limited dissemination infrastructure^26^.

We hypothesized that systematic generation and application of experimental and predictive evidence would resolve VUS at scale^27–29^. To enable us to test this hypothesis, as part of the NHGRI’s Impact of Genomic Variation on Function (IGVF) Consortium^30^, we:

● Established production-scale MAVEs to systematically characterize all possible missense and near-intronic variants in clinically relevant genes, generating high-quality experimental data for 62,215 variants across 10 genes^12,13,31–36^ (**Fig. 1a,b**). These data accurately discriminate known pathogenic and benign variants and demonstrate disease association for loss-of-function variants in a population biobank.
● Developed high-throughput arrayed phenotypic assays that reveal how 1,407 variants in 163 genes impact function, including protein abundance, protein-protein interactions, and subcellular localization. These results established protein mislocalization as a hallmark of pathogenic variants and prioritized phenotypes for future MAVEs^37^ (**Fig. 1a,b**).
● Integrated our IGVF-generated experimental data with data from 68 published assays across an additional 30 clinically relevant genes and scores from top-performing variant effect predictors^18,20,21,38,39^ (**Fig. 1c,d**). This integrated variant effect dataset contains variant effect measurements and predictions for 255,354 variants and enables systematic variant classification.
● Developed automated calibration methods to translate experimental and predictive data into standardized evidence compatible with established guidelines for clinical variant classification^33,40–42^(**Fig. 1e**). These new methods increase the accuracy of experimental and predictive evidence^25,43,44^ and enable the classification of more variants (**Fig. 1f**).
● Enabled discovery of our calibrated experimental evidence through the IGVF catalog^41^ and MaveMD, a clinically-focused interface for MaveDB^26,38,45^ (**Fig. 1g**). Additionally, we present PredictMD, a platform which enables discovery of calibrated predictive evidence. These resources facilitate exploration of the evidence we provide, enabling variant classification by clinicians.

**Fig. 1:**
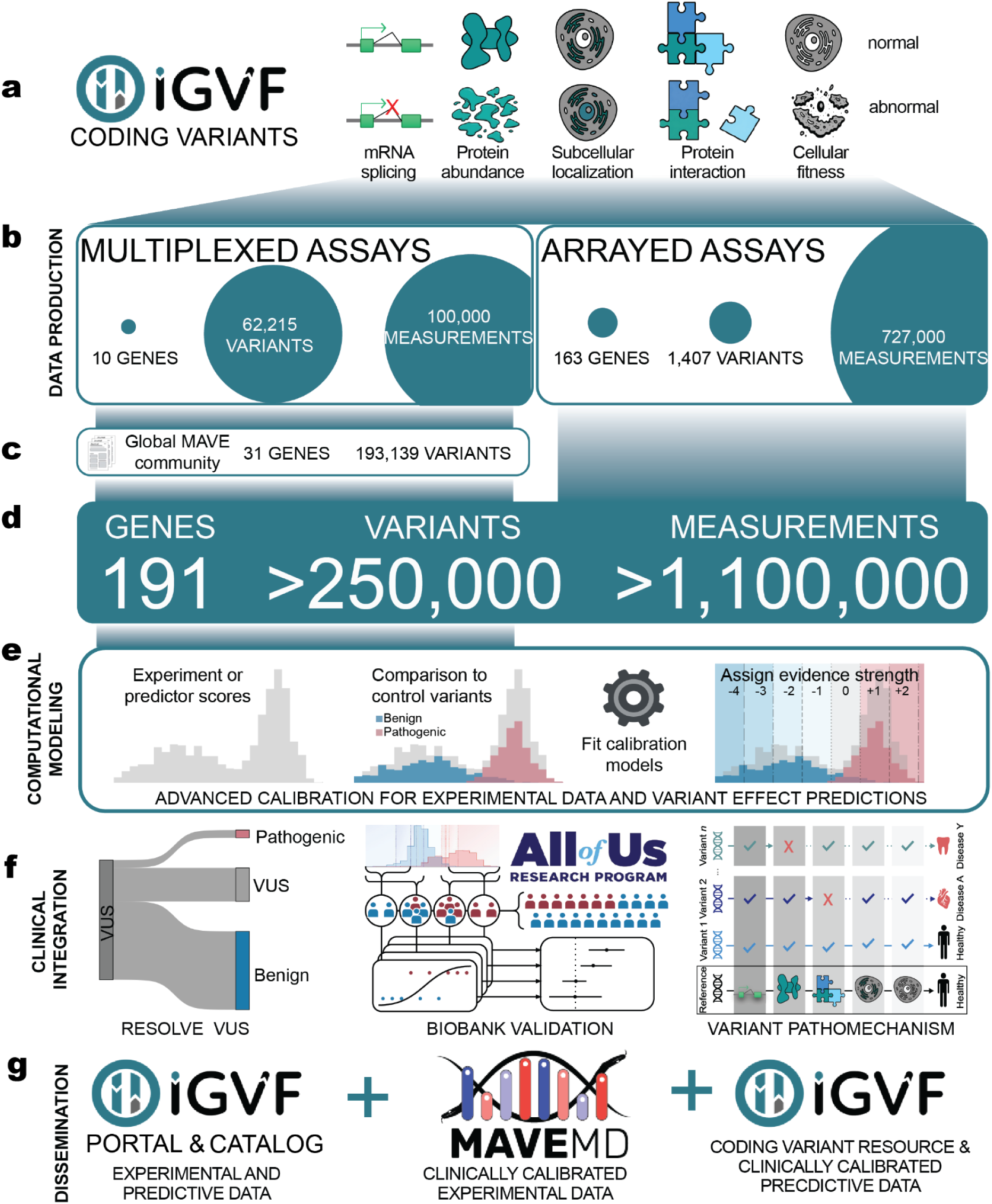
Integrated framework for experimental and computational characterization of coding variant effects (a) The IGVF Coding Variants Focus Group characterizes variant effects across diverse functional readouts including mRNA splicing, protein abundance, subcellular localization, protein-protein interactions, and cellular fitness. (b) IGVF data production teams generated experimental data using multiplexed assays measuring 60,000 variants across 10 genes (100,000 measurements) and arrayed assays measuring 1,407 variants across 163 genes (600,000 measurements). (c) Integration with curated experimental data from the global MAVE community expands data coverage. (d) The integrated variant effect dataset comprises over 1 million functional measurements across 262,000 variants in 200 clinically relevant genes. (e) IGVF computational modeling teams developed advanced calibration methods for both experimental data and variant effect predictions. (f) The integrated variant effect dataset enables systematic VUS resolution, biobank validation of variant classifications in diverse populations, and mechanistic characterization of variant pathomechanisms. (g) Data is disseminated through three complementary resources: the IGVF Portal (raw experimental and predictive data), IGVF Catalog (browsable experimental and predictive data), and MaveMD (clinically calibrated experimental data for immediate clinical use).

Leveraging our experimental data and calibrations, we establish the first scalable workflow for resolving VUS, capable of providing definitive classifications for 75% (12,135 of 16,115) of VUS in the 40 clinically relevant genes we study. To minimize future VUS, we preclassify 90,530 previously unobserved missense variants in these genes, with 62% receiving sufficient evidence to be classified as pathogenic or benign once they are observed. Our workflow results in <1% of known pathogenic or benign variants receiving discordant classifications with ClinVar, and our pathogenic evidence and classifications are strongly associated with disease in a population biobank. Thus, comprehensive functional characterization of genetic variants can simultaneously advance our understanding of gene and protein biology while enabling clinical variant classification at scale, helping make VUS obsolete and enabling equitable precision healthcare.

### Production-scale MAVEs provide variant effect measurements for 62,215 variants across 10 genes

We applied two complementary MAVEs to systematically measure variant effects across 10 clinically relevant genes. Variant Abundance by Massively Parallel sequencing (VAMP-seq) measures steady-state protein abundance for all possible amino acid substitutions by fusing variants to GFP, followed by fluorescence-activated cell sorting and sequencing^12^ (**Fig. 2a**). VAMP-seq is effective for identifying pathogenic missense variants because a primary mechanism of missense pathogenicity for most proteins is reduced steady-state abundance due to protein destabilization and degradation^46–49^. Saturation Genome Editing (SGE) measures the effect of SNVs in essential genes on cellular fitness in haploid human cells, and thus detects the effects on RNA expression, splicing, and protein function in a single assay^11,13^ (**Fig. 2b**).

**Fig. 2:**
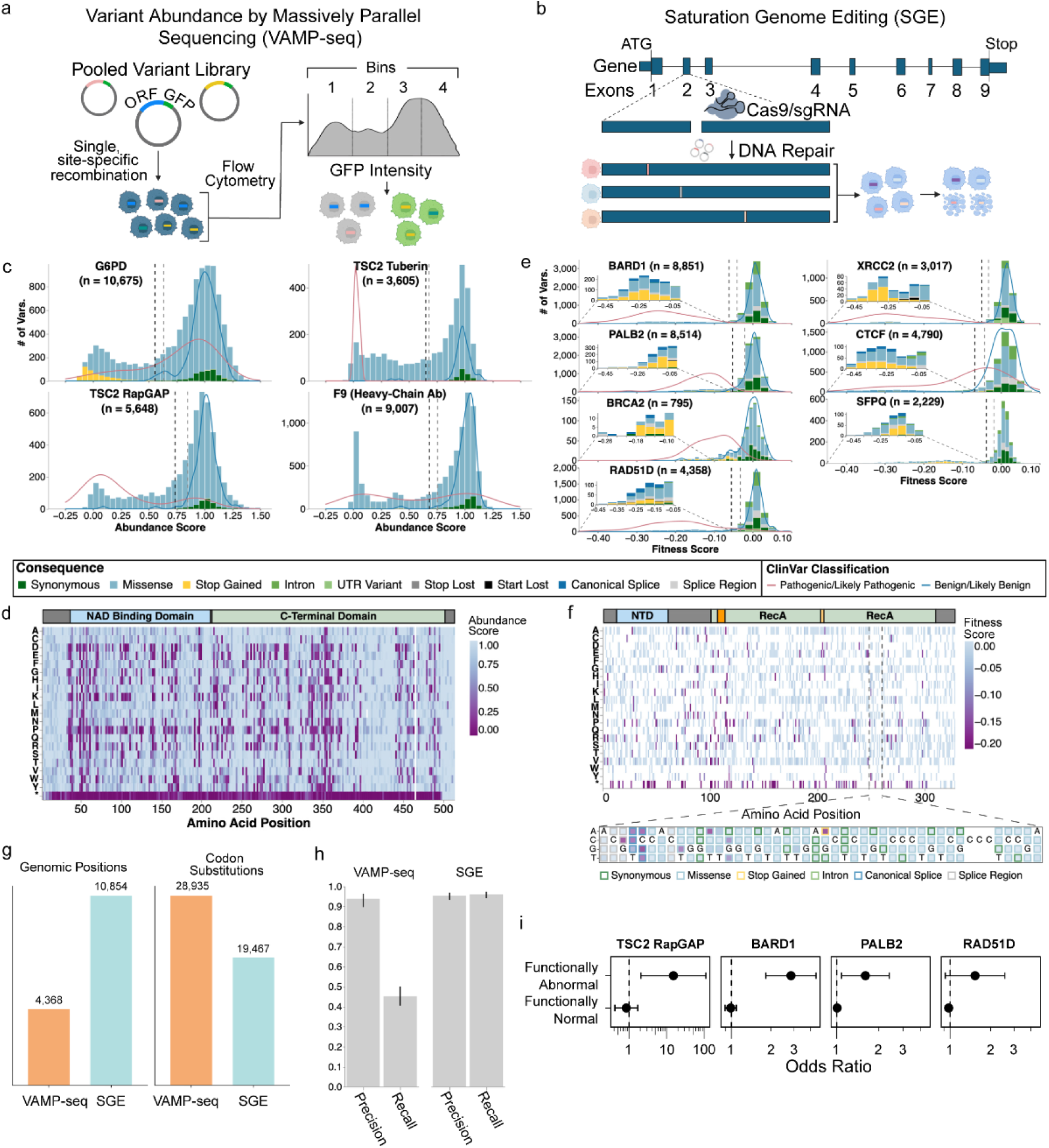
IGVF-produced MAVE data for 62,000 variants (a) Schematic of experimental workflow for Variant Abundance by Massively Parallel Sequencing (VAMP-seq) (b) Schematic of experimental workflow for Saturation Genome Editing (SGE) (c) Histograms showing the distribution of VAMP-seq scores for the indicated genes and domains. The molecular consequence of variants are colored as indicated. Black and gray dashed vertical lines indicate thresholds for variants called functionally abnormal or functionally normal respectively. Overlaid density plots show the distribution of pathogenic/likely pathogenic (red) and benign/likely benign (blue) from ClinVar. (d) Heatmap representation of VAMP-seq scores for G6PD. A cartoon of G6PD domains is at the top. Amino acid positions in G6PD are on the X-axis and amino acid and stop-gained (*) substitutions are indicated on the Y-axis. Abundance scores are colored as indicated. (e) Histograms showing the distribution of SGE fitness scores for the indicated genes. The molecular consequence of variants are colored as indicated. Black and gray dashed vertical lines indicate thresholds for variants called functionally abnormal or functionally normal respectively for datasets with at least 1,000 variants scored. Insets highlight functionally abnormal variants or all variants scoring <-0.10 for datasets without thresholds. Overlaid density plots represent the distribution of pathogenic/likely pathogenic (red) and benign/likely benign (blue) from ClinVar for each gene with at least five PLP and BLB variants. (f) Amino acid-level heatmap of SGE fitness scores for missense and stop-gained variants in RAD51D with a cartoon of RAD51D protein domains (top). A nucleotide-level heatmap for the indicated region of *RAD51D* (bottom). At each position, the wild-type base is written and alternate bases are denoted on the Y-axis. Colored borders for each variant indicate the molecular consequence and fitness scores for each variant are filled as indicated. (g) Bar plot comparing the number of nucleotide positions (left) targeted by VAMP-seq (orange) and SGE (teal) across genes and number of unique codon substitutions (right) assayed by VAMP-seq and SGE. (h) Bar plot showing the precision and recall for VAMP-seq (right) and SGE (left) datasets on known pathogenic/likely pathogenic ClinVar variants. Black vertical lines at the end of each bar denote a 95% confidence interval. (i) Odds ratios for occurrence of disease in individuals with functionally abnormal and functionally normal variants in the All of Us biobank for the TSC2 RapGAP domain, *BARD1*, *PALB2* and *RAD51D*. X-axis denotes the odds ratio with the black vertical dashed line indicating an odds ratio of 1. Y-axis denotes the functional consequence of variants used in the analysis. Points denote the estimated odds ratio and whiskers denote the 95% confidence interval.

We applied VAMP-seq to three proteins with distinct protein folds and disease associations. For *G6PD*, which encodes glucose-6-phosphate dehydrogenase and where pathogenic loss-of-function variants are associated with increased risk of hemolytic anemia, we measured the abundance of 10,675 variants^35^. For *TSC2*, which encodes the tumor suppressor tuberous sclerosis complex subunit 2, we focused on the tuberin and RapGAP domains where pathogenic missense variants cluster, measuring the abundance of 9,253 variants^34^. For *F9*, which encodes factor IX and where pathogenic variants cause hemophilia B, we deployed a modified VAMP-seq protocol using antibody labeling to make 45,035 measurements of the secretion, folding, and post-translational modifications of 9,007 variants^32^. Together, these VAMP-seq datasets represent 64,963 measurements across 27,097 missense, 1,287 synonymous, and 551 nonsense variants (**Fig. 2c,d; Supplementary Data 1**). Abundance score distributions show clear separation of synonymous variants, which reflect normal abundance, from nonsense and low abundance missense variants (**Fig. 2c**).

We used SGE to measure the effects of SNVs on cellular fitness for seven genes, five of which encode proteins in the homology-directed DNA break repair pathway: *BARD1* (8,851 SNVs)^36^, *PALB2* (8,514), *BRCA2* N-terminus (795), *RAD51D* (4,358), and *XRCC2* (3,743) (**Fig. 2e,f**). In these genes, pathogenic variants are established or suspected risk factors for hereditary breast and/or ovarian cancer^50,51^, leading to VUS accumulation due to the inclusion of these genes on cancer risk panel tests. We also measured the effects of 4,790 SNVs in *CTCF*, required for genome organization, where pathogenic *de novo* variants cause neurodevelopmental delay and autism^52^; and 2,229 SNVs in *SFPQ*, where variants are suspected to cause intellectual disability^53^. Together, these SGE datasets represent 33,280 SNVs across coding exons, exon-intron junctions, and UTRs (**Fig. 2e**). Unlike VAMP-seq, SGE’s single-nucleotide resolution enables assessment of splice variants and loss or gain of start and stop codons (**Fig. 2f**).

All datasets were uniformly processed and met rigorous IGVF quality control standards for variant coverage and replicate correlation (<u>IGVF standards</u>). Together, VAMP-seq and SGE queried 48,402 codon substitutions at 15,222 nucleotide positions in 10 genes, with VAMP-seq covering more variants and SGE more positions (**Fig. 2g**). VAMP-seq saturates amino acid variants, permitting deep exploration of sequence-structure-function relationships whereas SGE saturates single nucleotide variants, capturing splicing and covering more genomic space (**Fig. 2d,f,g**). VAMP-seq shows high precision for identifying pathogenic variants (range = 0.897 - 0.965), while SGE demonstrates both high precision (range = 0.934 - 0.969) and recall (range = 0.942 - 0.974) in discriminating loss-of-function pathogenic variants from benign variants (**Fig. 2h**). We validated these experimental measurements in the All of Us biobank, where participants carrying variants identified as loss-of-function in *TSC2, BARD1*, *PALB2* and *RAD51D* showed elevated risk for cancer phenotypes (**Fig. 2i, Supplementary Data 2**).

## Curating experimental data from the community to generate an integrated variant effect dataset

To augment our experimental data, we assembled 68 community-generated large-scale datasets encompassing 295,058 variant effect measurements and 31,858 composite scores for 193,139 unique variants of 30 additional genes associated with rare diseases, cancer predisposition, metabolic diseases and cardiovascular phenotypes (see **Supplementary Data 3** and references therein). We curated these community data, adding clinically relevant metadata, mapping all variants to genomic coordinates, standardizing variant nomenclature, and integrating them with our newly generated IGVF datasets to create a harmonized experimental dataset that includes more than 400,000 variant effect measurements for 255,354 variants of 40 unique genes^38^. The 40 genes in the dataset are components of 7,885 registered genetic tests^54^ and nearly half (18) are listed as ACMG Secondary Findings genes^55^, indicating clinical actionability (**Fig. 3a**). Experimental data in the set was generated using multiple model systems across three major modalities: reporter assays measuring protein expression or activity, cell-fitness assays, and direct protein-function assays evaluating biochemical or biophysical properties (**Fig. 3b**). VAMP-seq and SGE were the most common assays, and 25% of all variant effect measurements were generated by the IGVF Consortium (**Fig. 3c**).

**Fig. 3:**
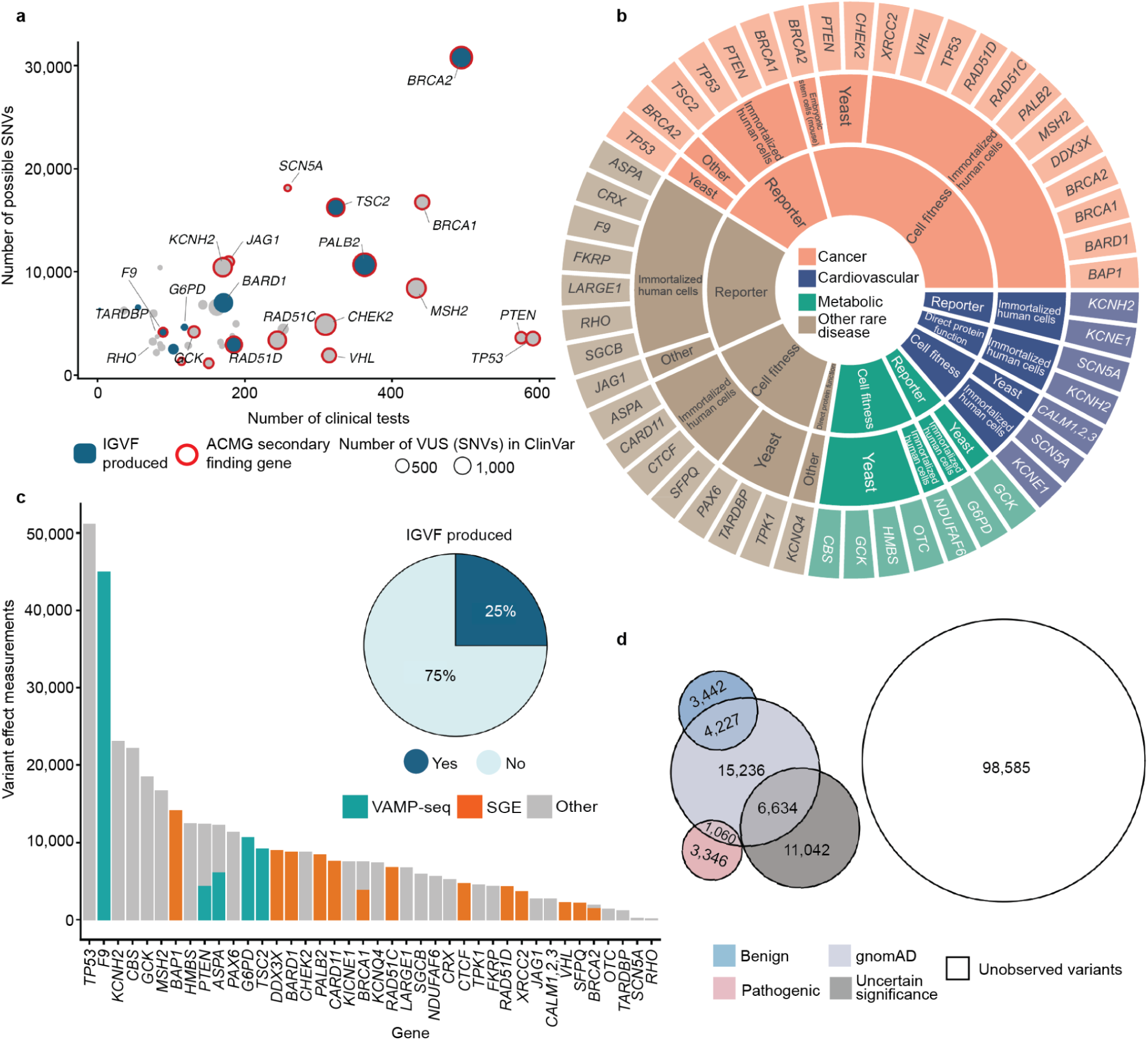
Curating an expanded set of experimental data from the MAVE community (a) Each bubble represents a gene in the integrated variant effect dataset. The number of genetic tests from the Genetic Testing Registry is on the x-axis and the y-axis indicates the total number of possible coding single-nucleotide variants (SNVs). Bubble size is proportional to the number of SNV ClinVar VUS. Bubble color denotes whether experimental data was generated by the IGVF consortium and red outlines denote ACMG secondary findings genes. (b) Sunburst plot summarizing the distribution of experimental data across genes (outer ring), experimental modalities (middle ring) and model systems (inner ring). Segments are colored according to clinical association for each gene as indicated (center). (c) Bar plot showing the number of variant effect measurements (y-axis) per gene (x-axis). Bars are colored by assay type (SGE, VAMP-seq, or other functional assays). A pie chart indicates the proportion of variant effect measurements generated by the IGVF consortium. (d) Venn diagram showing the number of variants that have not yet been observed as well as those in gnomAD or ClinVar (pathogenic/likely pathogenic, benign/likely benign, and VUS colored as indicated).

We annotated variants with scores from AlphaMissense^18^ MutPred2^21^ and REVEL^20^. AlphaMissense and REVEL had accessible scores for non-synonymous SNVs, annotating ∼37% of our variants and MutPred2 had scores for multinucleotide codon changes, annotating ∼77% of our variants. In addition, we annotated variants with ClinVar clinical significance and presence in gnomAD. This integrated variant effect dataset encompasses 17,676 VUS, 12,075 control variants with known pathogenic or benign classifications, 27,157 variants observed in gnomAD and 98,585 possible SNVs that have not been observed in population screening^56^ or genetic testing^1^ (**Fig. 3d**, **Supplementary Data 1**).

### Transforming experimental and predictive data into clinical evidence with improved calibration methods

The first step in using experimental or predictive data for clinical variant classification is calibration, which translates the data into evidence conforming to American College of Medical Genetics/Association for Molecular Pathology (ACMG/AMP) variant classification guidelines^57,58^. In this framework, experimental data yields ‘functional evidence’ and data from variant effect predictors yields ‘computational evidence’. Calibration is based on the ability of the data to discriminate pathogenic from benign control variants and results in each measurement or prediction having a point value ranging from-8, the strongest benign evidence, to +8, the strongest pathogenic evidence. Points from different sources of evidence, including allele frequency, segregation, and patient phenotype as well as experimental and predictive data, are combined to generate a final classification (Benign ≤-6, Likely Benign-5–-1, VUS 0–5, Likely Pathogenic 6–9, Pathogenic ≥10)^59^. However, current calibration methods have shortcomings. For experimental data, the ClinGen-accepted OddsPath method relies on study-specific thresholds that define broad functional categories (e.g., loss of function or functionally normal) without leveraging the full distribution of scores^25^. For predictive data, the accepted ‘genome-wide’ method aggregates variants across disease-associated genes, masking gene-specific predictor behavior and leading to errors in evidence assignment^43,44,60^. We developed new approaches to address these shortcomings.

For calibrating experimental data, we created ExCALIBR (Experimental score CALIBRator). ExCALIBR calculates gene-specific priors from gnomAD frequencies; fits mixture models to assay score distributions of gnomAD, synonymous, pathogenic/likely pathogenic (PLP) and benign/likely benign (BLB) variants; and calculates a posterior probability of pathogenicity for each variant that scales evidence strength based on these distributions^33^ (**Fig. 4a**). We applied ExCALIBR to 37 of the 40 genes in our dataset, excluding *F9* and *TP53* where classifier models were used to integrate multiple datasets^32,61^ and *SFPQ* which remains a candidate disease gene. ExCALIBR assigned at least 1 point of pathogenic or benign evidence to variants of 35 of 37 genes (**Extended Data Fig. 1, Supplementary Data 4**).

**Fig. 4:**
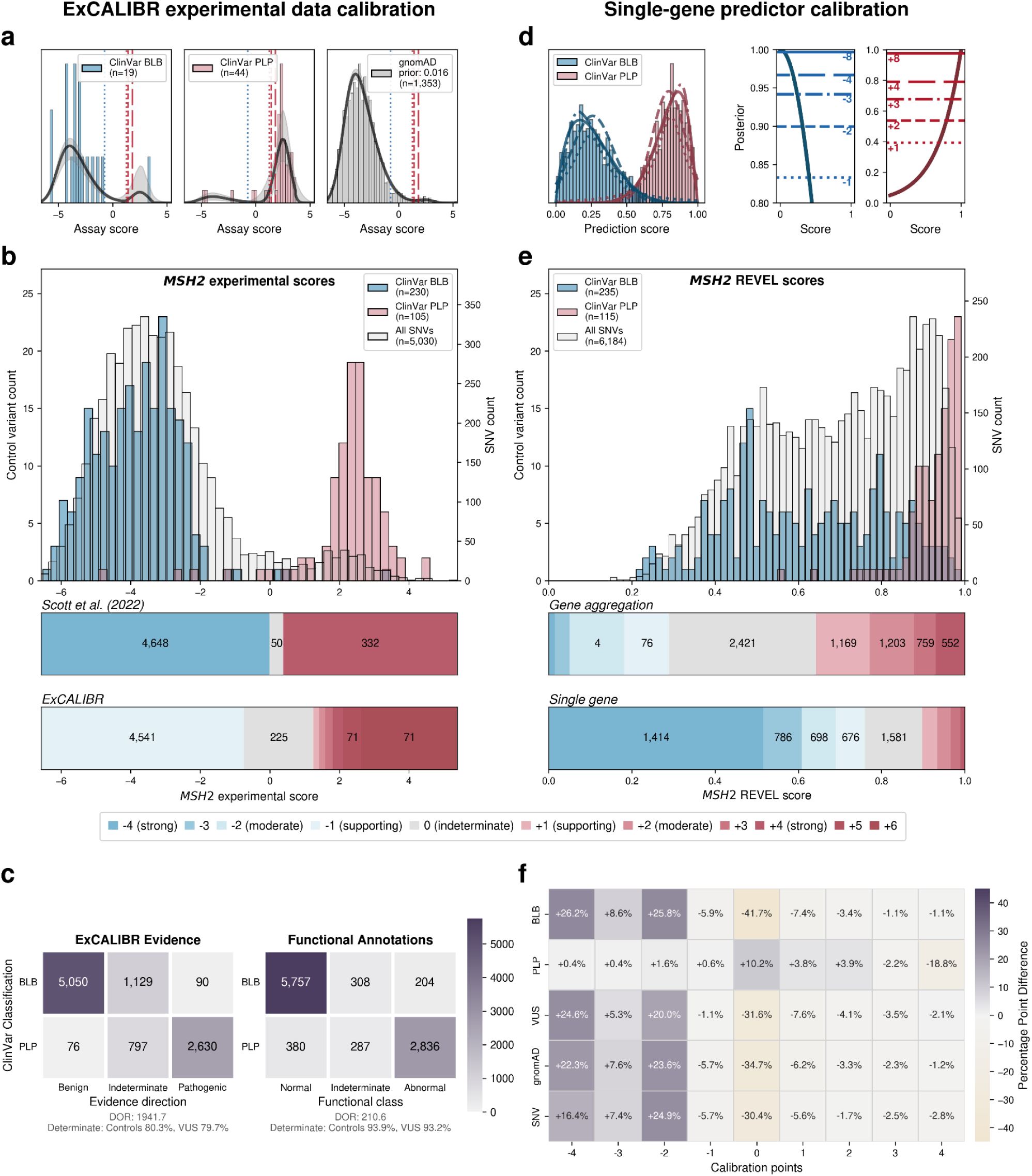
Improved gene-and variant-specific calibration methods improve calibration of experimental and predictive data (a) ExCALIBR experimental data calibration strategy illustrated using *MSH2* data^14^. Score distributions of BLB (blue) and PLP (red) variants from ClinVar (December 2018), as well as population variants from gnomAD (gray), are jointly modeled using skew normal mixtures. The prior probability of pathogenicity is estimated using the gnomAD sample, which is used alongside the local positive likelihood ratio to determine evidence strength thresholds. Evidence strength thresholds are drawn as vertical lines, with red and blue representing pathogenic and benign evidence strength thresholds, respectively (short dashes = 1 point, medium dashes = 2 points, long dashes = 4 points). (b) Comparison of *MSH2* evidence strength thresholds. Top: experimental score distributions for all SNVs (gray), ClinVar BLB (blue) and PLP (red) controls (January 2025). Middle: author-defined evidence thresholds^62^. Bottom: ExCALIBR calibrated thresholds (using December 2018 ClinVar release). Total SNV counts per evidence category are displayed in each bin. (c) Comparison of ExCALIBR evidence assignments (left) and functional annotations (right) for ClinVar PLP and BLB variants for 28 genes with annotated thresholds. For *BRCA1*, *MSH2*, and *PTEN*, ClinVar controls from the December 2018 release were evaluated. Evidence assignments less than zero indicate benign direction; greater than zero indicate pathogenic direction; zero is indeterminate. Diagnostic odds ratios are computed on determinate calls. Determinate assignment rates are shown separately for control variants and VUS. (d) Data-adaptive, single-gene calibration framework. For genes with sufficient control variants, simulation analyses evaluate calibration feasibility and identify the best-fitting gene-specific control distribution. Across 30 simulations and 10 calibration methods, posterior probabilities are estimated. The optimal method is selected based on ACMG/AMP-aligned miscalibration metrics, prioritizing accuracy, robustness, and control of overestimation, and is used to derive calibrated score thresholds and corresponding evidence assignments. (e) Calibration results for *MSH2* REVEL scores. Top: Histogram of REVEL scores for ClinVar PLP (red), BLB (blue), and all possible missense SNVs (gray), with SNV counts shown on the right y-axis. Middle: The number of variants with the indicated evidence assignment for REVEL using the genome-wide^43^or (bottom) for single-gene calibrations. (f) A confusion matrix shows the difference in the percentage of variants assigned evidence points between the gene-specific method and the genome-wide method for all 8 genes^43^ for variants with ClinVar classifications, gnomAD variants and all possible SNVs.

For the *MSH2* dataset^14^, ExCALIBR expands the indeterminate range, better reflecting the mixture of pathogenic and benign control variants compared to the accepted threshold-based approach^62^ (**Fig. 4b**). Moreover, evidence strength increases continuously as variant scores become more extreme, rather than being uniformly strong beyond a fixed threshold (**Fig. 4b**).

ExCALIBR evidence assignments agreed with ClinVar PLP and BLB control variants for 97.9% of variants compared to functional annotations, which agreed for 93.6% of variants (**Fig. 4c**). However, the gain in accuracy comes at the cost of reduced coverage; the proportion of VUS assigned to the indeterminate score range increased from 6.8% to 20.3% (**Fig. 4c**). Nonetheless, because ExCALIBR takes advantage of control variants beyond ClinVar, it was able to calibrate an additional 19 datasets contributing 78,022 variant effect measurements for 48,707 variants that could not be calibrated using the established OddsPath approach. 14 of 17 gene-disease pairs had significant associations between pathogenic evidence and disease phenotypes in the All of Us biobank (**Extended Data Fig. 2a, Supplementary Data 2**). Overall, our calibrated experimental data provides benign evidence for 122,708 variant effect measurements and pathogenic evidence for 74,652 variant effect measurements representing 166,562 unique variants across 35 genes.

For predictive data, we developed a gene-specific calibration method to address the pitfalls of the accepted genome-wide method that we previously identified^60^. Eight of our 40 genes had enough control variants for gene-specific calibration: *BRCA1*, *BRCA2*, *F9*, *JAG1*, *MSH2*, *SCN5A*, *TP53*, *TSC2*^42^ (**Supplementary Data 4**). For each of these genes, we computed gene-specific priors^40,42^ and applied post-hoc calibration via a data-adaptive framework, fitting calibration models to assign posterior probabilities and evidence strength for REVEL, MutPred2 and AlphaMissense (**Fig. 4d, Extended Data Fig. 3**)^42^.

For *MSH2*, REVEL scores are higher overall, causing the accepted genome-wide calibration method to inappropriately assign pathogenic evidence to some variants and no evidence to others (**Fig. 4e**). Our gene-specific calibration aligned the evidence thresholds with the *MSH2* control variant distributions, preventing the misassignment of pathogenic evidence to BLB variants and reducing the number of variants receiving no evidence (**Fig. 4e**). Our gene-specific calibration also yielded odds ratios for cancer association with variants assigned benign evidence in the All of Us Biobank closer to 1 (0.98, 95% CI 0.90-1.07) than the genome-wide method (1.61, 95% CI 0.69-3.78), supporting the superiority of our approach (**Extended Data Fig. 2b, Supplementary Data 2**). When considering all eight genes calibrated using our method, BLB variants misassigned to the indeterminate category was reduced by 41.7% for REVEL, and 30.4% (17,846) more variants were assigned pathogenic or benign evidence^43,44^. Thus, our new calibration methods reduce evidence misassignment, improve the number of variants receiving evidence and move further toward automated calibration by removing subjective assignment of thresholds for both data sources.

### Variant classification workflow using experimental and predictive evidence is highly accurate

We hypothesized that systematically generated, curated, and calibrated experimental and predictive evidence could resolve most VUS in the absence of other evidence. Therefore, we developed a scalable workflow using experimental and predictive evidence to classify variants according to the points-based ACMG/AMP variant classification framework^59^. In particular, we:

● Applied calibrated experimental evidence from single assays, except for *F9* and *TP53* where models were used to integrate data from multiple assays^32,61^, removing variants that had conflicting scores from multiple assays.
● Applied predictive evidence, calibrated with our new gene-specific method where available, and with the genome-wide method for the remainder^43,44^.
● Removed the start-loss variants, variants predicted to affect splicing and variants used in training REVEL and MutPred2 when appropriate.
● Combined experimental and predictive evidence to generate classifications for the remaining 94% (239,836 of 255,354) of variants.

The accepted variant classification workflow involves combining all available evidence including population frequency, segregation, and clinical presentation^57,63–65^. To determine the accuracy of our scalable workflow combining only experimental and predictive evidence we assessed its ability to recapitulate established BLB and PLP classifications from ClinVar that were made with all available evidence^1^. 69% of variants (7,085/10,228) remained consistent with original ClinVar BLB or PLP classifications, and 30% of variants reverted to VUS due to the absence of other evidence (**Fig. 5a**). Critically, our workflow did not result in definitive (i.e. PLP or BLB) classifications that disagreed with ClinVar. The rate of discordance was very low, ranging from 0.66% (67 of 10,228 variants) for classifications using experimental evidence plus predictive evidence from REVEL to 0.69% using evidence from MutPred2 (63 of 9,099 variants) and 1.14% (117 of 10,250 variants) for those employing AlphaMissense evidence (**Fig. 5b,c, Extended Data Fig. 4a-d, Supplementary Data 5**). Of the 67 discordant variants, 62 were PLP to BLB and 5 were BLB to PLP (**Fig. 5b**). The three genes with the most discordant variants were *TP53* (10), *VHL* (8) and *F9* (3) (**Extended Data Fig. 5)**. Thus, our scalable workflow using only experimental and predictive evidence was largely concordant with ClinVar classifications that incorporated all available evidence, was robust across predictors and led to a remarkably low rate of discordance among definitive classifications.

**Fig. 5:**
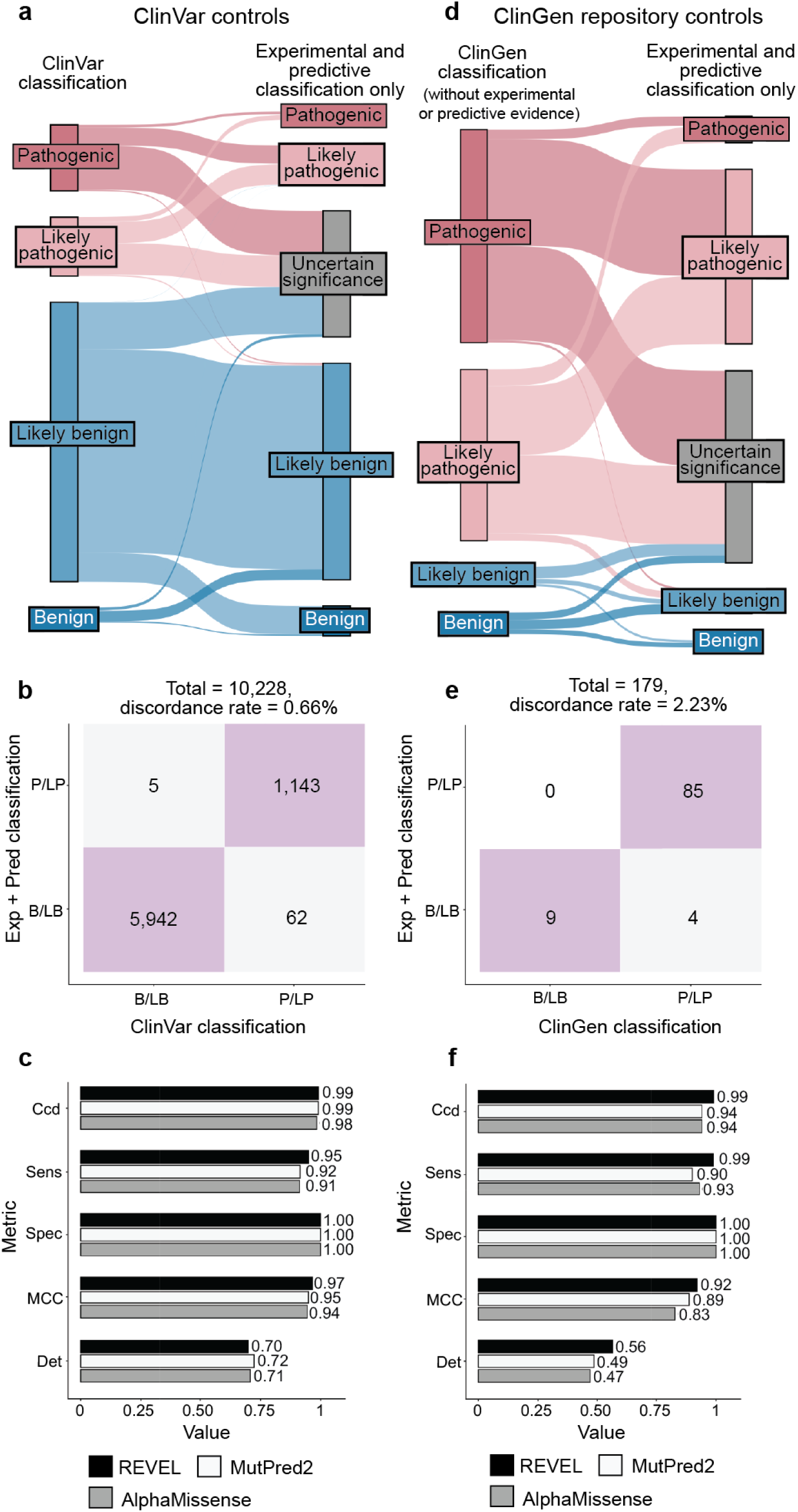
99.3% of PLP and BLB classifications from experimental and predictive evidence match ClinVar (a) Sankey diagram depicting the relationship of the variant classifications in ClinVar and their classifications produced by our scalable workflow using only experimental and predictive (REVEL) evidence. (b) Confusion matrix comparing original ClinVar classifications to classifications from our workflow, summarizing concordant (lilac) and discordant (grey) variant classifications. (c) Bar plots summarizing the concordance of classifications using only experimental and predictive evidence with ClinVar controls. Predictive evidence from REVEL, AlphaMissense, and MutPred2 as indicated. Ccd = concordance, Sens = sensitivity, Sepc = specificity, MCC = Matthew’s correlation coefficient, Det = determinate classifications. (d) Sankey diagram depicting the relationship between variant classifications from the ClinGen Evidence Repository, updated to exclude experimental and predictive evidence (see Methods) to classifications assigned by only experimental and predictive (REVEL) evidence. (e) Confusion matrix comparing updated ClinGen Evidence Repository classifications updated to exclude experimental and predictive evidence to classifications assigned using only experimental and predictive evidence, colored as in (b). (f) Bar plots summarizing the concordance of classifications using only experimental and predictive evidence with updated ClinGen Evidence Repository controls. Predictive evidence from REVEL, AlphaMissense, and MutPred2 as indicated.

We further evaluated the accuracy of our scalable workflow using the ClinGen Evidence Repository, which contains variants annotated with all evidence used (**Fig. 5d**). To avoid circularity, we removed all experimental and predictive evidence that contributed to the original classifications; 194 variants across 11 genes retained sufficient evidence to remain BLB (19) or PLP (175). The resulting ClinGen Evidence Repository classifications that did not depend on experimental or predictive evidence were highly concordant with our classifications using only experimental and predictive evidence, with discordance rates ranged from 2.23% (REVEL) to 2.86% (MutPred2), further validating the low error rate of our scalable workflow (**Fig. 5e,f, Extended Data Fig. 4e-h, Supplementary Note 1 and Supplementary Data 5**).

Finally, we compared variant classifications made using evidence strength computed with our new calibration methods to classifications using evidence strength computed using the currently accepted approaches^25,43,44^. Using accepted calibrations 6,269 (60%) of ClinVar BLB and PLP variant classifications were consistent with the original classification, 39% reverted to VUS and 99 (0.95%) were discordant (**Extended Data Fig. 6a-f, Supplementary Data 6**). 81 (45.76%) of ClinGen Evidence Repository variants were consistent and 3 (1.69%) were discordant (**Extended Data Fig. 6g-l**). Calibrations produced with accepted methods performed well when compared to ClinVar classifications, but our new calibrations produced fewer discordant classifications (0.66% vs 0.95%) and fewer VUS reversions (30% vs 39%). Overall, our improved calibration methods enhanced both the accuracy and classification power of experimental and predictive evidence for clinical variant classification.

### Experimental and predictive evidence alone can reclassify 75% of existing VUS and preclassify 62% of unobserved variants

Our scalable workflow using only experimental evidence and predictive evidence from REVEL yielded a discordance rate of 0.66%, less than the uncertainty thresholds acceptable for pathogenic and benign classifications^66^. This accuracy enables, for the first time, systematic variant classification at scale using standardized experimental and predictive evidence. We applied experimental and predictive evidence from REVEL to three categories of variants across 40 genes: VUS identified through genetic testing in ClinVar^1^, variants detected in human populations via exome or genome sequencing and in gnomAD^56^ and all possible missense SNVs not yet observed by population or clinical sequencing (**Fig. 3d**).

For the 16,115 ClinVar VUS in the 40 genes in our study, more definitive classifications would be immediately useful to improve the efficacy of genetic testing. Of these VUS, 12,135 (75%) received sufficient evidence to be classified as BLB or PLP using REVEL as the predictor; 11,184 (69%) reached BLB and 951 (6%) PLP (**Fig. 6a**, **Supplementary Data 5)**. This outcome is consistent both with the expectation that most VUS are benign^4,67–69^ and with our use of the points-based classification scheme^59^, which reduces the evidence required to reach likely benign compared to previous variant classification framework^57^. Variants receiving pathogenic evidence (combined points ≥ 1) had significant disease associations: 10 cancer predisposition genes showed elevated odds ratios for cancer phenotypes, *SCN5A*, *KCNQ4* and *GCK* variants showed significant associations with long QT syndrome, hearing loss and maturity-onset diabetes of the young, respectively (**Fig. 6b, Supplementary Data 2**). VUS resolution rates using MutPred2 and AlphaMissense (78% and 79% respectively) were comparable (**Extended Data Fig. 7**). 82% of variants with sufficient evidence to be classified as PLP had experimental and predictive evidence, whereas 18% had only experimental evidence. 7,223 of 11,184 BLB variants (65%) had evidence from both sources, 3039 (27%) had only experimental evidence and 922 (8%) had only predictive evidence, suggesting that experimental data drove more reclassifications (**Fig. 6c**).

**Fig. 6:**
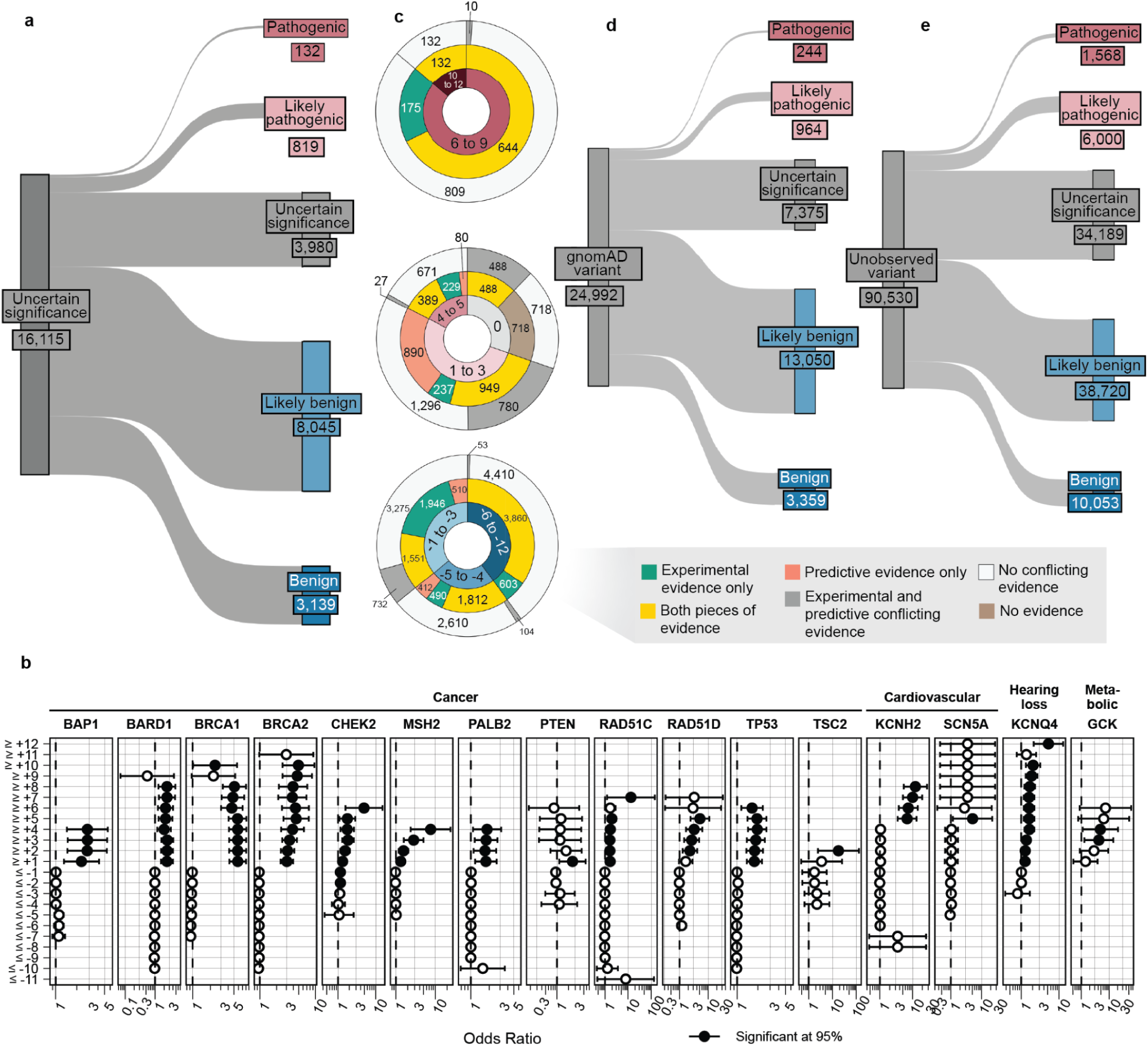
Experimental and predictive evidence can resolve ∼75% of VUS and preclassify 62% of unobserved variants (a) Sankey diagram depicting the relationship of ClinVar VUS and classifications using only experimental and predictive evidence. (b) Odds ratios for occurrence of disease in individuals in the All of Us biobank with variants meeting each indicated evidence strength threshold for genes that were found to have significant association with disease at some level of pathogenic evidence (**Supplementary Data 2**). X-axis denotes the odds ratio with the black vertical dashed line indicating an odds ratio of 1. Y-axis denotes the total evidence points for sets variants. Circles denote the estimated odds ratio and whiskers denote the 95% confidence interval. (c) Donut plots summarizing the strength, source, and consistency of evidence across ClinVar VUS shown in (a). The inner ring denotes the cumulative evidence points assigned to variants. The middle ring indicates the source of evidence as indicated. The outer ring shows the number of variants with concordant (evidence in the same direction) versus conflicting (evidence in opposite directions) evidence for each evidence source category. (d) Sankey diagram depicting the relationship of gnomAD variants to classifications using only experimental and predictive evidence. (e) Sankey diagram depicting the relationship of unobserved variants to classifications using only experimental and predictive evidence.

Of the 16,115 ClinVar VUS, 3,980 (25%) did not receive enough evidence (0 to 5 total points) to be reclassified. 531 (13%) of the unresolved VUS had concordant evidence from both sources, but the evidence was too weak to change the classification. Another 2,154 (54%) of the unresolved VUS were missing one or more sources of evidence: 466 (12%) had only experimental evidence, 970 (24%) had only predictive evidence and 718 (18%) had no evidence from either source. The remaining 33% of the unresolved VUS (**Fig. 6c**) had discordant experimental and predictive evidence, which most often resulted in 0 total points (no evidence). Notably, 698 (17%) of the unresolved VUS accrued 4 or 5 points of pathogenic evidence, suggesting they could reach likely pathogenic classification with additional evidence. The unresolved VUS were distributed unevenly across genes based on the strength of experimental and predictive evidence available for each gene (**Extended Data Fig. 8**). Thus, experimental and predictive evidence alone can reclassify the majority of existing VUS, addressing a substantial portion of variants already discovered through clinical genetic testing.

We also used our scalable workflow to assess the 24,992 variants in our 40 genes in gnomAD. 1,208 (5%) gnomAD variants received sufficient evidence to be classified as PLP, a modest but statistically significant reduction relative to the 6% of ClinVar VUS reclassified as PLP in ClinVar (p = 2.4e-6, chi-squared test, **Fig. 6d**, **Extended Data Fig. 9, Supplementary Data 5**). 16,409 (66%) gnomAD variants received sufficient evidence for BLB classification, a modest but significantly lower percentage than for ClinVar VUS (69%, p = 3.3e-15, chi-squared test). Thus, ClinVar VUS and variants in gnomAD yielded a similar fraction of variants that could be classified as PLP or BLB.

Under the existing variant classification framework, VUS are appearing rapidly^3^. Moreover, all possible SNVs compatible with life will eventually be observed in the human population^70^. Ideally, all newly discovered variants would receive definitive classifications at the time of detection rather than becoming VUS that require reevaluation. To avoid future VUS, we introduce the concept of variant preclassification, where experimental and predictive evidence are systematically generated, calibrated and applied to variants that have not yet been observed in the human population.

We therefore applied our scalable workflow to the 90,530 possible SNVs that have not yet been observed in ClinVar or gnomAD (**Fig. 6e, Extended Data Fig. 9**). Of these, 56,341 (62%) received sufficient evidence to be immediately classified as BLB or PLP. While the overall classification rate of 62% is lower than for ClinVar VUS (75%) or gnomAD variants (71%), a significantly higher proportion were classified as PLP: 8% (7,568) compared to 6% in ClinVar (p < 2.e-16, chi-squared test) and 5% in gnomAD (p < 2e-16). This result is consistent with the findings that variants absent from population databases are more likely to be deleterious^71–76^. We predict that, by providing preclassifications for most variants, application of our scalable workflow to unobserved variants will prevent most new VUS in these genes.

### Identifying protein properties disrupted by variants guides future MAVEs and provides mechanistic insights into pathogenicity

To complement MAVEs and provide broader gene coverage, we developed high-throughput arrayed assays measuring three core protein properties often affected by variants^37,77,78^. Here, variant impacts on protein abundance (DUAL-IPA), subcellular localization (Variant Painting), and protein-protein interactions (sqY2H) were measured for 791 (91 genes), 794 (116 genes) and 502 (74 genes) variants, respectively (**Fig. 7a, Supplementary Data 7**). Altogether, 1,407 variants across 163 genes were experimentally profiled. These assays distinguished between pathogenic and benign variants: 32% of ClinVar PLP variants were abnormal compared to only 9% of BLB variants (p = 2 x 10^-9^, two-sided Fisher’s exact test) (**Fig. 7b**). Each assay independently discriminated pathogenic from benign variants (DUAL-IPA: p = 2 x 10^-7^; Variant Painting: p = 9 x 10^-10^; sqY2H: p = 0.04 Mann-Whitney test) (**Fig. 7c-e**).

**Fig. 7:**
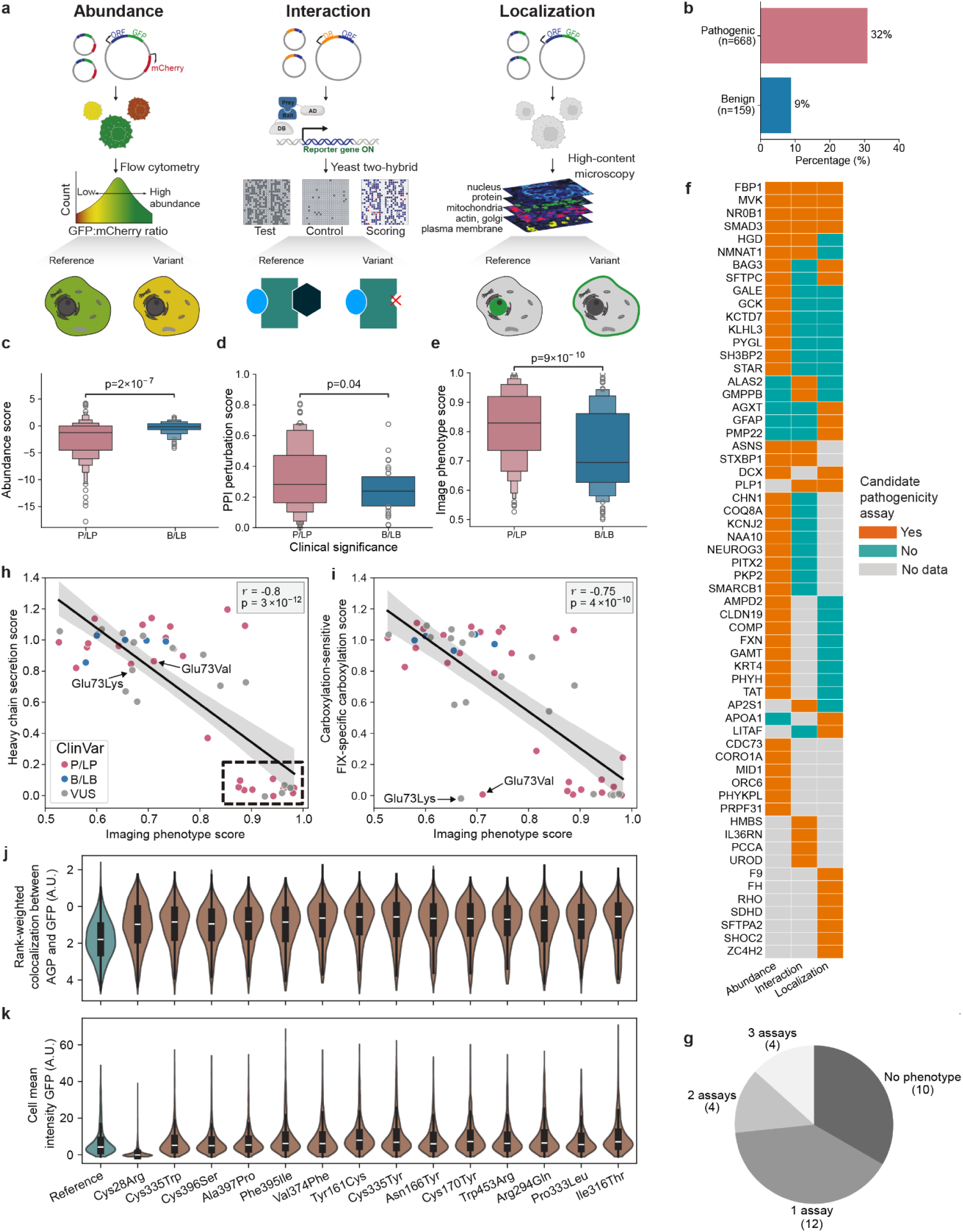
Profiling protein properties to guide future MAVEs and provide mechanistic insights into pathogenicity (a) Schematic of assays for variant effects on abundance (DUAL-IPA), protein-protein interactions (sqY2H) and subcellular localization (Variant Painting). (b) Percentage of variants scoring as hits in 163 genes identified by any of the three assays for PLP and BLB variants. Statistical significance was assessed using Fisher’s two-sided exact test. (c) Distribution of DUAL-IPA abundance scores. Statistical significance was assessed using Mann-Whitney tests. (d) Distribution of PPI perturbation scores. Statistical significance was assessed using Mann-Whitney tests. (e) Distribution of Variant Painting imaging phenotype scores for PLP and BLB variants. Statistical significance was assessed using Mann-Whitney tests. (f) Heatmap showing whether each assay can discriminate PLP from BLB variants and thus serve as a candidate assay for future MAVE development. A gene is assigned a candidate assay if at least one pathogenic variant is identified as a hit in the assay while none of its benign or reference variants are hits. Only genes with at least one candidate assay are shown; other genes are listed in **Supplementary Data 7**. (g) Percentage of genes screened by all three assays that have 3, 2, 1, or no candidate assays. (h) Correlation between imaging phenotype scores and factor IX heavy-chain secretion scores. Boxed variants (n=14) are poorly secreted and show increased Golgi co-localization, indicating secretory pathway retention. (i) Correlation between imaging phenotype scores and factor IX-specific carboxylation score. (j) Distribution of single-cell normalized values for rank-weighted colocalization between AGP staining (actin, Golgi, plasma membrane markers) and GFP-tagged variants (k) Distribution of single-cell normalized values for mean GFP intensity per cell, across the factor IX reference allele and selected variants.

Among the 30 genes profiled across all three assays, 67% exhibited detectable phenotypes for pathogenic variants in at least one assay. Among them, 40% exhibited phenotypes in only one assay, 13% in two assays, and 13% in all three (**Fig. 7f,g**). This phenotypic orthogonality demonstrates that different genes require different assays to detect pathogenic variants, providing a roadmap for prioritizing future MAVE targets and illustrating that pathogenic variants in different genes act via distinct mechanisms.

Integrating arrayed assay data with MAVE results provided insight into variant pathomechanism. For example, we applied a modified version of VAMP-seq to factor IX (*F9*) revealing that 39.5% of missense variants caused reduced secretion of the protein from cells^32^ (**Fig. 2d**). This assay could not distinguish degradation of misfolded protein from intracellular retention. To resolve this ambiguity, we leveraged Variant Painting, a high-content cellular imaging assay that characterizes the localization pattern of missense variants and changes to cellular morphology^37^. Variant Painting-derived phenotype scores, which summarize variant effects across thousands of image-derived features, correlated strongly with VAMP-Seq secretion measurements (**Fig 7h,i**). Individual feature analysis revealed that 14 poorly secreted variants showed increased co-localization with Golgi markers, suggesting retention in the secretory pathway (**Fig 7j**). In contrast, the signal peptide variant p.Cys28Arg showed loss of staining that appeared cytoplasmic, consistent with translocation failure and rapid degradation (**Fig. 7k**). These mechanistic distinctions may have therapeutic implications since total loss of factor IX expression can lead to the development of inhibitory anti-factor IX antibodies in response to exogenous factor IX therapy and worse clinical outcomes^79^. Moreover, variants retained in the secretory pathway represent ‘rescuable’ targets amenable to pharmacological stabilization, while degradation-prone variants may require alternative strategies.

### An ecosystem of resources to disseminate and visualize experimental and predictive data and evidence

Despite the value of experimental and predictive evidence for variant classification, clinical implementation is challenged by several usability barriers^26,80,81^. Experimental data are scattered across supplemental tables and repositories with inconsistent formats, requiring specialized expertise to evaluate quality and clinical relevance. Predictive data is available from multiple sources^82–84^, but the appropriateness of predictive evidence for particular genes and variants is reliant on expert guidance^85,86^. Moreover, no resource allows discovery, assessment, and use of experimental and predictive evidence simultaneously. To address these challenges, we developed MaveMD, to enable the direct clinical use of curated experimental evidence. We also created a dedicated resource for IGVF coding variant data that provides centralized access to the data, analyses and calibrated evidence presented here.

MaveMD (mavemd.mavedb.org) is a clinical interface that transforms experimental data into clinically interpretable evidence^38^. MaveMD implements flexible variant search using clinical notation, interactive histograms contextualizing variant scores within complete functional distributions, and overlays of ClinVar classifications to reveal concordance between experimental data and clinical classifications. Users can easily compare evidence assignments from multiple calibration methods, including ClinGen-specified OddsPath^25^ and ExCALIBR, our improved approach^33^. Standardized “assay facts” labels distill complex metadata into essential criteria, including assay type, molecular phenotype, model system, and variant consequences detected by the assay.

We developed a dedicated resource for IGVF coding variant data (igvf.mavedb.org) that contains all experimental data, calibrations, and analysis tools. We built on the MaveMD framework to present the initial version of PredictMD, which includes calibrated variant effect predictions from three state-of-the-art predictors (REVEL, AlphaMissense, MutPred2) comparing three calibration methods, enabling systematic comparison of variant effect predictions. Together, MaveMD and PredictMD directly address requirements identified in clinical user surveys by enabling detailed, variant-wise comparison of predictions and experimental evidence for the first time, and are designed to substantially reduce technical barriers to routine integration of functional data into clinical practice.

## Discussion

We integrated expertise across experimental genomics, computational biology and clinical genetics to address the VUS challenge. Systematically generated experimental data, combined with calibrated predictions, can resolve 12,135 (∼75%) of VUS across 40 clinically relevant genes. Moreover, we introduce the concept of preclassification, showing that experimental and predictive evidence would enable definitive classification of 62% of the ∼90,000 variants in these 40 genes when they are eventually observed. By establishing production-scale multiplexed and arrayed experimental platforms, developing automated calibration methods, and creating clinically-oriented data resources, we provide both immediate clinical utility and a generalizable framework for eliminating most VUS.

Our production-scale MAVEs generated approximately 25% of extant clinically validated variant effect data. We, along with the wider community^29,87^, established quality standards^39,88^ and demonstrated the feasibility of comprehensive variant characterization across clinically relevant genes. We show that experimental data is required to reach a PLP classification for most variants and that experimental data generally provides more power to classify variants as benign than predictive data. Thus, experimental data for additional genes would directly improve the utility of clinical testing, but new MAVE technologies that increase scale, reduce costs and capture a wider array of phenotypes are needed^89^. This need is highlighted by the gene-specific variation in variant classification accuracy we observed and by our arrayed assays showing that pathogenic variants disrupt multiple functions necessitating diverse assays. Our emerging MAVE platforms dramatically expand scale and mechanistic scope: LABEL-seq measures protein abundance, interactions, and pathway activity at scale^90^, imaging-based assays capture multimodal variant effects and new editing approaches enable assays in physiologically relevant cellular and diploid contexts^37,91–93^.

Our automated calibration approaches represent a fundamental shift from expert-defined thresholds to empirical, unbiased evidence assignment. To calibrate experimental data, we fit mixture models to control variant distributions and calculate gene-specific priors. This approach eliminates subjective priors and thresholds, enabling systematic calibration across datasets^33^. To calibrate predictive data, we developed a gene-specific approach, ameliorating the inappropriate evidence generated for some genes by the previous genome-wide approach^42^. We also developed a promising domain-aggregation approach that groups domains based on similar predictor behavior to expand the number of genes that can be calibrated while increasing calibration power over the genome-wide approach^42^. These automated calibration approaches can keep pace with rapid experimental data production and new predictors while reducing error and improving classification power, although these new approaches should be further vetted by the expert community, including ClinGen Expert Panels.

Our scalable variant classification workflow makes numerous simplifying assumptions, including using only experimental and predictive evidence, discarding variants with conflicting experimental evidence instead of adjudicating these conflicts, ignoring mode of inheritance, and not stratifying ClinVar assertions by mechanism of pathogenicity (e.g., loss-of-function versus gain-of-function) when evaluating concordance. Remarkably, this simplified approach achieves an 0.66% variant discordance rate with ClinVar and 2.23% with ClinGen Evidence Repository control variants. However, our approach does have drawbacks. Using only experimental and predictive evidence led to ∼30% of control variants reverting to VUS. 4.9% (693/13,050) of our BLB reclassifications relied on weak, single-source evidence (-1 point). In the current ACMG/AMP points-based classification system, such weak evidence is sufficient for a likely benign classification, but many of these classifications will be incorrect. For example, an experiment or predictor generating such weak evidence would miss ∼40% of pathogenic variants (**Extended Data Fig. 10 and Supplementary Note 2**). Thus, while these weak, single-source likely benign classifications follow current guidelines, they should be used with caution and ideally augmented with other sources of evidence.

Making the term ‘VUS’ obsolete will require expanding the approach demonstrated here in several dimensions. Experimental data must be produced for thousands of clinically relevant genes, which will require new technologies and expanded data production capacity. Calibration methods must be refined to preserve accuracy with the smaller numbers of control variants available for most genes. New data generated by the community must be curated, harmonized and incorporated into infrastructure that connects experimental and predictive evidence to clinical workflows. Other evidence types must be systematically curated, calibrated and incorporated into a scalable framework. We demonstrate that this investment, if made, could eliminate most current and future VUS.

## Methods

### Multiplexed Assays of Variant Effect

#### Variant Abundance by Massively Parallel Sequencing (VAMP-seq)

VAMP-seq was performed as described in Briar et al. 2026 and Geck et al 2025^34,35^. <u>IGVF</u> <u>standards documents for VAMP-seq</u> include links to protocols.io and quality control metrics. For factor IX, we performed a modified version of VAMP-seq for secreted proteins called MultiSTEP, as described in Popp et al. 2025^32^. <u>IGVF standards documents for MultiSTEP</u> include links to protocols.io and detailed quality control metrics.

#### Saturation Genome Editing (SGE)

SGE was performed as in Woo et al. 2025^36^ with the exception that experiments for *RAD51D* included an extra timepoint at day 17. <u>IGVF standards documents for SGE</u> include links to protocols.io and detailed quality control metrics.

For both Vamp-seq and SGE assays, raw functional assay data were deposited in the IGVF Data Portal (portal.igvf.org), following FAIR principles and consortium data standards, accession numbers are in **Supplementary Table 1**.

#### Arrayed Phenotypic Assays

For all arrayed phenotypic assays, raw functional assay data were deposited in the IGVF Data Portal (portal.igvf.org), following FAIR principles and consortium data standards, accession numbers are in **Supplementary Data 7**.

#### Protein Abundance by DUAL-IPA

To measure variant impact on protein abundance, we developed DUAL-Impact on Protein Abundance (DUAL-IPA), advancing the DUAL-FLUO system^78^. GFP-fused variants are co-expressed alongside a housekeeping mCherry control from a single plasmid in human cells. Flow cytometry quantifies both green and red fluorescences across thousands of cells per variant, normalizing GFP signal to mCherry within each cell to account for cell-to-cell expression variation. The comprehensive experimental protocol is available at: dx.doi.org/10.17504/protocols.io.36wgqpn55vk5/v1

#### Protein-protein interactions

To profile the impact of variants on protein-protein interactions, we adapted the systematic yeast two-hybrid (Y2H) screening pipeline we previously described^94^. Semi-quantitative Y2H (sqY2H) employs 3-Amino-1,2,4-triazole (3-AT) titration to identify optimal stringency where wild-type interactions approach the detection limit, enabling detection of variant-induced weakening. Automated image analysis provides objective, continuous measurement of reporter activation. To ensure data quality, interaction profiles generated by sqY2H were subsequently validated using the mammalian NanoLuc Two-Hybrid (mN2H) protocol we developed^95^. Step-by-step experimental procedures for the semi-quantitative Y2H (sqY2H) and the mN2H validation are available at dx.doi.org/10.17504/protocols.io.bp2l6eknzgqe/v1 78 dx.doi.org/10.17504/protocols.io.rm7vzem8rvx1/v1^78^ 78 To visualize the impact of genetic variants on protein mislocalization, we adapted the previously established Cell Painting V3^37,96^ to create Variant Painting, which relies on the expression of mNeonGreen (mNG)-tagged open reading frames. Changes in protein mislocalization were determined computationally and visually by comparing protein localization of the mutant allele to the corresponding WT reference. ^78^ dx.doi.org/10.17504/protocols.io.36wgqnwkxgk5/v1

### Image Analysis

We used CellProfiler bioimage analysis software (version 4.2.6)^97^ to process the images using classical algorithms following a prior protocol^96^. Illumination variations in images were corrected using the CellProfiler CorrectIlluminationApply module. We segmented nuclei and cells using CellProfiler’s IdentifyPrimaryObjects module with Minimum Cross-Entropy thresholding and IdentifySecondaryObjects module with Otsu three-class watershed thresholding, respectively, and measured feature categories including fluorescence intensity, texture, granularity, density, and location (see http://cellprofiler-manual.s3.amazonaws.com/CellProfiler-4.2.4/index.html for more details) across all segmented compartments and all imaged channels. We obtained ∼3,200 feature measurements from each of about 800 cells per well. We parallelized our image processing workflow using Distributed-CellProfiler [https://doi.org/10.1371/journal.pbio.2005970]. The image analysis pipelines we used are available in the Cell Painting Gallery^98^. (https://cellpainting-gallery.s3.amazonaws.com/index.html#cpg0020-varchamp/).

The overall methods for processing single-cell profiles to produce population-level profiles were developed previously and are standard practice in image-based profiling^99^. The imaging wells with low signal-to-noise ratios (calculated by taking the ratio between 99th-percentile and 50th percentile pixel intensity of the imaging plate) in the GFP (protein localization) channel were removed from downstream analyses. A small number of new steps were incorporated to handle data artefacts unique to single-cell data. First, missing values were handled by filtering out any feature with more than 100 missing values, and then filtering out any cell that still had missing values. Next, we computed plate-level statistics (median and median absolute deviation [MAD]) for each feature and removed features with extremely low variance, defined as having a robust coefficient of variation (MAD divided by the absolute median) below 1e-3 in any plate. Data were then MAD-normalized by plate, which involved subtracting the plate-matched median from each feature value and dividing the result by the plate-matched MAD. After normalization, we filtered out CellProfiler features with extreme values (the 99th percentile is greater than 100) caused by technical artefacts and additionally clipped extreme feature values to-100 or 100. We performed feature selection to remove uninformative and redundant features with Pycytominer using the “variance_threshold” and “correlation_threshold” with default settings. Finally, downstream analysis revealed that occasionally small cell debris was incorrectly identified as cells or many cells were incorrectly identified as one very large cell. These segmentation errors were identified by computing the ratio between the nuclei area and the cell area and filtering to only include cells with an area ratio between 0.1 and 0.4.

#### Protein Mislocalization Analysis

The objective of this analysis was to computationally detect whether a variant allele has a different protein localization pattern than its matched reference allele. We did this by training a set of binary XGBoost classifiers to discriminate between single-cell protein profiles from variant alleles and reference alleles from the same gene. Because the reference and variant alleles were screened on the same plate in separate wells with four replicates each of the two batches, we used a 4-fold cross-validation strategy where samples were stratified by plate. Variant impact was quantified as the model’s AUROC score, with the rationale that variants with greater subcellular localization perturbations would have higher performing classifiers. We refer to the model’s AUROC scores as the Variant Painting-derived phenotype scores in the main text.

Plate layout impacts image-based profiles and therefore single-cell profiles from two different wells can usually be partially distinguished even if the perturbation is the same. Therefore, we needed an estimate of the well position effect to robustly infer which variant-reference classifier scores were greater than expected due to technical effects alone. To estimate the well position effect, we trained an additional set of binary XGBoost classifiers between pairwise combinations of repeated controls on each plate. This resulted in a group of binary classifiers between two wells with different positions on the plate but the same allele per batch. We defined the upper limit of the well position effect as the 95th percentile of the AUROC scores from the repeated control allele thresholds. We defined initial mislocalization hits as variants with variant-reference AUROC scores that surpassed these thresholds.

Finally, we applied various QA/QC filters to call the final mislocalization hits. First, we filtered out any variant-reference pair where there was a class imbalance of more than 3-fold. When there are large differences in cell count, image-based profiles will be easily distinguishable simply due to the differences in cell shape at different densities, and so we couldn’t be confident that hits with large cell count differences actually represented differences in protein localization. Final hits were defined as variants that surpassed the well-position threshold in both biological replicate batches.

### Experimental data curation and integration workflow

#### Gene prioritization and literature review

Experimental data was collected from the published literature through January 2025. Experimental data was prioritized based on the disease association of the gene studied, potential clinical utility of experimental evidence and number of variants experimentally assessed. Disease association was defined using the GenCC (The Gene Curation Coalition) database (downloaded March 28th, 2025)^2^ at moderate or higher disease association. Clinical utility was defined by inclusion in the ACMG Secondary Findings list^55^ or by expert review of gene-disease associations. For each gene, we tallied control variants in ClinVar (pathogenic, likely pathogenic, benign, or likely benign), and genes with more than 20 control variants were prioritized for inclusion. Experimental data curation focused on datasets generated using multiplexed assays of variant effect (MAVEs) as well as experimental assays that tested a large number of single variants, prioritizing datasets with published and accessible raw or composite scores.

### Metadata curation and harmonization

Metadata associated with each assay were curated according to the minimum information standards for MAVE datasets^39^ providing a framework for harmonizing key fields and controlled vocabularies. Specifically, we curated information such as the variant library generation method, cellular or model system used, variant library integration method, and phenotype measured. We further extended this schema to include additional variables specific to our data model, such as the biological mechanism tested, alignment with gene-disease mechanism, and ability to detect splicing and nonsense mediated decay. In total, we curated 68 datasets of which 6 were meta-analyses, consisting of classifier models trained on more than one experimental dataset for a given gene or composite scores derived from the original datasets (**Supplementary Data 3**).

After curating assay-level metadata, we collected and standardized all variant identifiers across datasets using custom scripts. Raw data were retrieved from multiple sources, including supplementary files, GitHub repositories, Zenodo archives, and other public databases (<u>variant</u> <u>score files</u>). Minor manual corrections were made to ensure consistent parsing and are documented in **Supplementary Data 3, column‘Additional Notes’**

Each experimental dataset targeted a specific gene or protein. For each dataset we identified and mapped the corresponding reference transcript using custom scripts to align author-provided protein sequences to reference protein sequences, when not provided by the authors. 11 datasets had a single amino acid discrepancy from the translated reference transcript; these are documented in **Supplementary Data 3, column‘Transcript ID Additional Details’**. Mapping to a reference transcript was critical for subsequent annotations as ClinVar, gnomAD, and computational predictors store genomic coordinates.

Experimental datasets fell into two broad categories:

1. Nucleotide-level assays (*e.g.,* SGE) that introduce variants directly into the endogenous locus, allowing genomic coordinates to be assigned directly.
2. Protein-level assays where variants are expressed as a cDNA (*e.g.,* VAMP-seq) that include single and multinucleotide codon changes or small deletions. These datasets lack genomic coordinates and require mapping to reference transcripts and genomic coordinates.

For protein-level assays, we used transcript identifiers and the standard nuclear genetic code to infer corresponding HGVS annotations (v.21.0.4,^100,101^) for variants. For example, protein variant p.M1V in PTEN corresponds to four possible codon changes encoding valine: ENST00000371953:c.1_3delinsGTT, ENST00000371953:c.1_3delinsGTC, ENST00000371953:c.1_3delinsGTA, ENST00000371953:c.1_3delinsGTG. For each variant in protein-level assays, all possible codon substitutions were compiled in HGVS format (as shown above) and annotated with Variant Effect Predictor (VEP) v113^82^ to add genomic coordinates, variant consequences, and additional sequence-based metadata (https://zenodo.org/records/18637474).

Annotations from VEP^82^ were then integrated with standardized variant identifiers and functional scores. Protein-level datasets that already reported nucleotide changes (BRCA2_Hu_2024, JAG1_Gilbert_2024, RHO_Wan_2019, LARGE1_Ma_2024, FKRP_Ma_2024) were retained as originally reported and were not expanded to all possible nucleotide substitutions to preserve the authors’ intended variant representations. A small subset of variants unable to be annotated by VEP were annotated using VariantValidator^102,103^ (version 3.0.1). Variants that could not be mapped using either VEP or VariantValidator were flagged with “ * “ (**Supplementary Data 1, column‘Flag’**). This process produced a variant effect dataset with harmonized variant identifiers and metadata across 68 published experimental assays (**Supplementary Data 1)**.

### Integration with ClinVar, computational predictors and IGVF experimental data

The harmonized experimental dataset was annotated with data from multiple external resources, including ClinVar classifications^1^ (variant summary files from January 2025 and December 2018 releases), SpliceAI^104^ predictions (precomputed SpliceAI tables based on Gencode v24), gnomAD v4 minor allele frequency^56^, ClinGen Evidence Repository (version 2.5.0) and computational variant effect predictions from AlphaMissense^18^, MutPred2^21^, and REVEL^20^. For REVEL and MutPred2, we recorded which variants appeared in the training sets of these predictors (personal communication from authors). IGVF datasets (BARD1_IGVF, BRCA2_IGVF, CTCF_IGVF,PALB2_IGVF, RAD51D_IGVF, SFPQ_unpublished, XRCC2_IGVF, TSC2_IGVF, G6PD_IGVF, and all F9 datasets) were incorporated and standardized using the same pipeline.

Overall, by combining curated, published datasets with newly generated data, we created an integrated variant effect dataset encompassing 83 experimental datasets across 40 clinically relevant genes, including 15 genes with more than one dataset. This resource comprised 434,166 variant effect measurements for 255,354 unique variants (295,058 measurements for 187,000 variants from published datasets and 31,858 composite scores from published meta-analyses (6,139 variants with composite scores generated from BRCA2_Sahu_2025_SGE); 98,243 newly generated measurements for 62,215 unique variants and 9,007 newly generated composite scores from a meta-analysis (*F9*)).

This integrated variant effect dataset is provided in an expanded format: each row corresponds to a nucleotide-level variant where all variants from protein level assays are mapped to all possible nucleotide substitutions (1,079,133 variants) and the associated protein-level experimental measurements are replicated and applied to each unique nucleotide-level variant (**example** table below) (**Supplementary Data 1**).

| PMID | Gene | Chromosome | Hg38 Start | Hg38 End | Ref allele | Alt allele | HGVS protein | Experimental score |
| --- | --- | --- | --- | --- | --- | --- | --- | --- |
| 29785012 | *PTEN* | 10 | 87864470 | 87864472 | ATG | GTT | p.Met1Val | 1.065143 |
| 29785012 | *PTEN* | 10 | 87864470 | 87864472 | ATG | GTC | p.Met1Val | 1.065143 |
| 29785012 | *PTEN* | 10 | 87864470 | 87864472 | ATG | GTA | p.Met1Val | 1.065143 |
| 29785012 | *PTEN* | 10 | 87864470 | 87864470 | A | G | p.Met1Val | 1.065143 |

### Calibration of experimental and predictive data

#### Calibration of experimental data

##### OddsPath calibrations

As part of our metadata curation, we extracted reported functional classifications determined by the authors representing functionally normal or functionally abnormal classes, and for some published datasets (PTEN_Matreyek_2018, select F9_Popp_2025 datasets) and all newly generated IGVF datasets, created new functional classifications using a Gaussian mixture model as in Woo et al 2025 (^36^. Variants classified as indeterminate or belonging to variant classes not relevant to the disease mechanism under study (e.g. hypermorphic) were annotated but removed from calculations. When more than one functionally normal or functionally abnormal interval or functional classification was provided (ex: ‘slow depleting’ and ‘fast depleting’ for functionally abnormal), they were combined to create one class. We calculated the Odds of Pathogenicity (OddsPath) using ClinVar controls. ClinVar controls were defined as variants with clinical significance of Pathogenic, Likely pathogenic, Pathogenic/Likely pathogenic, Benign, Likely benign, or Benign/Likely benign from the variant summary file. To map these variant classifications to protein-level assays, all possible nucleotide-level substitutions for a given amino acid variant were mapped to ClinVar assertions. When multiple nucleotide substitutions mapped to the same amino acid substitution with concordant ClinVar classifications, the less confident classification was retained (e.g. LP would be chosen instead of P). Amino acid substitutions with discordant nucleotide-level classifications were removed.This resulted in at most one ClinVar assertion per amino acid variant for a given gene and transcript pair (see example below). For *BRCA1*, *TP53*, *MSH2*, and *PTEN*, ClinVar control classifications from 2018 were used to avoid circularity when calibrating or evaluating experimental evidence, as experimental data from these assays may have contributed to ClinVar classifications since their publication^61,62^. For MSH2_Jia_2021 the clinically relevant thresholds from Scott et al. 2022 were used^62^. We followed the framework described by Brnich et al.^25^ including the addition of one misclassified variant per functional class (when necessary) to avoid posterior probabilities of 0 or infinity. Separate values for OddsNormal and OddsAbnormal were derived to assign evidence strength as ACMG/AMP points^59^. Not all experimental datasets included author-defined functional classifications or the thresholds were not provided in an accessible format preventing us from calculating OddsPath. In total, we assigned some evidence (either OddsNormal or OddsAbnormal) to 43 of 83 experimental datasets using the OddsPath calibration method (**Supplementary Data 4, OddsPath calibrations**)

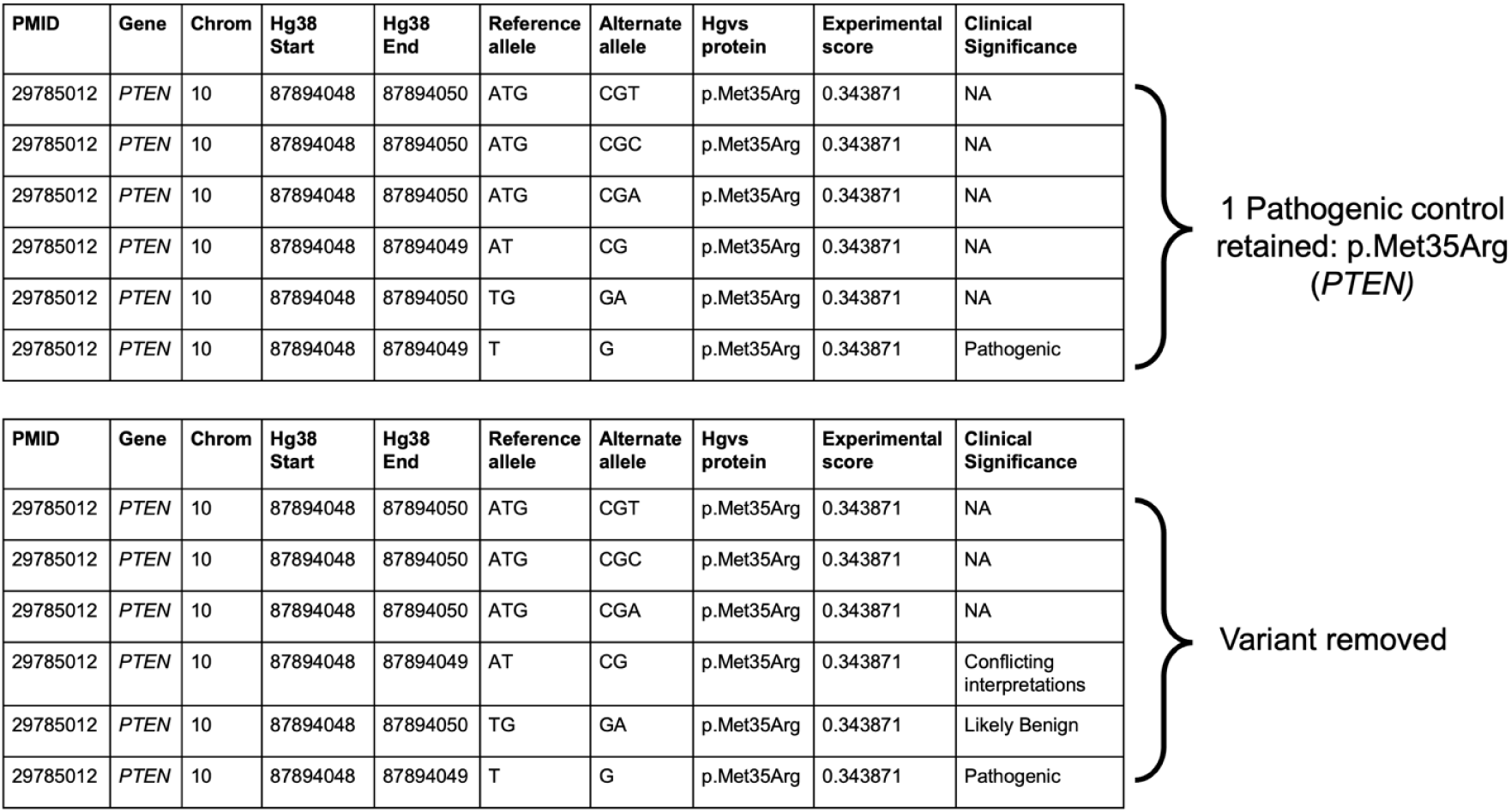

##### ExCALIBR calibrations

We developed ExCALIBR, a statistical method to calibrate high-throughput experimental data^33^. Briefly, we jointly modeled scores of pathogenic, benign, gnomAD, and synonymous variants as skew normal mixtures. For *BRCA1*, *TP53*, *MSH2*, and *PTEN*, ClinVar control classifications from 2018 were used. We performed 1,000 bootstrap iterations of each dataset, using the 5th and 95th percentiles of the local positive likelihood ratio along with the median estimated prior probability of pathogenicity to define pathogenic and benign evidence strengths expressed in ACMG/AMP points with the exception that our approach enables assignment of all possible evidence points (e.g. ±1 to ±8) (**Supplementary Data 4, ExCALIBR calibrations)**. The prior probability of pathogenicity was estimated using the score distribution of gnomAD variants as a population reference^33^. When comparing ExCALIBR calibrations to author-designated functional classes **(Fig. 4c)**, evidence was assigned in an out-of-bag manner; i.e., each variant’s evidence only considered bootstrap models that excluded it during fitting, ensuring unbiased evaluation. Conversely, threshold-based approaches may classify ClinVar controls that were simultaneously used to define the score intervals representing functional classes, potentially inflating performance.

##### Calibration of computational variant effect predictions

We developed a gene-specific calibration workflow that adaptively selects the optimal calibration strategy for each gene based on score distribution characteristics and data availability. Our strategy is divided into four steps.

**Step 1** (gene-specific prior estimation): For each gene, we first estimated the prior probability of pathogenicity using an adapted *DistCurve* algorithm^40,42^ leveraging ClinVar control pathogenic variants together with unlabelled variants from gnomAD. This procedure yields a gene-specific prior probability of pathogenicity that was subsequently used both to generate simulated datasets and to define the posterior probability ranges corresponding to ACMG/AMP evidence points.

**Step 2** (score distribution modeling): For each gene, we independently modeled the pathogenic and benign variant score distributions using four parametric approaches (*Beta*, *Truncated Normal*, *Truncated Skew-t*, and *Truncated Skew-Cauchy*) and selected the best-fitting model from each approach by maximum log-likelihood.

**Step 3** (simulation and method evaluation): Using the fitted distributions and the estimated gene-specific prior, we generated 30 synthetic datasets per gene with known posterior probabilities. For each gene, we evaluate 9 local calibration configurations (varying window size and gnomAD smoothing fraction) as in Pejaver et al.^43^ and 10 alternative post hoc calibration methods^42^, each assessed with 1,000 bootstrap iterations. Calibration thresholds were estimated from variants not used in model training (out-of-bag), and we use the 5th percentile of posterior estimate for each threshold to mitigate overestimation of evidence strength.

**Step 4** (multi-stage model selection): A single calibration model was selected for each gene through a three-stage filtering procedure: (i) an overestimation risk filter that removed calibration models exhibiting clinically unsafe behavior—defined as ≥2 evidence point error at the 75th percentile or ≥3 points maximum in pathogenic-or benign-supporting regions in at least 25% of simulations; (ii) average misestimation ranking, retaining the three calibration models with the lowest mean ACMG-point error across pathogenic, benign, and indeterminate regions; and (iii) an overestimation frequency filter that selected the calibration model with the lowest combined pathogenic-and benign-fraction, reflecting the fewest ≥1-point overestimations in key evidence regions. The selected optimal calibration model was then applied to the target gene to produce final calibrated posterior probabilities and to define predictor score intervals corresponding to each ACMG/AMP evidence strength.

#### Scalable workflow for applying experimental and predictive evidence to variant classification

##### Filtering variants for classification analyses

All analyses described below began with filtering of the integrated variant effect dataset (434,166 variant effect measurements) **Supplementary Data 1**). We excluded all variants in *SFPQ* because it does not have a definitive gene-disease association, a subset of variants in *CHEK2* deemed inappropriate for clinical analysis by the original authors, and all variants flagged as having unreliable or inconsistent variant annotations (**Supplementary Data 1,‘Flag’column)**. Variants receiving conflicting experimental evidence (i.e. assigned both positive and negative evidence points from two or more assays) were removed.

We excluded *TP53* and *F9* variant effect measurements, using OddsPath calibrations on output from classifier models that integrate these data instead^32,61^. We excluded all splice-altering variants defined as having a SpliceAI score greater than or equal to 0.2 for donor (loss or gain) and acceptor (loss or gain) because although some assays are sensitive to splice variants (e.g. SGE), the REVEL, MutPred2 and AlphaMissense are not trained to detect them. We also exclude start-loss variants in non-SGE assays. Applying these filters resulted in 307,558 variant effect measurements. These filters resulted in a set of 239,836 variants across 39 genes for classification analysis.

| Filtering step | Variant effect measurements | $\Delta$ Measurements | Unique variants | $\Delta$ Variants |
| --- | --- | --- | --- | --- |
| Integrated variant effect dataset (**Supplementary Data 1**) | 434,166 |  | 255,354 |  |
| Replace F9 and TP53 datasets with model calibrations; remove SFPQ | 325,151 | -109,015 | 252,755 | -2,599 |
| Exclude *CHEK2* variants considered unsuitable for clinical analysis based on author guidance<sup>105</sup> | 322,446 | -2,705 | 250,050 | -2,705 |
| Remove variants with conflicting experimental evidence | 317,479 | -4,967 | 248,161 | -1,889 |
| Remove predicted splice variants (Splice AI $\geq 0.2$) | 307,778 | -9,701 | 239,967 | -8,194 |
| Remove start-loss variants in non-SGE assays | 307,651 | -127 | 239,907 | -60 |
| Remove variants with inconsistent mapping and annotations (Flag column = *) | 307,558 | -93 | 239,836 | -71 |

In addition to these filters, when evaluating REVEL and MutPred2, we excluded variants in the respective predictor that were used in training (REVEL, 115 variants pre-filtering; MutPred2, 3,703 variants pre-filtering).

##### Summing evidence points to generate classifications

To generate variant classifications, we summed experimental and predictor evidence points. The experimental evidence points were generated by either the ExCALIBR or OddsPath calibrations (**Supplementary Data 4, OddsPath calibrations, ExCALBR calibrations**). The predictive evidence points were generated for REVEL, MutPred2, or AlphaMissense using gene-specific calibrations (**Supplementary Data 4, Gene-specific calibrations**) or genome-wide calibrations from Bergquist et al. for all other genes^42–44^. The summed evidence points yielded a classification in the points-based ACMG/AMP framework^59^ **(Supplementary Data 5, Supplementary Data 6)**. For all downstream analysis, for duplicate variants across assays, we retained the variant classification associated with the largest absolute point value from our scalable workflow.

##### ClinVar control analysis

ClinVar controls were defined as variants with clinical significance of Pathogenic, Likely pathogenic, Pathogenic/Likely pathogenic or Benign, Likely benign, Benign/Likely benign from the ClinVar variant summary file downloaded from <u>January 2025</u> and <u>December 2018</u> releases. ClinVar controls were filtered to retain variants with 1+ stars in ClinVar meaning that some assertion criteria were provided by the submitter. We used controls from ClinVar January 2025 except for *BRCA1*, *TP53*, *MSH2*, and *PTEN* where classifications from 2018 were used to avoid comparison with variants classified with evidence from these datasets. For protein-level assays, at most one ClinVar assertion per amino acid variant for a given gene and transcript pair was retained (OddsPath calibrations). After applying filters mentioned above, we retained 10,228 ClinVar control variants for analysis with REVEL (excluding variants used in REVEL training), 10,250 ClinVar controls for AlphaMissense, and 9,099 ClinVar controls for MutPred2 (excluding variants used in MutPred2 training).

To quantify the performance of our scalable workflow, we assessed the concordance between each variant’s classification from our workflow with its original ClinVar classification. True positives (TP) were defined as variants for which our pathogenic or likely pathogenic (PLP) classifications were concordant with the original ClinVar classification PLP. True negatives (TN) were defined as variants for which our benign or likely benign (BLB) classifications were concordant with the original ClinVar classification of BLB. False positives (FP) were defined as variants classified as PLP by our scalable workflow but classified as BLB in ClinVar. False negatives (FN) were defined as variants classified as BLB by our scalable workflow but classified as PLP in ClinVar. We then calculated the following metrics to assess our framework:

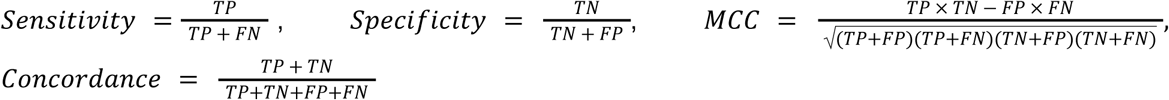

We deemed classifications of BLB and PLP “determinate” (in contrast to VUS). We therefore computed the fraction of determinate classifications by dividing the number of variants classified as PLP and BLB by the total number of variants classified. We deemed classifications “discordant” if our workflow gave a determinate classification that disagreed with ClinVar (e.g. a variant we classified as BLB that was PLP in ClinVar and vice versa). Thus, we computed the fraction of discordant classifications by dividing the number of BLB->PLP and PLP->BLB classifications by the total number of variants analyzed (**Supplementary Data 5, Supplementary Data 6)**.

##### ClinGen Evidence Repository analysis

We retrieved all nucleotide variants from the ClinGen Evidence Repository for our genes which included all evidence codes used to classify each variant (downloaded November 10th, 2025, v 2.5.0). We mapped all possible nucleotide substitutions for amino acid variants from protein-level assays to variants found in the ClinGen Evidence Repository, and found no conflicting classifications. 351 variants had determinate (PLP, BLP) classifications in the Repository and experimental data. For each variant, we removed the experimental evidence (BS3, PS3) and/or predictive evidence (PP3, BP4) used in the original classifications and then recalculated the classification without this evidence^57^. 194 variants across 11 genes retained sufficient evidence to remain BLB (19) or PLP (175). Of these, post filtering, we retained 179 variants for analysis with REVEL (excluding variants used in REVEL training), 182 variants for AlphaMissense, and 35 variants for MutPred2 (excluding variants used in MutPred2 training) We then assessed the concordance of our classifications using only experimental or predictive evidence to the expert panel classifications generated from all other sources of evidence using the same performance metrics as the ClinVar control analysis (**Supplementary Data 5, Supplementary Data 6)**.

##### ClinVar VUS, gnomAD, and unobserved variants analysis

VUS were defined as variants classified as Uncertain significance in the January 2025 ClinVar variant summary file (**Supplementary Data 1**). gnomAD variants were defined as variants observed in gnomAD (i.e., with a reported allele frequency). Unobserved variants were defined as single nucleotide variants absent from both ClinVar in January 2025 and gnomAD v4. Post filtering, we retained 16,115 VUS, 24,992 gnomAD variants, and 90,530 unobserved variants for analysis with REVEL. For analysis with AlphaMissense, we retained 16,149 VUS, 25,063 gnomAD variants, and 90,550 unobserved variants. For MutPred2 we retained 15,705 VUS, 23,918 gnomAD variants and 89,463 unobserved variants.

Variant classifications for ClinVar VUS, gnomAD variants, and unobserved variants were determined using the same scalable framework described above. For each nucleotide variant, experimental and predictive evidence points were summed, and the resulting total points were used to assign a classification according to the framework described by Tavtigian et al^59^. (**Supplementary Data 5, Supplementary Data 6**).

##### Biobank validation of variant classes

To further validate our scalable workflow, we assessed the relationship between variant classes (defined by experimental data, evidence assigned by calibrations of experimental and predictive data, and classifications) and clinical phenotypes in ∼400,000 participants in the All of Us biobank. The validation was performed within the All of Us Workbench using the All of Us Controlled Tier Dataset v8^106^ Curated Data Repository (CDR) version 8 Release Notes – User Support.

##### Gene-specific cohort definition

For each of the 40 clinically-relevant genes we analyzed, we evaluated whether the gene is known to be associated with an autosomal dominant phenotype that can be defined in terms of diagnosis codes (reviewed by a medical genetics physician and a molecular pathologist), measurements, and surveys recorded in All of Us (AoU) and whether there is a sufficient number of participants (at least 40) with whole genome sequencing data in the biobank that fit each definition and have variants in the corresponding gene. We determined that 23 gene-disease pairs did not meet these criteria (**Supplementary Table 2**), yielding 17 gene-disease pairs for analysis. For each of these 17 gene-disease pairs, we established a case cohort by selecting participants that had specific conditions, measurements, or survey answers that indicated they had the phenotype of interest. For each gene-disease pair, we also established a control cohort by excluding participants that had broad conditions, measurements, or survey answers that indicated they might have the phenotype of interest or a closely related phenotype (**Supplementary Table 3**, **Supplementary Data 8**).

##### Definition of carrier status for a set of variants

We aggregated variants into sets based on four classification schemes: (i) experimental data functional classes, (ii) ExCALIBR-calibrated experimental evidence aggregated by assigned points, (iii) calibrated predictive evidence aggregated by assigned points (iv) the sum of experimental and predictive evidence aggregated by assigned evidence points. For each variant set, we queried the All of Us short-read whole genome sequencing matrix to identify carriers, defined as participants with one or more variants in that set.

##### Odds ratio estimation

For every set of variants considered (e.g. functional class, variants with ≥2 points of experimental evidence), we estimate the odds ratio of disease phenotype given carrier status by fitting a logistic regression model to the cohort of cases and controls established for the corresponding gene as described above. When the variant set is defined by computational predictors or assays that do not measure the effect of splicing, we additionally exclude all participants with any variants in the gene that have a SpliceAI score above 0.2 for donor or acceptor gain or loss. The model response variable is case or control status, and explanatory variables are carrier status for the variant set, sex assigned at birth (trichotomized into male, female, and other), age at the time of AoU data release in years, and 16 genetic ancestry PCA components computed by the AoU research program^107^. We report the point estimate, 95% confidence interval, and Wald test p-value for the regression coefficient associated with the carrier status variable, which is the natural log of the odds ratio of interest (**Supplementary Data 2**, **Supplementary Data 8**).

## Data Availability

Raw and analyzed SGE and VAMP-seq data, calibrations, arrayed assay data, and **Supplementary Data 1** are available from the IGVF Portal, see **Supplementary Table 1** for accession numbers.

SGE and VAMP-seq results are available on MaveDB and MaveMD see **Supplementary Table 1** for MaveDB urns.

All supplementary data files and large supporting files for analysis and figure generation are also hosted at https://zenodo.org/records/18637474.

## Code Availability

Code supporting this study is publicly available at the following repositories:

SGE analysis pipeline: https://github.com/bbi-lab/sge-pipeline/

VAMP-seq analysis pipeline (CountESS): https://github.com/CountESS-Project/CountESS Pipelines for arrayed assays: https://github.com/broadinstitute/2025_Pillar_VarChAMP Biobank analyses https://github.com/Craven-Biostat-Lab/igvf-cvfg-biobank

The integrated variant effect datasets pipeline and variant classification analysis are available at: https://github.com/bbi-lab/IGVF-cvfg-pillar-project, including scripts to generate main and extended data figures.

## Supplementary Notes

### Supplementary Note 1

#### PTEN Variant Misclassification

One misclassified variant was *PTEN* (NM_000314.6:c.500C>A), a *de novo* variant associated with autism spectrum disorder. Experimental evidence supported a benign classification (−2 points) while predictive evidence supported pathogenicity (+1 point), resulting in −1 combined points and an LB classification. This discordance suggests that the experimental assay measuring lipid phosphatase activity^108^ may not fully capture the ASD-relevant mechanism for PTEN. Specifically, NM_000314.8(PTEN):c.500C>A (p.Thr167Asn) was identified *de novo* in a 99-month-old white female with autism spectrum disorder from the Simons Simplex Collection (SSC; proband 11390.p1)^109^. The individual presented with macrocephaly, speech delay with regression, social communication deficits, ADHD, pica, sleep disturbances, recurrent otitis media, and gastrointestinal issues. Family history is notable for autism features in the mother, ADHD in a sibling, and an extensive family history of neuropsychiatric conditions. Both parents and the sibling did not have macrocephaly. While functional evidence does not show that this variant is low abundance or lacking lipid phosphatase activity, we have recently learned PTEN variants associated with neurodevelopmental disorders may disrupt PTEN function through other mechanisms. A recent study which used visual phenotype to assess variant effects on PTEN subcellular localization demonstrated that this variant is nuclearly mislocalized, similar to other PTEN variants known to be associated with neurodevelopmental disorders^91^.

### Supplementary Note 2

#### Minimum Sensitivity for BS3_supporting

The likelihood ratio for a normal assay result (LR−) equals (1 − sensitivity) divided by specificity. Setting LR− equal to the BS3_supporting threshold of 0.48 and solving for sensitivity at 90% specificity yields a minimum sensitivity of 56.8%, calculated as 1 − (0.48 × 0.90). An assay with 90% specificity can therefore have sensitivity as low as 57% and still qualify for BS3_supporting, meaning such an assay fails to detect 43% of pathogenic variants.

#### False Omission Rate at Different Priors

The false omission rate (FOR) equals the product of (1 − sensitivity) and the prior probability, divided by the sum of that product and the product of specificity and (1 − prior). At 57% sensitivity and 90% specificity, the FOR is 4.5% when the prior probability of pathogenicity is 10%, rises to 13.7% at a 25% prior, and reaches 32.3% at a 50% prior. When the prior exceeds the ACMG/AMP assumption of 10%, the proportion of misclassified pathogenic variants increases substantially. At a 50% prior, nearly one in three likely benign classifications based on single-source weak evidence would be incorrect.

## Acknowledgements

This work was primarily supported by the following National Institute of Health (NIH) National Human Genome Research Institute (NHGRI) Impact of Genomic Variation on Function (IGVF) consortium center awards: the Center for Actionable Variant Analysis (CAVA) to D.M.F. and L.M.S. (UM1HG011969), the Radivojac Center to P.R. (U01HG012022), the Craven Center to M.C. (U01HG012039), the Variant Characterization Across the Mendelian Proteome (VarChAMP) Center to M.V. (UM1HG011989) and the Data Analysis and Coordinating Center (DACC) to B.C.H. (U24HG012012). Additional financial support was provided by an NHGRI Advancing Genome Medicine Research consortium award (R01HG013025, A.E.M., A.F.R., D.M.F., A.B.S. and L.M.S.), an NHGRI GREGoR consortium award (U01HG011744, support to L.M.S., S.C. and S.B.), NHGRI grants (R01HG013350, V.P.; P50HG004233, M.V.), NIH National Institute of Mental Health (NIMH) grants (R56MH128365, L.M.I.; R21MH128827, L.M.I.), an NIH National Heart, Lungs and Blood Institute (NHLBI) grant (R01HL152066, J.M.J. and D.M.F.), the NIH Intramural Research Program, a Clinical and Translational Science Awards (CTSA) grant from the National Center for Advancing Translational Sciences (UL1TR004419), awards from the NIH Office of Research Infrastructure (S10OD026880; S10OD030463), the generous support of the Brotman and Baty families through the Brotman Baty Institute for Precision Medicine, a Chan Zuckerberg Initiative Foundation grant (CZIF2024-010284, D.M.F. and L.M.S.), a Simons Foundation Autism Research Initiative (SFARI) grant (896503, L.M.I. and M.V.), the Australian Government (A.F.R.), a RUNX1 Foundation/Alex’s Lemonade Stand for Childhood Cancer Early Career award (21-25037, A.E.M.), a Brotman Baty Institute Catalytic Collaborations grant (CC28, A.E.M.), a Medical Genetics Postdoctoral Fellowship (T32GM007454, S.F.), Belgian American Educational Foundation (BAEF) Doctoral Research Fellowships (F.L., M.T.), Wallonia-Brussels International (WBI)-World Excellence Fellowships (F.L., G.C.), a Fonds de la Recherche Scientifique (FRS-FNRS)-Télévie grant (FC31747–7459421F, F.L.), a Herman-van Beneden Prize (F.L.), a Josée and Jean Schmets Prize (F.L.), the Léon Fredericq Foundation (F.L.), a FRS-FNRS-Fund for Research Training in Industry and Agriculture (FRIA) grant (FC31543–1E00419F, G.C.), a University of Toronto Open Fellowship (C.R.), a Cecil Yip Doctoral Research Award (C.R.) and Dana-Farber Cancer Institute Center for Cancer Systems Biology (CCSB) Deborah F. Allinger Fellowships (A.Y., L.L.). M.V. is a Chercheur Qualifié Honoraire from FRS-FNRS (Wallonia-Brussels Federation, Belgium). We gratefully acknowledge All of Us participants for their contributions and thank the NIH All of Us Research Program for making available the participant data. We acknowledge support from the Minerva computational and data resources and staff expertise provided by the Scientific Computing and Data group at the Icahn School of Medicine at Mount Sinai, and from the Explorer Cluster, supported by Northeastern University’s Research Computing team. Support for title page creation and format was provided by AuthorArranger, a tool developed at the NIH National Cancer Institute (NCI). The contributions of the NIH authors are considered Works of the United States Government. The findings and conclusions presented in this document are those of the authors and do not necessarily reflect the views of the NIH or the U.S. Department of Health and Human Services. We thank past and current members of our groups for helpful discussions and guidance.

## Author Contributions

This project was conceptualized by M.Te., Y.C., A.E.M., R.St., Y.S., F.L., D.Z., S.F., J.S., M.V., M.Ta., A.E.C., M.A.C., M.C., V.P., A.F.R., P.R., D.M.F. and L.M.S. The original manuscript was drafted by M.Te., Y.C., A.E.M., R.St., Y.S., F.L., I.W., R.Sh., S.F., P.R., D.M.F. and L.M.S. The manuscript was reviewed by all authors and edited by M.Te., Y.C., A.E.M., R.St., Y.S., F.L., I.W., R.Sh., S.F., P.G., M.B., S.P., J.M., L.M.I., S.S., B.A.C., M.Ta., A.E.C., M.A.C., V.P., A.F.R., P.R., D.M.F. and L.M.S. MAVE data was generated by S.C., Z.R.W. and M.W.S., with contributions from I.W., S.B., L.R.S., O.S., A.J.V., C.W., M.K.W., S.P., D.H., A.X., A.H., under the supervision of D.M.F. and L.M.S. Experimental data from the community was curated and integrated by M.Te., P.G., and A.E.M., with feedback from D.M.F. and L.M.S. Experimental data calibration was conducted by R.St. and D.Z., with contributions from S.J., and feedback from V.P. and P.R. Predictive data calibration was led by Y.C. and S.F., with contributions from M.B. and S.J., and feedback from V.P., P.R., D.M.F. and L.M.S. Biobank analyses were performed by Y.S., with feedback from R.D.S. and M.C. Clinical reclassification of VUS and unobserved variants was conducted by M.Te., A.E.M., Y.C., R.St., P.G., S.F., M.B., under the supervision of V.P., P.R., D.M.F. and L.M.S. Protein abundance data was generated by Z.Y. and F.L, with feedback from M.A.C. Interactomic data was generated by F.L., G.C. and A.Y., with help from K.S.F., and supervision from M.A.C. Imaging data was generated by C.R. and T.T., with help from F.L., and feedback from M.Ta. Sequence confirmation of DNA clones used in arrayed phenotypic assays was performed by A.G.C., M.G., W.v.L. and K.M.F., under the supervision of F.P.R. Analysis of the protein abundance dataset was led by R.Sh., with assistance from M.Ti., J.D.E., F.L. and G.C., and feedback from M.A.C. and A.E.C. Analysis of the interactomic dataset was led by M.Ti., with assistance from G.C., J.D.E. and F.L., and feedback from A.E.C., L.L. and M.A.C. Image processing and analysis pipelines were developed by Z.S.C., M.H., A.K.H. and J.D.E., and the imaging dataset was analyzed by R.Sh., E.A.M. and E.W., with feedback from S.S., M.Ta., B.A.C. and A.E.C. The specific analysis on factor IX was conducted by R.Sh., with feedback from A.E.C., J.M.J. and D.M.F. MaveDB development was led by B.J.C. and D.R. MaveMD was developed by A.E.M., J.S., E.Y.D. and S.G. PredictMD was built by J.S. The CVFG website was designed and launched by J.S.; development of these four platforms received feedback from D.M.F. and L.M.S. The Data Analysis and Coordinating Center was managed by B.C.H., I.G., K.L. and S.D. Data submission to DACC was handled by N.S. and T.H. Figures were assembled by M.Te., Y.C., R.St., Y.S., F.L., I.W., R.Sh. and C.R., with feedback from M.A.C., A.F.R., P.R., D.M.F. and L.M.S. The multi-institutional IGVF Coding Variant Focus Group was supervised by L.M.I., S.S., B.A.C., F.P.R., R.G.J., M.V., M.Ta., A.E.C., M.A.C., M.C., V.P., A.F.R., P.R., D.M.F. and L.M.S., and overall project administration was handled by J.S., S.H. and L.M. Funding was acquired by J.M.J., A.B.S., L.M.I., F.P.R., M.V., M.Ta., A.E.C., M.A.C., M.C., A.F.R., P.R., D.M.F. and L.M.S.

## Extended Data Figures

**Extended Data Figure 1:**
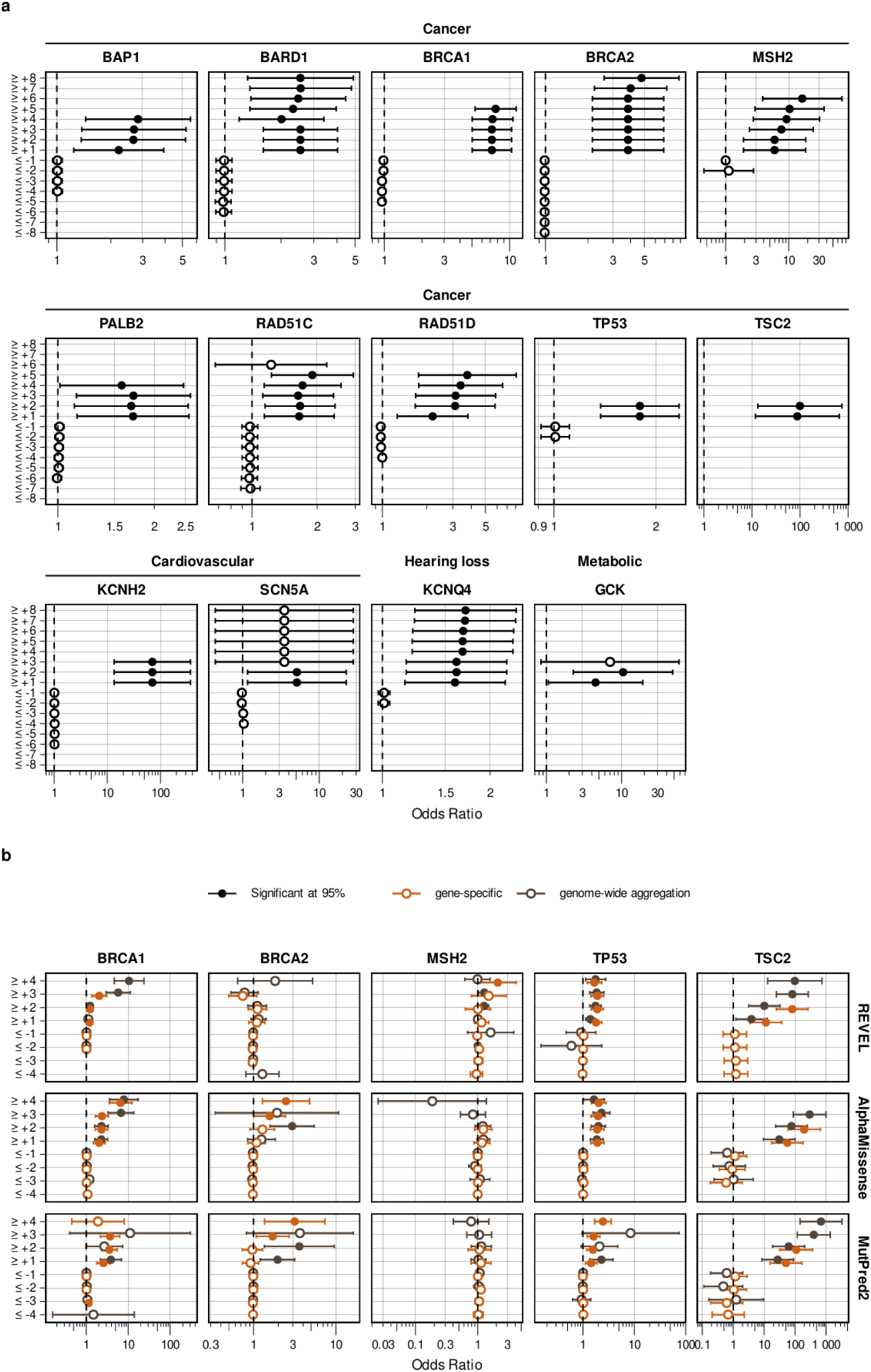
ExCALIBR model fits and assigned evidence for experimental datasets **(See Extended Data Figures 1 and 3 pdf)** ExCALIBR fits, variant score distributions, and calibrated evidence strengths are shown for 61 experimental datasets spanning 37 genes across 11 pages, with each dataset visualized in a single panel containing three components: top, ExCALIBR fits for pathogenic variants (red), benign variants (blue), gnomAD population variants (grey), and synonymous variants (green; exclusive with other variant classes when available), where ClinVar variants are from the December 2018 release for *BRCA1*, *MSH2*, *TP53*, and *PTEN*, and the January 2025 release for all other genes; middle, score distributions for pathogenic (red) and benign (blue) control variants from the January 2025 ClinVar release, overlaid with the distribution of all possible SNVs (light grey); bottom, score intervals corresponding to evidence strength assignments (up to 8 pathogenic or benign points) determined by ExCALIBR, with the number of SNVs assigned to each interval labeled within sufficiently large bins.

**Extended Data Figure 2:** Biobank validation of calibrated evidence. Odds ratios for occurrence of disease in individuals with variants meeting each evidence strength threshold in the All of Us biobank for genes that were found to have significant association with disease at some level of pathogenic evidence. X-axis denotes the odds ratio with the black vertical dashed line indicating an odds ratio of 1. Y-axis denotes the variant grouping by assigned points. Circles denote the estimated odds ratio and whiskers denote the 95% confidence interval. Filled circles indicate that the odds ratio is significant at 95% according to a two-sided Wald test. (a) Odds ratios for variant groups based on ExCALIBR-calibrated experimental evidence aggregated by assigned points for one representative functional study per gene (**Supplementary Data 2**), except for TP53, for which points are assigned based the OddsPath calibration from the (Fayer et al. 2021) meta-analysis^61^. (b) Calibrated predictive evidence aggregated by assigned points, comparing gene-specific calibration (orange) to genome-wide aggregation calibration (brown).

**Extended Data Figure 3:** Gene-specific predictor calibrations across 8 genes for REVEL, MutPred2, and AlphaMissense **(See Extended Data Figures 1 and 3 pdf)** Gene-specific calibrations for REVEL, MutPred2 (MP2), and AlphaMissense (AM), respectively, across 8 genes, each visualized with three components: top, pathogenic (red) and benign (blue) control missense variants from the ClinVar January 2025 release overlaid with all possible missense SNVs (light grey); middle, genome-wide aggregation score thresholds; bottom, gene-specific calibration score intervals, with both middle and bottom panels labeled by evidence strength (up to 4 pathogenic or benign points) and SNV counts shown within sufficiently large bins. Calibrations of MutPred2 scores for *F9* are not shown because after excluding variants used in MutPred2 training, the remaining control variants were insufficient to support a robust gene-specific calibration.

**Extended Data Figure 4:**
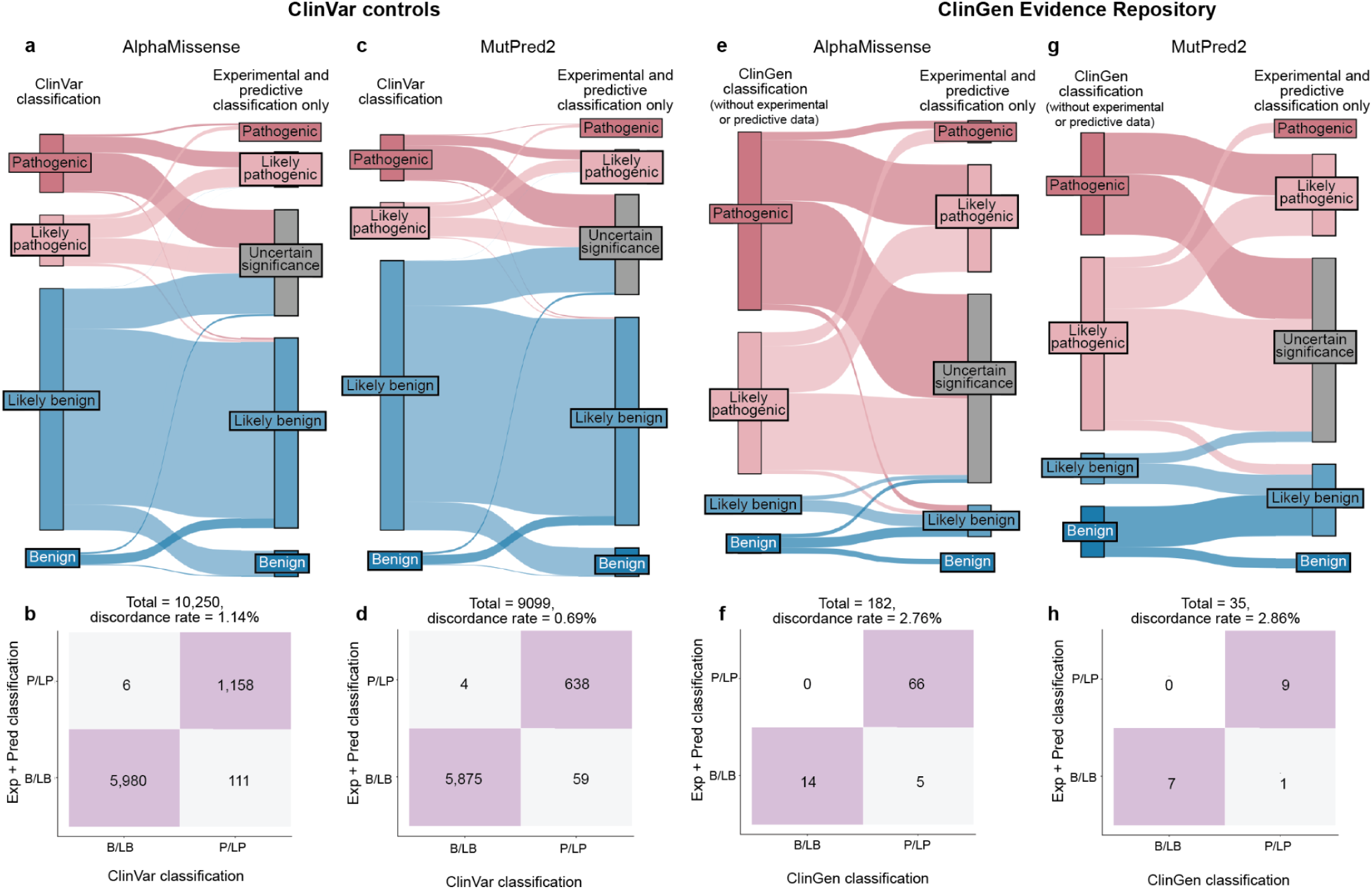
Calibrated experimental and predictive data are highly accurate for AlphaMissense and MutPred2 with a 0.69-1.14% discordance rate (a,c) Sankey diagram depicting the relationship of the variant classifications in ClinVar and their classifications produced by our scalable workflow using only experimental and predictive evidence for (a) AlphaMissense and (c) MutPred2.(b,d) Confusion matrix comparing original ClinVar classifications to classifications from our workflow, summarizing concordant (lilac) and discordant (grey) variant classifications for (b) AlphaMissense and (d) MutPred2. (e,g) Sankey diagram depicting the relationship of the variant classifications from the ClinGen Evidence Repository (v.2.5.0) with updated classifications with experimental or predictive evidence removed and their classifications produced by our scalable workflow using only experimental and predictive (e) AlphaMissense, (g)MutPred2 evidence. (f,h) Confusion matrix comparing updated ClinGen Evidence Repository classifications to classifications assigned using experimental and predictive evidence only, summarizing concordant (lilac) and discordant (grey) variant classifications for (f) AlphaMissense and (h) MutPred2.

**Extended Data Figure 5:**
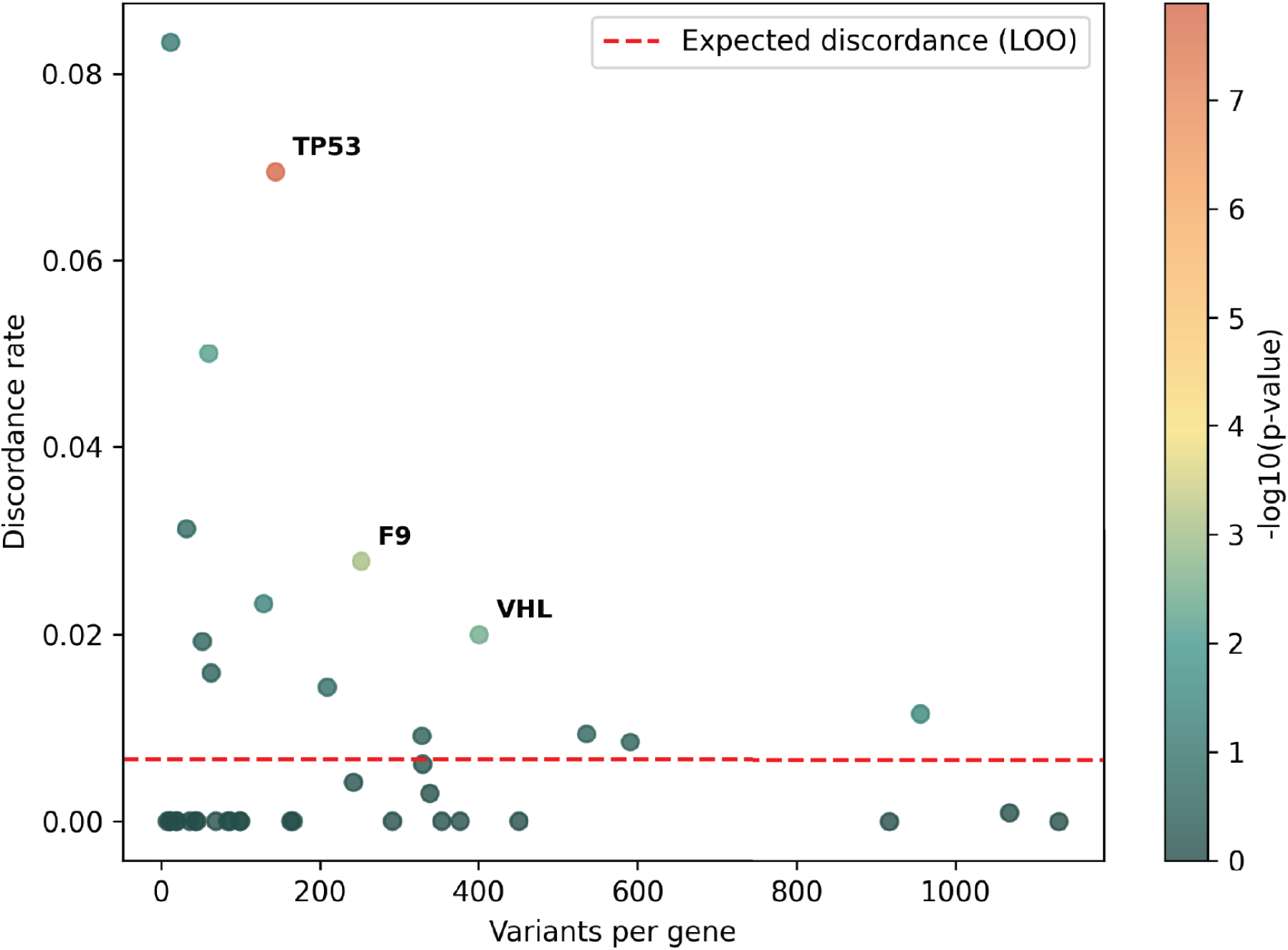
Gene-level enrichment of discordant variant classifications. Genes are plotted by variant count (x-axis) and discordance rate (y-axis) between ClinVar classifications and classifications using only experimental and predictive data (REVEL predictor). Discordance reflects pathogenic-benign classification flips when comparing ClinVar classifications and calculated classifications. The expected discordance rate (red dashed line) was estimated using a leave-one-gene-out background discordance. Colors indicate −log10(p-value) from a binomial test of excess discordance; significantly enriched genes (FDR q < 0.05, ≥10 variants) are labeled.

**Extended Data Figure 6:**
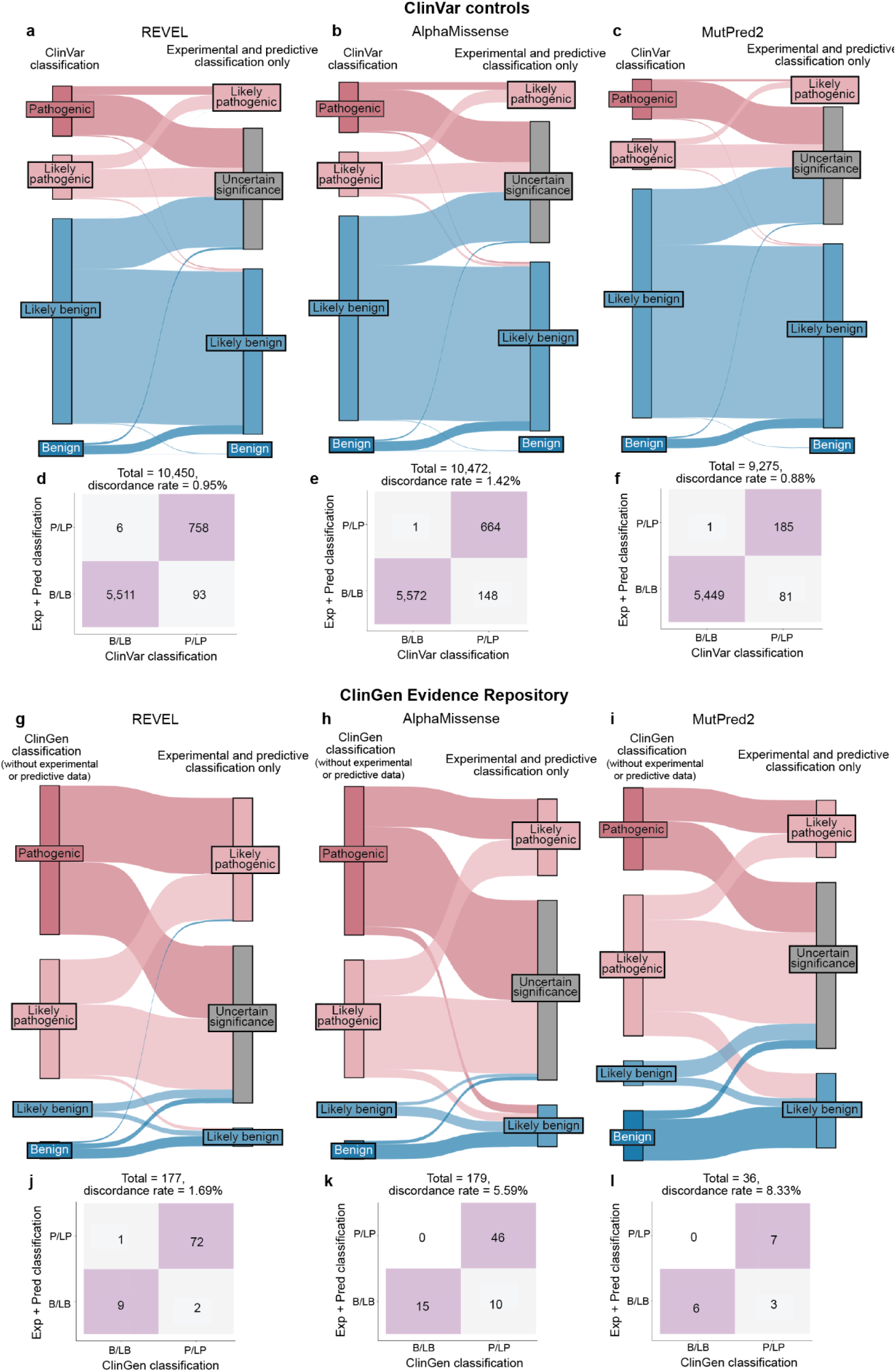
Experimental evidence calibrated using OddsPath shows similar, though lower, accuracy when applied to ClinVar and ClinGen controls (a–c) Sankey diagrams showing the relationship of ClinVar variant classifications and their classifications produced by combining only experimental evidence from OddsPath calibrations^25^ and genome-wide predictive evidence^44^ from (a) REVEL, (b) AlphaMissense, (c) and MutPred2. (d–f) Confusion matrices comparing original ClinVar classifications to classifications assigned using experimental and predictive evidence only (OddsPath and genome-wide predictive evidence)summarizing concordant (lilac) and discordant (grey) variant classifications for (d) REVEL, (e) AlphaMissense, and (f) MutPred2. (g–i) Sankey diagrams showing the relationship of variants from the ClinGen Evidence Repository (v2.5.0), using updated classifications in which experimental and predictive evidence were removed, to classifications assigned using experimental and predictive evidence only (OddsPath and genome-wide predictive evidence), using (g) REVEL, (h) AlphaMissense, and (i) MutPred2. (j–l) Confusion matrices comparing updated ClinGen Evidence Repository classifications to classifications assigned using experimental and predictive evidence only (OddsPath and genome-wide predictive evidence), summarizing concordant (lilac) and discordant (grey) variant classifications for (j) REVEL, (k)AlphaMissense, and (l) MutPred2.

**Extended Data Figure 7:**
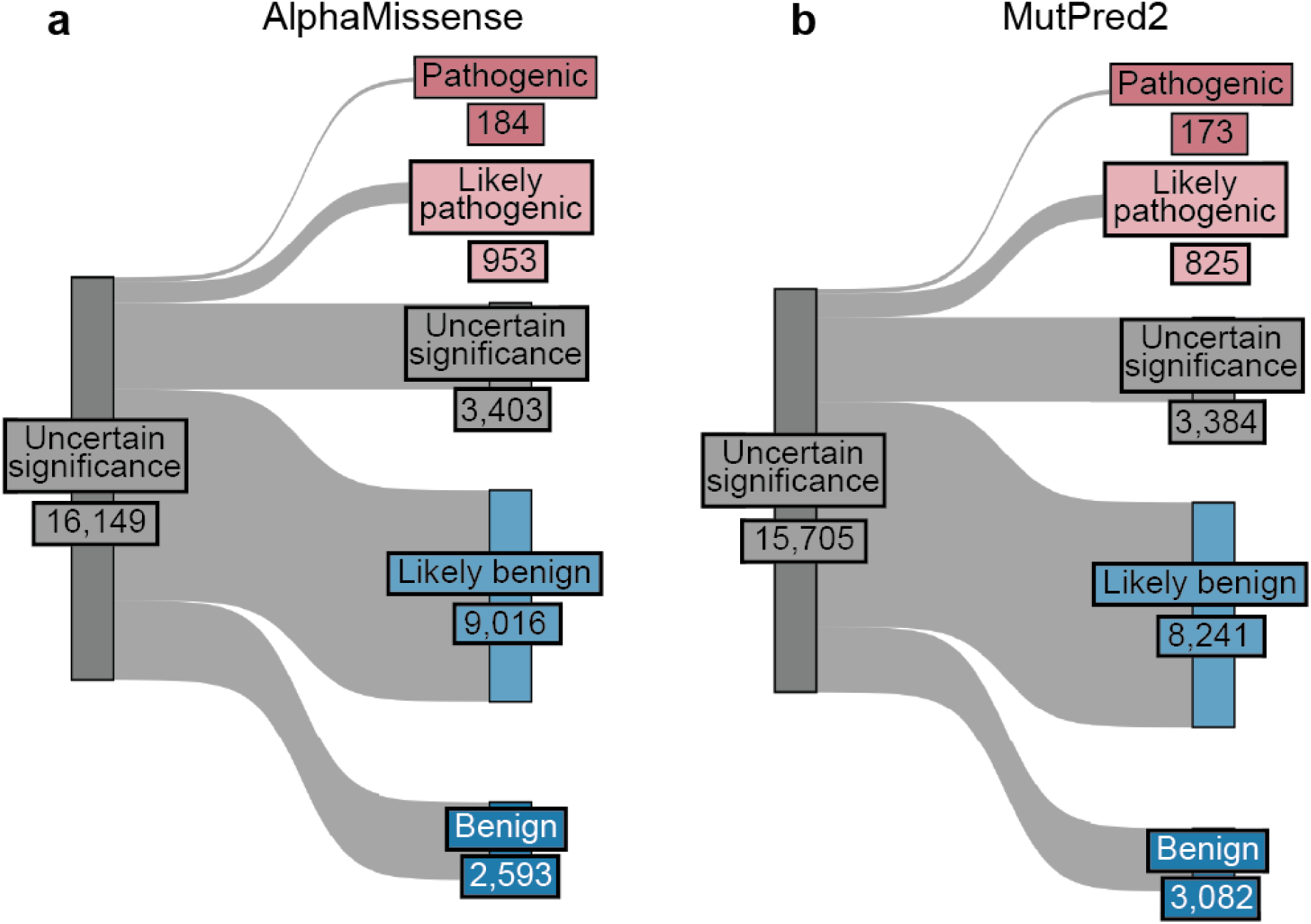
Calibrated experimental and predictive data resolve ∼78% of VUS when using AlphaMissense and MutPred2 (a) Sankey diagram showing the relationship of variants of uncertain significance (VUS) from uncertainty to classifications produced using our scalable workflow using experimental and predictive evidence only for (a) (AlphaMissense) and (b) MutPred2

**Extended Data Figure 8:**
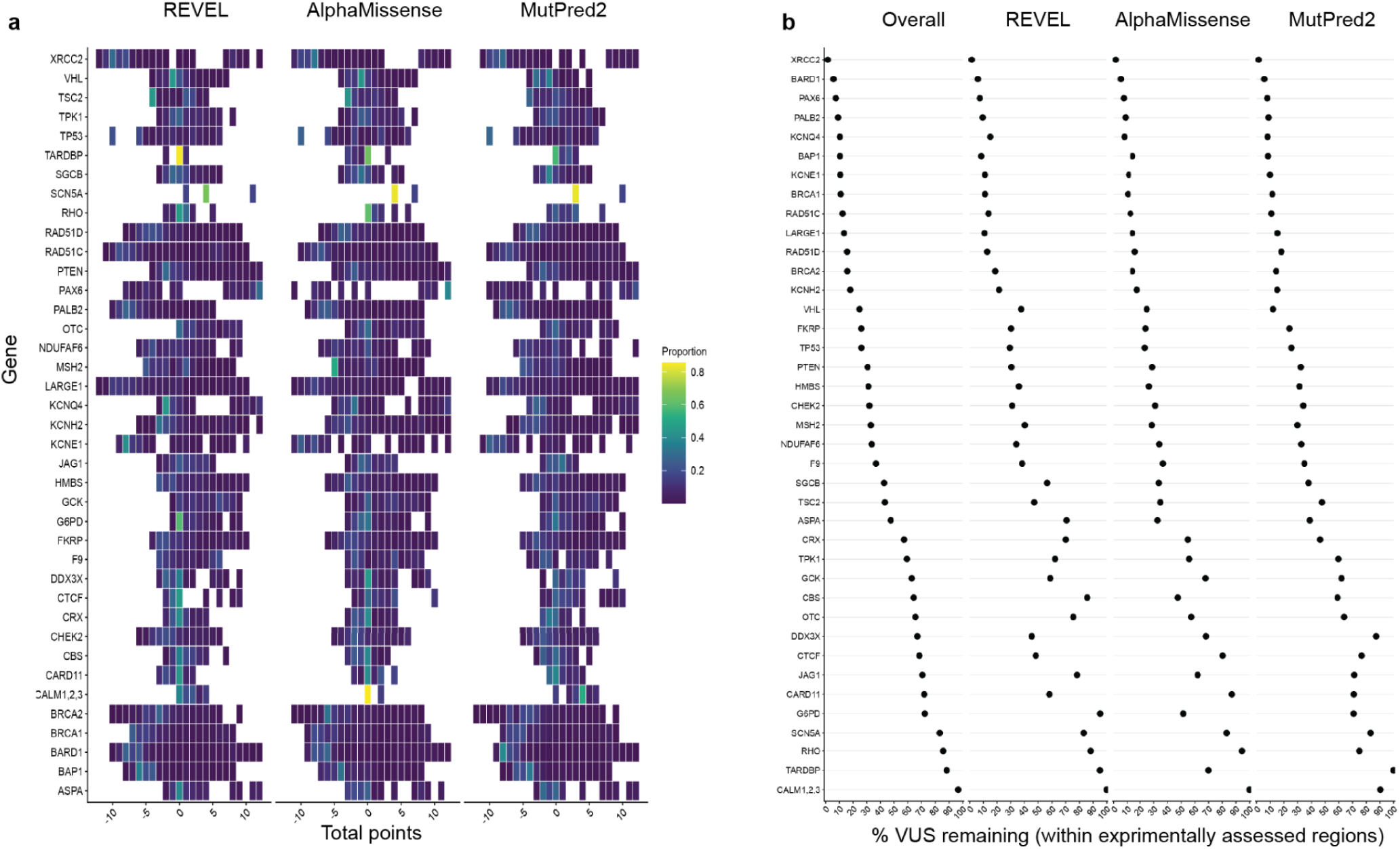
Predictor-specific distribution of total points and remaining VUS burden across genes (a) Heatmaps showing the distribution of total combined evidence points (experimental and predictive evidence) across variants for each gene, stratified by predictor (REVEL, AlphaMissense, MutPred2). For each gene, tile color denotes the proportion of variants assigned to a given total point value, normalized within the gene. Darker shading (towards blue) indicates a lower fraction of variants occupying that point range, highlighting gene-and predictor-specific differences in how variants are distributed across total combined points (b) Rank-based dot plots showing the percentage of variants remaining uncertain (0–5 total points) for each gene (within experimentally assayed regions). Genes are ordered by the overall VUS burden (computed across predictors), and panels display the corresponding value for each individual predictor (REVEL, AlphaMissense, MutPred2)

**Extended Data Figure 9:**
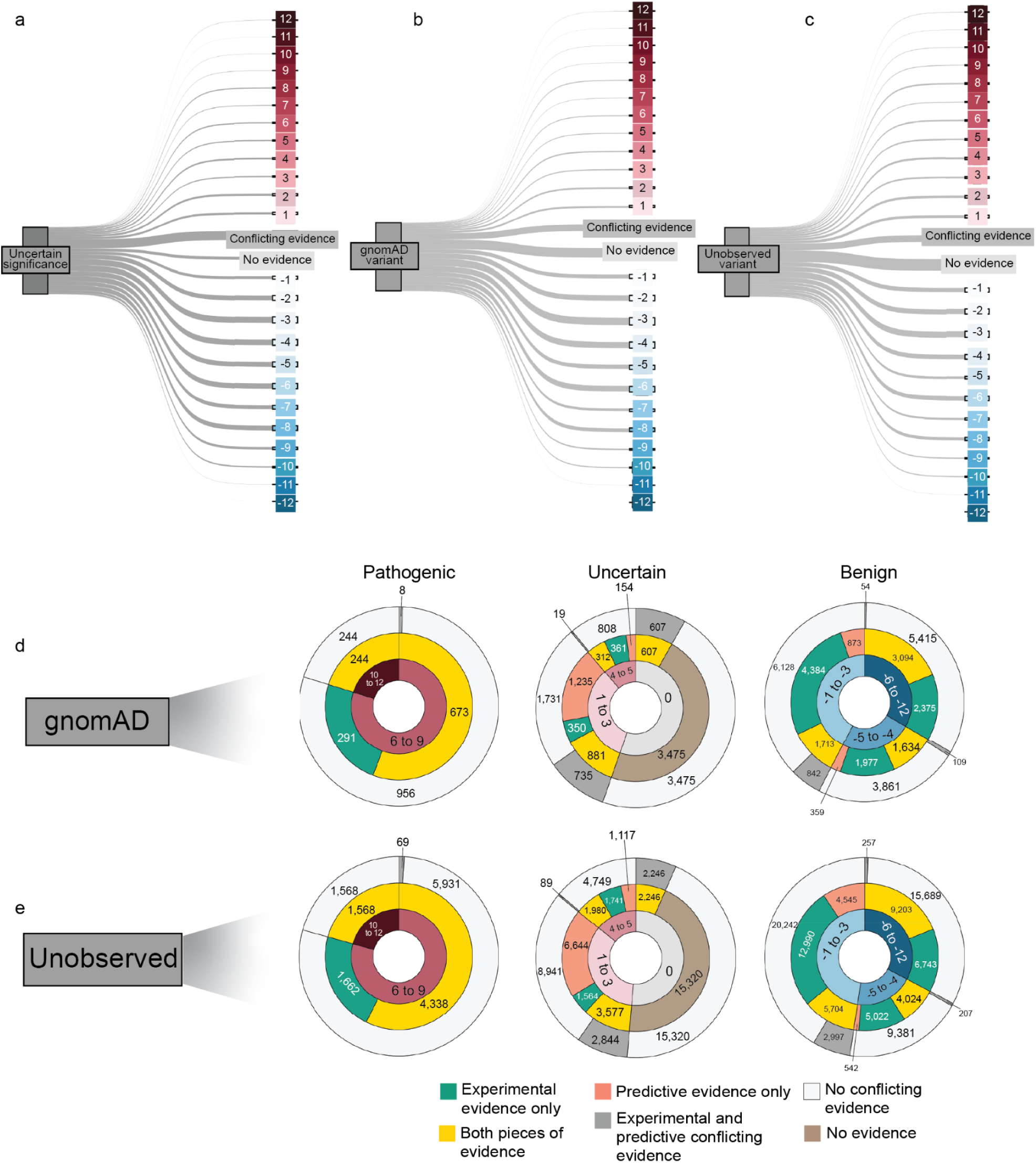
Distribution of variant evidence points across ClinVar VUS, gnomAD and unobserved variants (a–c) Sankey diagrams illustrate the distribution of variants across total evidence point categories (experimental and predictive evidence) for (a) ClinVar variants of uncertain significance (VUS), (b) gnomAD variants, and (c) unobserved variants. The width of each flow is proportional to the number of variants assigned to the corresponding evidence point category. (d–e) Multi-ring donut plots summarizing the evidence source and points for (d) gnomAD variants and (e) unobserved variants. Variants are grouped into pathogenic, uncertain, and benign categories. The innermost ring represents total evidence point ranges, the middle ring denotes the source of evidence (experimental, predictive, or both), and the outermost ring indicates concordance or conflict between evidence sources.

**Extended Data Figure 10:**
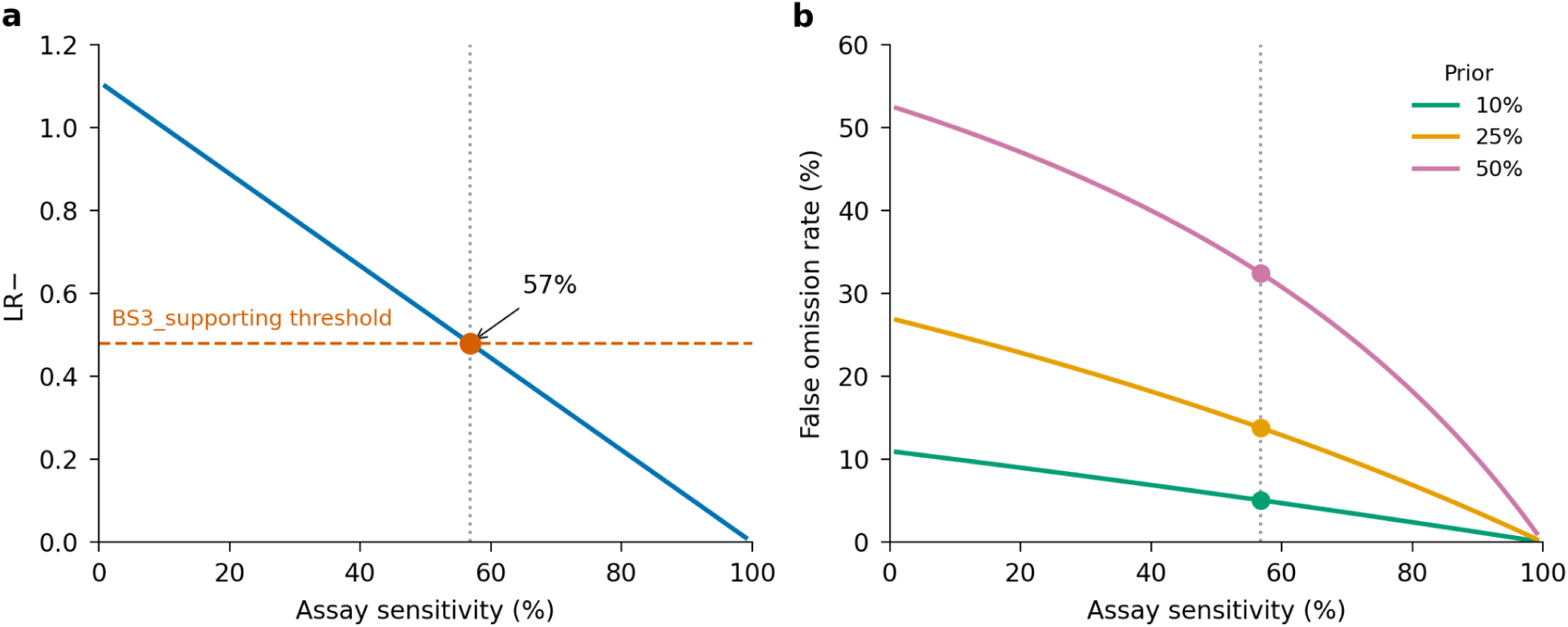
Minimum assay sensitivity for BS3_supporting evidence and resulting false omission rates. (a) Relationship between assay sensitivity and the negative likelihood ratio (LR−) at 90% specificity. The dashed line indicates the LR− threshold (0.48) required for BS3_supporting (−1 evidence point). At this threshold, assays with sensitivity as low as 57% qualify for weak benign evidence. (b) False omission rate as a function of assay sensitivity for different prior probabilities of pathogenicity. At the minimum qualifying sensitivity (57%, dotted line), the false omission rate ranges from 4.5% at the standard ACMG/AMP prior of 10% to 32.3% when the prior is 50%. These results demonstrate that single-source weak benign evidence may lead to misclassification of pathogenic variants, particularly when assay sensitivity is low or when the prior probability of pathogenicity exceeds ACMG/AMP assumptions.

## Supplementary Tables

**Supplementary Table 1:**
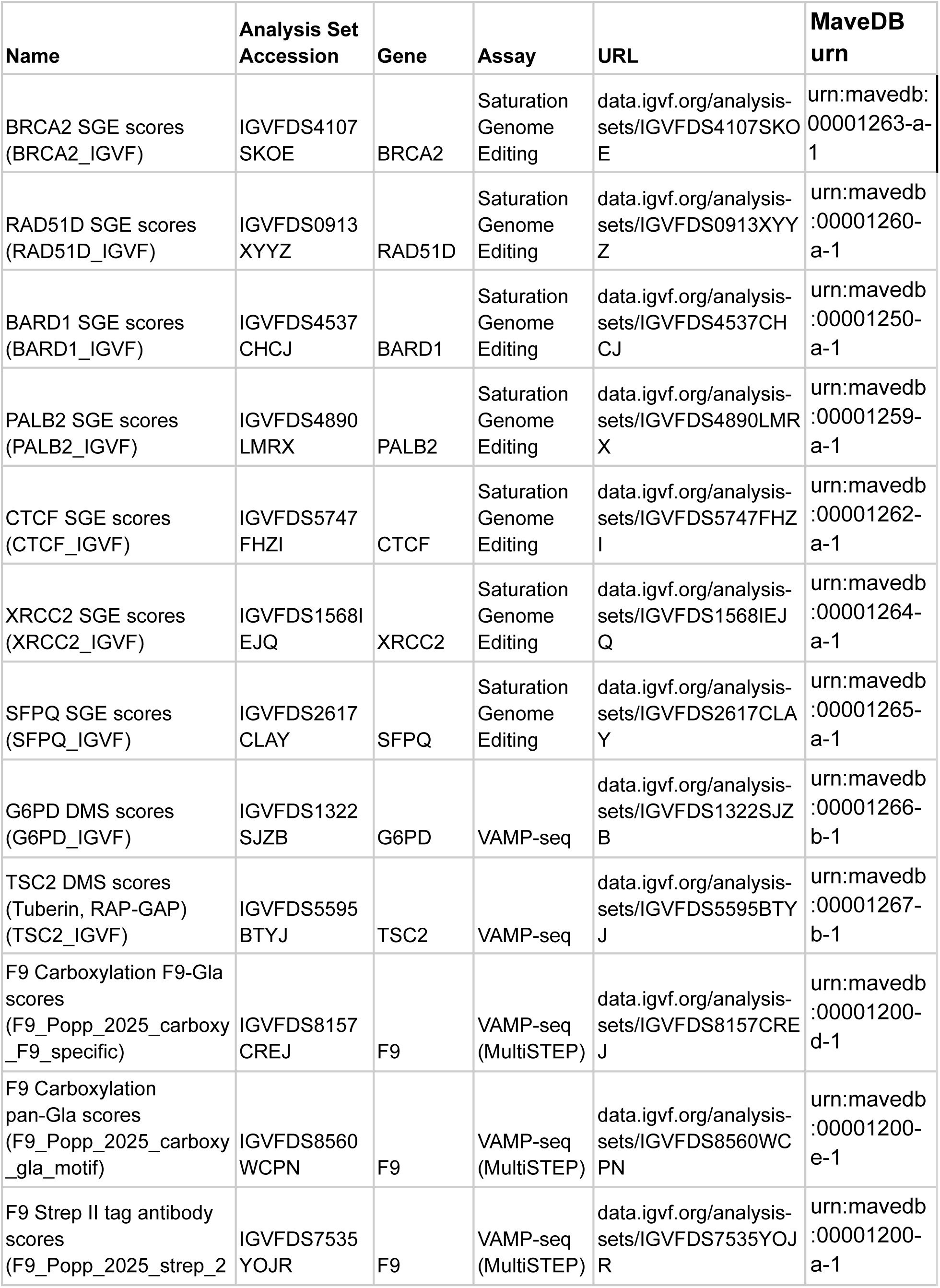
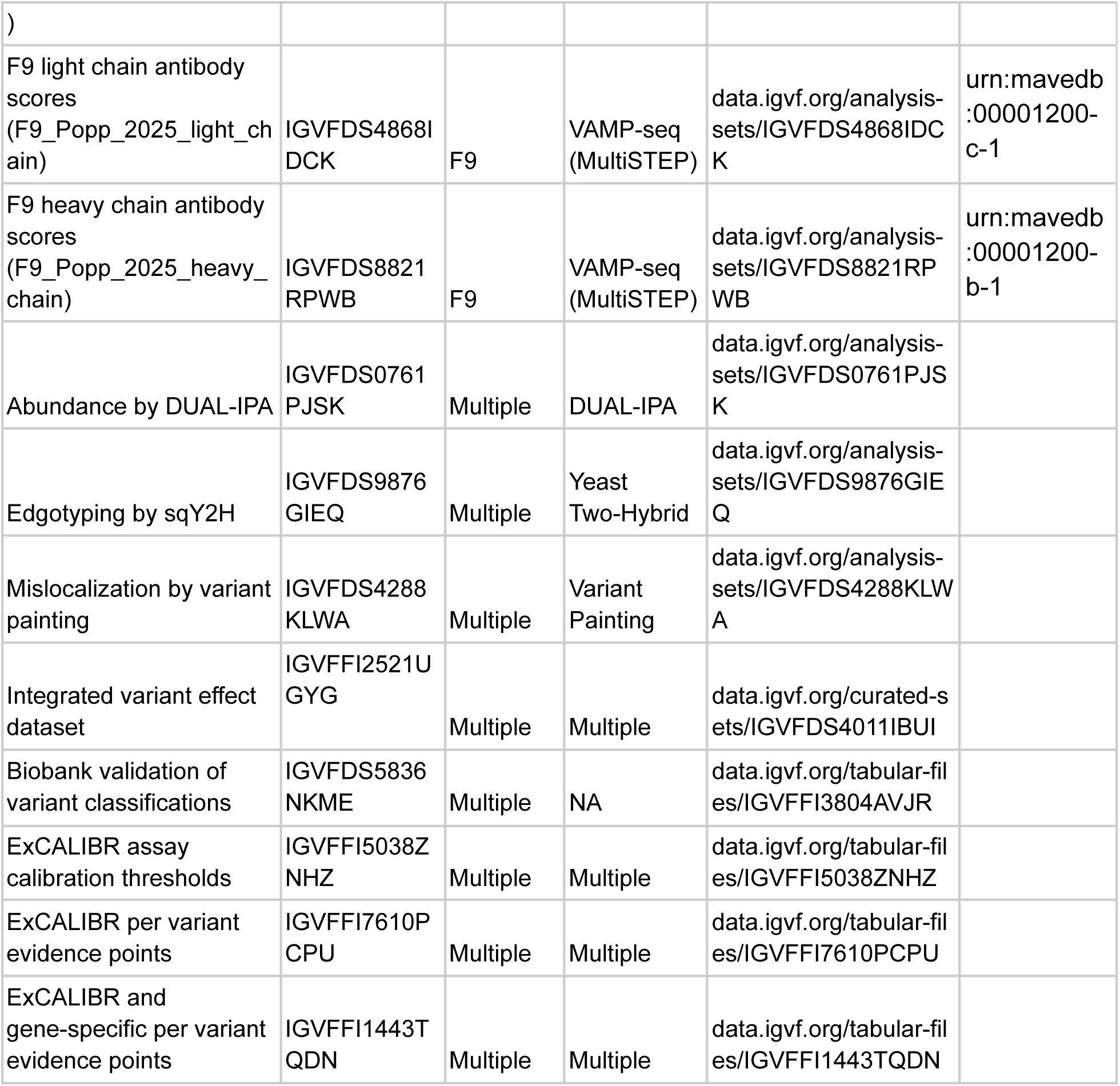
Table listing IGVF accession numbers and corresponding MaveDB URNs for access to raw variant effect measurements and functional calibration data used in this analysis.

**Supplementary Table 2:**
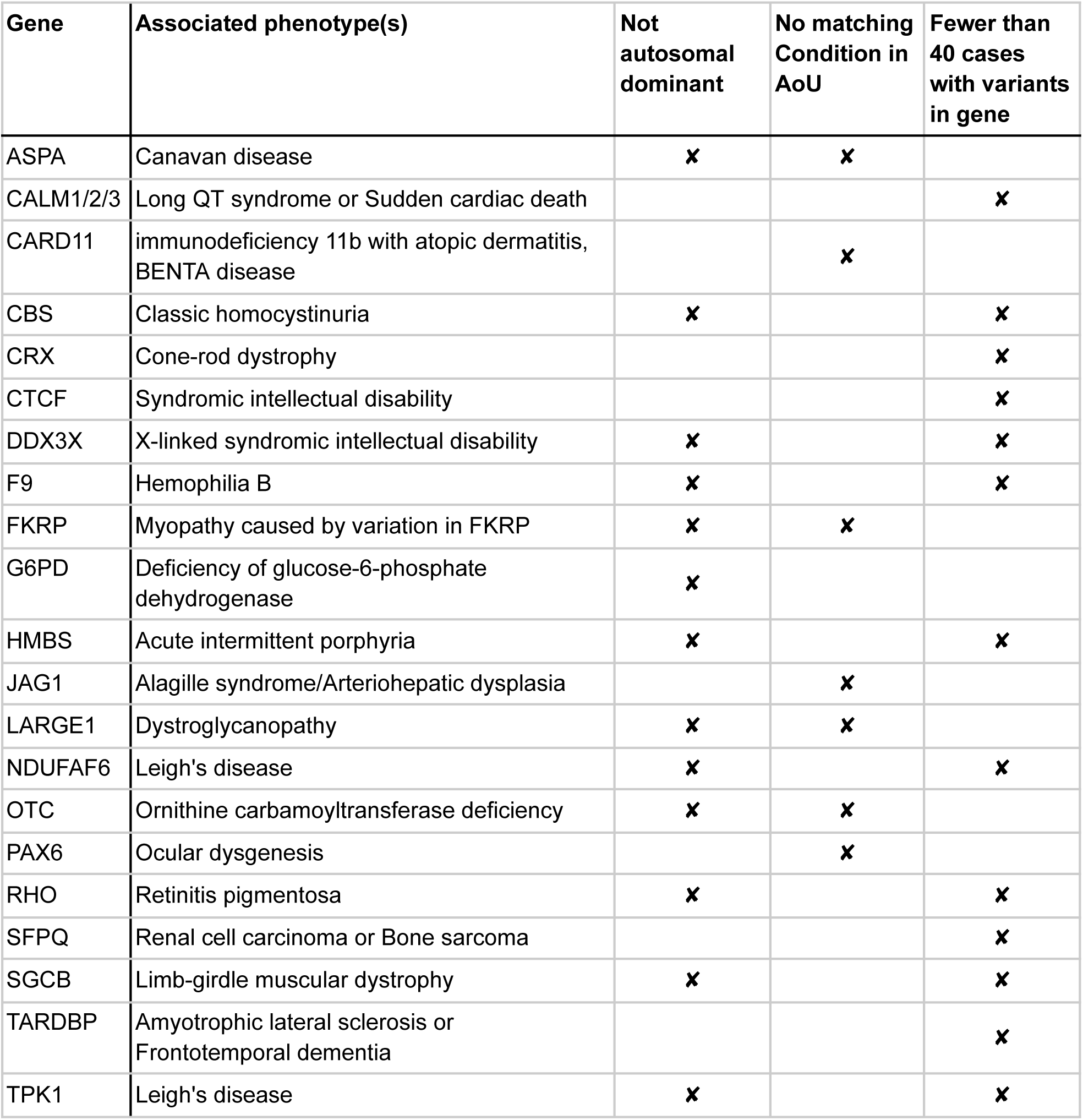
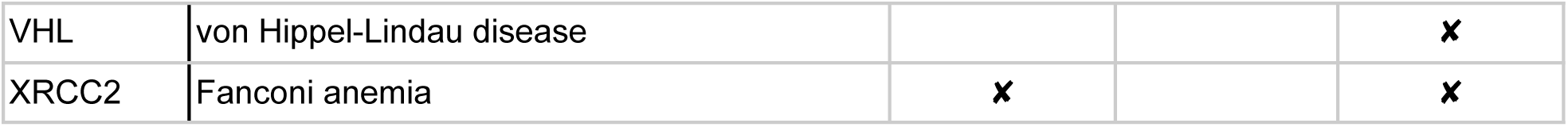
Table of gene-disease pairs that were excluded from biobank validation, detailing for each pair whether it was excluded because the mode of inheritance is not autosomal dominant, due to the absence of a matching phenotype in AoU, or due to a lack of participants with the phenotype and variants in the gene.

**Supplementary Table 3:**
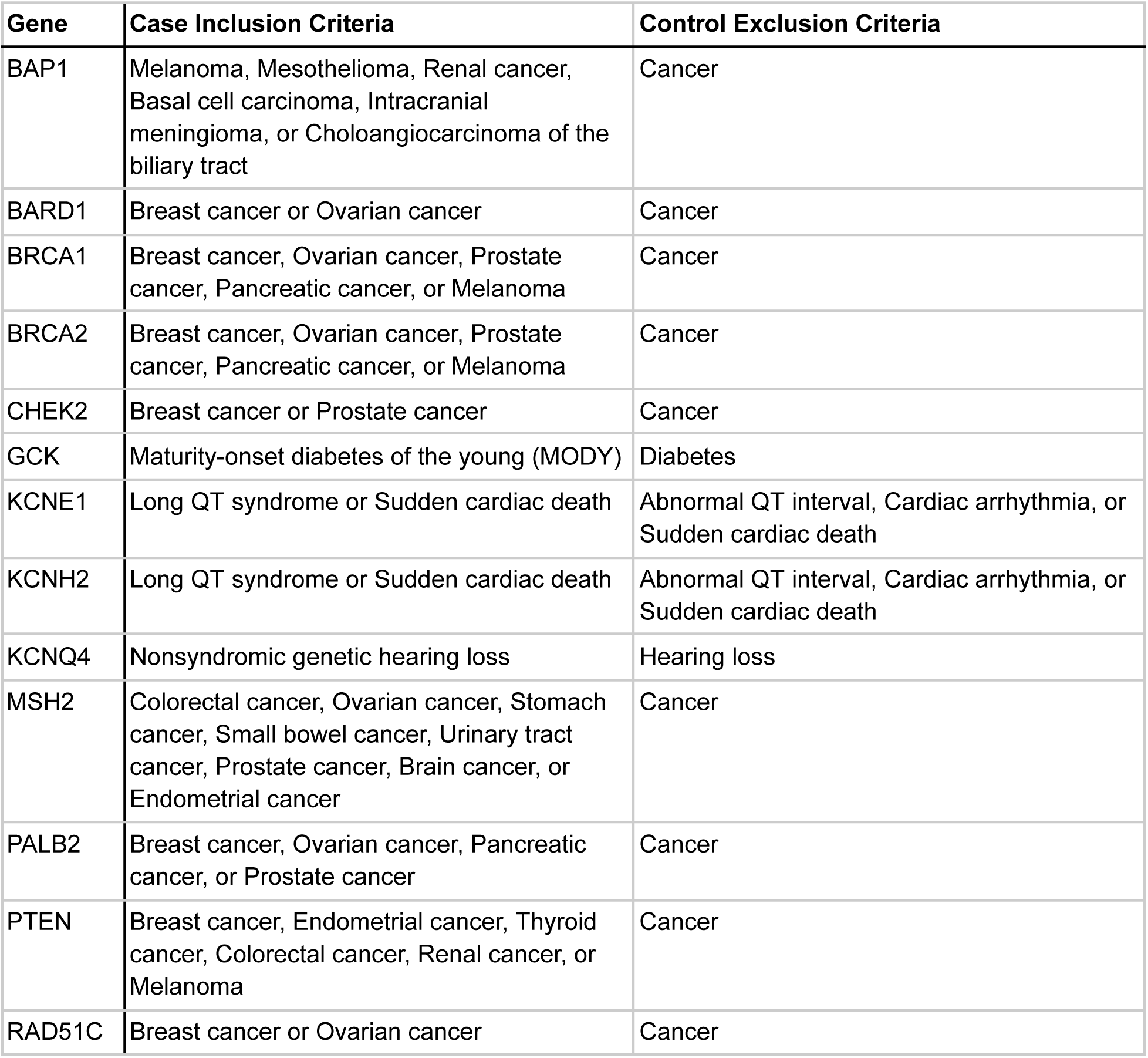
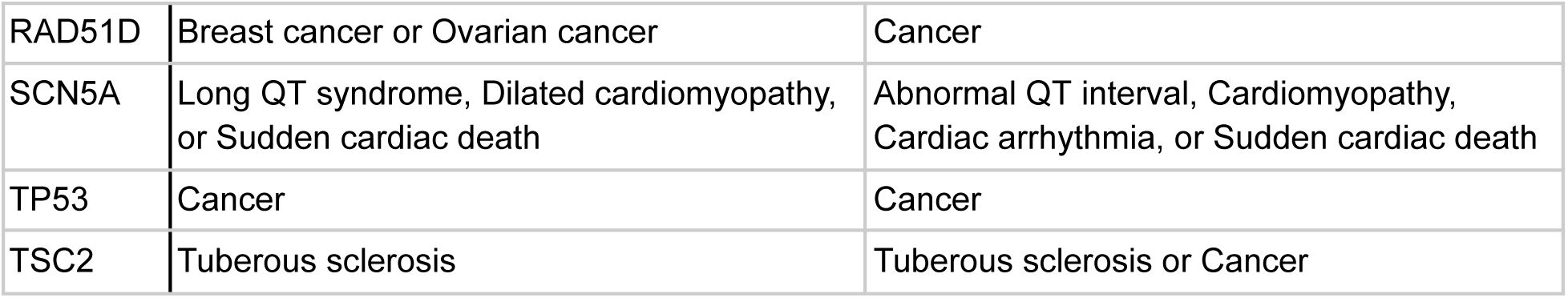
Table of gene-disease pairs detailing for each gene the criteria for including participants in the gene’s case cohort and the criteria for excluding participants from the gene’s control cohort.

## References

1. Landrum, M. J. et al. ClinVar: improving access to variant interpretations and supporting evidence. Nucleic Acids Res. 46, D1062–D1067 (2018).

2. DiStefano, M. T. et al. The Gene Curation Coalition: A global effort to harmonize gene-disease evidence resources. Genet. Med. 24, 1732–1742 (2022).

3. McEwen, A. E., Tejura, M., Fayer, S., Starita, L. M. & Fowler, D. M. Multiplexed assays of variant effect for clinical variant interpretation. Nat. Rev. Genet. (2025) doi:10.1038/s41576-025-00870-x.

4. Chen, E. et al. Rates and classification of variants of uncertain significance in hereditary disease genetic testing. *JAMA Netw*. Open 6, e2339571 (2023).

5. Gold, J. I. et al. Racial and socioeconomic disparities in genetic evaluation and testing in the adult patient population. Am. J. Hum. Genet. 113, 29–40 (2026).

6. Sirugo, G., Williams, S. M. & Tishkoff, S. A. The missing diversity in human genetic studies. Cell 177, 26–31 (2019).

7. Popejoy, A. B. et al. The clinical imperative for inclusivity: Race, ethnicity, and ancestry (REA) in genomics. Hum. Mutat. 39, 1713–1720 (2018).

8. Bianco, B. C. F. & Planello, A. C. Variant classification of hereditary cancer genes is affected by genomic underrepresentation of admixed populations. Mol. Genet. Genomics 300, 86 (2025).

9. Green, E. D. et al. Strategic vision for improving human health at The Forefront of Genomics. Nature 586, 683–692 (2020).

10. Fowler, D. M. & Fields, S. Deep mutational scanning: a new style of protein science. Nat. Methods 11, 801–807 (2014).

11. Findlay, G. M., Boyle, E. A., Hause, R. J., Klein, J. C. & Shendure, J. Saturation editing of genomic regions by multiplex homology-directed repair. Nature 513, 120–123 (2014).

12. Matreyek, K. A. et al. Multiplex assessment of protein variant abundance by massively parallel sequencing. Nat. Genet. 50, 874–882 (2018).

13. Findlay, G. M. et al. Accurate classification of BRCA1 variants with saturation genome editing. Nature 562, 217–222 (2018).

14. Jia, X. et al. Massively parallel functional testing of MSH2 missense variants conferring Lynch syndrome risk. Am. J. Hum. Genet. 108, 163–175 (2021).

15. Giacomelli, A. O. et al. Mutational processes shape the landscape of TP53 mutations in human cancer. Nat. Genet. 50, 1381–1387 (2018).

16. Boettcher, S. et al. A dominant-negative effect drives selection of TP53 missense mutations in myeloid malignancies. Science 365, 599–604 (2019).

17. Dawood, M. et al. Using multiplexed functional data to reduce variant classification inequities in underrepresented populations. Genome Med. 16, 143 (2024).

18. Cheng, J. et al. Accurate proteome-wide missense variant effect prediction with AlphaMissense. Science 381, eadg7492 (2023).

19. Orenbuch, R. et al. Deep generative modeling of the human proteome reveals over a hundred novel genes involved in rare genetic disorders. Res. Sq. (2024) doi:10.21203/rs.3.rs-3740259/v1.

20. Ioannidis, N. M. et al. REVEL: An ensemble method for predicting the pathogenicity of rare missense variants. Am. J. Hum. Genet. 99, 877–885 (2016).

21. Pejaver, V. et al. Inferring the molecular and phenotypic impact of amino acid variants with MutPred2. Nat. Commun. 11, 5918 (2020).

22. Livesey, B. J. & Marsh, J. A. Using deep mutational scanning to benchmark variant effect predictors and identify disease mutations. Mol. Syst. Biol. 16, e9380 (2020).

23. Livesey, B. J. & Marsh, J. A. Updated benchmarking of variant effect predictors using deep mutational scanning. Mol. Syst. Biol. 19, e11474 (2023).

24. Rubin, A. F. et al. MaveDB v2: a curated community database with over three million variant effects from multiplexed functional assays. Genomics (2021).

25. Brnich, S. E. et al. Recommendations for application of the functional evidence PS3/BS3 criterion using the ACMG/AMP sequence variant interpretation framework. Genome Med. 12, 3 (2019).

26. Park, M. S. et al. Insights on improving accessibility and usability of functional data to unlock their potential for variant interpretation. Am. J. Hum. Genet. 112, 1468–1478 (2025).

27. Starita, L. M. et al. Variant interpretation: Functional assays to the rescue. Am. J. Hum. Genet. 101, 315–325 (2017).

28. Fowler, D. M. & Rehm, H. L. Will variants of uncertain significance still exist in 2030? Am. J. Hum. Genet. 111, 5–10 (2024).

29. Fowler, D. M. et al. An Atlas of Variant Effects to understand the genome at nucleotide resolution. Genome Biol. 24, 147 (2023).

30. IGVF Consortium. Deciphering the impact of genomic variation on function. Nature 633, 47–57 (2024).

31. Chen, Y. et al. Multi-objective prioritization of genes for high-throughput functional assays towards improved clinical variant classification. Pac. Symp. Biocomput. 28, 323–334 (2023).

32. Popp, N. A. et al. Multiplex, multimodal mapping of variant effects in secreted proteins. Genomics (2024).

33. Zeiberg, D. et al. Gene-based calibration of high-throughput functional assays for clinical variant classification. bioRxivorg (2026) doi:10.1101/2025.04.29.651326.

34. Biar, C. G. et al. An integrated, scaled approach to resolve TSC2 variants of uncertain significance. bioRxiv 2026.01.16.699909 (2026) doi:10.64898/2026.01.16.699909.

35. Geck, R.C. et al. Evidence for G6PD variant classification from multiplexed functional assays. bioRxivorg (2025) doi:10.1101/2025.08.11.669723.

36. Woo, I. et al. Saturation genome editing of BARD1 resolves VUS and provides insight into BRCA1-BARD1 tumor suppression. medRxiv (2025) doi:10.1101/2025.11.03.25339440.

37. Lacoste, J. et al. Pervasive mislocalization of pathogenic coding variants underlying human disorders. Cell 187, 6725–6741.e13 (2024).

38. McEwen, A. E. et al. MaveMD: A functional data resource for genomic medicine. Genetic and Genomic Medicine (2025).

39. Claussnitzer, M. et al. Minimum information and guidelines for reporting a multiplexed assay of variant effect. Genome Biol. 25, 100 (2024).

40. Zeiberg, D., Jain, S. & Radivojac, P. Fast nonparametric estimation of class proportions in the positive-unlabeled classification setting. Proc. Conf. AAAI Artif. Intell. 34, 6729–6736 (2020).

41. Li, D. et al. The IGVF catalog-from genetic variation to function. Nucleic Acids Res. 54, D1437–D1445 (2026).

42. Chen, Y. et al. in preparation. Gene-and domain-aware calibration increases the clinical utility of variant effect predictors. bioRxiv (2026) doi: 10.64898/2026.02.17.706269.

43. Pejaver, V. et al. Calibration of computational tools for missense variant pathogenicity classification and ClinGen recommendations for PP3/BP4 criteria. Am. J. Hum. Genet. 109, 2163–2177 (2022).

44. Bergquist, T. et al. Calibration of additional computational tools expands ClinGen recommendation options for variant classification with PP3/BP4 criteria. Genet. Med. 27, 101402 (2025).

45. Rubin, A. F. et al. MaveDB 2024: a curated community database with over seven million variant effects from multiplexed functional assays. Genome Biol. 26, 13 (2025).

46. Yue, P., Li, Z. & Moult, J. Loss of protein structure stability as a major causative factor in monogenic disease. J. Mol. Biol. 353, 459–473 (2005).

47. Redler, R. L., Das, J., Diaz, J. R. & Dokholyan, N. V. Protein destabilization as a common factor in diverse inherited disorders. J. Mol. Evol. 82, 11–16 (2016).

48. Wang, Z. & Moult, J. SNPs, protein structure, and disease. Hum. Mutat. 17, 263–270 (2001).

49. Taipale, M. Disruption of protein function by pathogenic mutations: common and uncommon mechanisms 1. Biochem. Cell Biol. 97, 46–57 (2019).

50. Breast Cancer Association Consortium et al. Breast cancer risk genes - association analysis in more than 113,000 women. N. Engl. J. Med. 384, 428–439 (2021).

51. Hu, C. et al. A population-based study of genes previously implicated in breast cancer. N. Engl. J. Med. 384, 440–451 (2021).

52. Fu, J. M. et al. Rare coding variation provides insight into the genetic architecture and phenotypic context of autism. Nat. Genet. 54, 1320–1331 (2022).

53. Dawood, M. et al. GREGoR: accelerating genomics for rare diseases. Nature 647, 331–342 (2025).

54. Rubinstein, W. S. et al. The NIH genetic testing registry: a new, centralized database of genetic tests to enable access to comprehensive information and improve transparency. Nucleic Acids Res. 41, D925–35 (2013).

55. Lee, K. et al. ACMG SF v3.3 list for reporting of secondary findings in clinical exome and genome sequencing: A policy statement of the American College of Medical Genetics and Genomics (ACMG). Genet. Med. 27, 101454 (2025).

56. Chen, S. et al. A genomic mutational constraint map using variation in 76,156 human genomes. Nature 625, 92–100 (2024).

57. Richards, S. et al. Standards and guidelines for the interpretation of sequence variants: a joint consensus recommendation of the American College of Medical Genetics and Genomics and the Association for Molecular Pathology. Genet. Med. 17, 405–424 (2015).

58. Tavtigian, S. V. et al. Modeling the ACMG/AMP variant classification guidelines as a Bayesian classification framework. Genet. Med. 20, 1054–1060 (2018).

59. Tavtigian, S. V., Harrison, S. M., Boucher, K. M. & Biesecker, L. G. Fitting a naturally scaled point system to the ACMG/AMP variant classification guidelines. Hum. Mutat. 41, 1734–1737 (2020).

60. Tejura, M. et al. Calibration of variant effect predictors on genome-wide data masks heterogeneous performance across genes. Am. J. Hum. Genet. 111, 2031–2043 (2024).

61. Fayer, S. et al. Closing the gap: Systematic integration of multiplexed functional data resolves variants of uncertain significance in BRCA1, TP53, and PTEN. Am. J. Hum. Genet. 108, 2248–2258 (2021).

62. Scott, A. et al. Saturation-scale functional evidence supports clinical variant interpretation in Lynch syndrome. Genome Biol. 23, 266 (2022).

63. Rivera-Muñoz, E. A. et al. ClinGen Variant Curation Expert Panel experiences and standardized processes for disease and gene-level specification of the ACMG/AMP guidelines for sequence variant interpretation. Hum. Mutat. 39, 1614–1622 (2018).

64. Wu, D. et al. How I curate: applying American Society of Hematology-Clinical Genome Resource Myeloid Malignancy Variant Curation Expert Panel rules for RUNX1 variant curation for germline predisposition to myeloid malignancies. Haematologica 105, 870–887 (2020).

65. Costa, M., García S, A., León, A. & Pastor, O. The promises and pitfalls of automated variant interpretation: a comprehensive review. Brief. Bioinform. 26, (2025).

66. Plon, S. E. et al. Sequence variant classification and reporting: recommendations for improving the interpretation of cancer susceptibility genetic test results. Hum. Mutat. 29, 1282–1291 (2008).

67. Mersch, J. et al. Prevalence of variant reclassification following hereditary cancer genetic testing. JAMA 320, 1266–1274 (2018).

68. Hahn, E. et al. Variant classification changes over time in the clinical molecular diagnostic laboratory setting. J. Med. Genet. 61, 788–793 (2024).

69. Tsai, G. J. et al. Outcomes of 92 patient-driven family studies for reclassification of variants of uncertain significance. Genet. Med. 21, 1435–1442 (2019).

70. Shirts, B. H., Pritchard, C. C. & Walsh, T. Family-specific variants and the limits of human genetics. Trends Mol. Med. 22, 925–934 (2016).

71. Rapaport, F. et al. Negative selection on human genes underlying inborn errors depends on disease outcome and both the mode and mechanism of inheritance. Proc. Natl. Acad. Sci. U. S. A. 118, e2001248118 (2021).

72. Weghorn, D. et al. Applicability of the mutation-selection balance model to population genetics of heterozygous protein-truncating variants in humans. Mol. Biol. Evol. 36, 1701–1710 (2019).

73. Cassa, C. A. et al. Estimating the selective effects of heterozygous protein-truncating variants from human exome data. Nat. Genet. 49, 806–810 (2017).

74. Gudmundsson, S. et al. Exploring penetrance of clinically relevant variants in over 800,000 humans from the Genome Aggregation Database. Nat. Commun. 16, 9623 (2025).

75. Karczewski, K. J. et al. The mutational constraint spectrum quantified from variation in 141,456 humans. Nature 581, 434–443 (2020).

76. Gudmundsson, S. et al. Variant interpretation using population databases: Lessons from gnomAD. Hum. Mutat. 43, 1012–1030 (2022).

77. Sahni, N. et al. Widespread macromolecular interaction perturbations in human genetic disorders. Cell 161, 647–660 (2015).

78. Fragoza, R. et al. Extensive disruption of protein interactions by genetic variants across the allele frequency spectrum in human populations. Nat. Commun. 10, 4141 (2019).

79. Santoro, C. et al. Inhibitors in hemophilia B. Semin. Thromb. Hemost. 44, 578–589 (2018).

80. Allen, S. et al. Workshop report: the clinical application of data from multiplex assays of variant effect (MAVEs), 12 July 2023. Eur. J. Hum. Genet. 32, 593–600 (2024).

81. Villani, R. M. et al. Consultation informs strategies for improving the use of functional evidence in variant classification. Am. J. Hum. Genet. 112, 1489–1495 (2025).

82. McLaren, W. et al. The Ensembl Variant Effect Predictor. Genome Biol. 17, 122 (2016).

83. Sherry, S. T., Ward, M. & Sirotkin, K. dbSNP-database for single nucleotide polymorphisms and other classes of minor genetic variation. Genome Res. 9, 677–679 (1999).

84. Zhou, H. et al. FAVOR: functional annotation of variants online resource and annotator for variation across the human genome. Nucleic Acids Res. 51, D1300–D1311 (2023).

85. Parsons, M. T. et al. Evidence-based recommendations for gene-specific ACMG/AMP variant classification from the ClinGen ENIGMA BRCA1 and BRCA2 Variant Curation Expert Panel. Am. J. Hum. Genet. 111, 2044–2058 (2024).

86. Fortuno, C. et al. A quantitative, Bayesian-informed approach to gene-specific variant classification: Updated Expert Panel recommendations improve classification of TP53 germline variants for Li-Fraumeni syndrome. Genome Med. 17, 128 (2025).

86. Fowler, D. et al. Atlas of Variant Effects 2030 Roadmap: resolving human variants of uncertain significance. Preprint at 10.5281/ZENODO.15420413 (2025).

88. Arbesfeld, J. A. et al. Mapping MAVE data for use in human genomics applications. Genome Biol. 26, 179 (2025).

89. Sinnott-Armstrong, N. et al. Understanding genetic variants in context. Elife 13, (2024).

90. Simon, J. J., Fowler, D. M. & Maly, D. J. Multiplexed profiling of intracellular protein abundance, activity, interactions and druggability with LABEL-seq. Nat. Methods 21, 2094–2106 (2024).

91. Pendyala, S. et al. Image-based, pooled phenotyping reveals multidimensional, disease-specific variant effects. bioRxivorg (2025) doi:10.1101/2025.07.03.663081.

92. Fayer, S. et al. Editing stem cell genomes at scale to measure variant effects in diverse cell and genetic contexts. medRxiv (2025) doi:10.1101/2025.11.12.25340127.

93. Friedman, C. E. et al. CRaTER enrichment for on-target gene editing enables generation of variant libraries in hiPSCs. J. Mol. Cell. Cardiol. 179, 60–71 (2023).

94. Luck, K. et al. A reference map of the human binary protein interactome. Nature 580, 402–408 (2020).

95. Choi, S. G. et al. Maximizing binary interactome mapping with a minimal number of assays. Nat. Commun. 10, 3907 (2019).

96. Cimini, B. A. et al. Optimizing the Cell Painting assay for image-based profiling. Nat. Protoc. 18, 1981–2013 (2023).

97. Stirling, D. R. et al. CellProfiler 4: improvements in speed, utility and usability. BMC Bioinformatics 22, 433 (2021).

98. Weisbart, E. et al. Cell Painting Gallery: an open resource for image-based profiling. Nat. Methods 21, 1775–1777 (2024).

99. Serrano, E. et al. Reproducible image-based profiling with Pycytominer. Nat. Methods 22, 677–680 (2025).

100. Hart, R. K. et al. HGVS Nomenclature 2024: improvements to community engagement, usability, and computability. Genome Med. 16, 149 (2024).

101. den Dunnen, J. T. et al. HGVS recommendations for the description of sequence variants: 2016 update. Hum. Mutat. 37, 564–569 (2016).

102. Freeman, P. J. et al. Standardizing variant naming in literature with VariantValidator to increase diagnostic rates. Nat. Genet. 56, 2284–2286 (2024).

103. Freeman, P. J., Hart, R. K., Gretton, L. J., Brookes, A. J. & Dalgleish, R. VariantValidator: Accurate validation, mapping, and formatting of sequence variation descriptions. Hum. Mutat. 39, 61–68 (2018).

104. Jaganathan, K. et al. Predicting Splicing from Primary Sequence with Deep Learning. Cell 176, 535–548.e24 (2019).

105. Gebbia, M. et al. A missense variant effect map for the human tumor-suppressor protein CHK2. Am. J. Hum. Genet. 111, 2675–2692 (2024).

106. All of Us Research Program Genomics Investigators. Genomic data in the All of Us Research Program. Nature 627, 340–346 (2024).

107. All of Us Genomic Quality Report. *User Support* https://support.researchallofus.org/hc/en-us/articles/29390274413716-All-of-Us-Genomic-Quality-Report.

108. Mighell, T. L., Evans-Dutson, S. & O’Roak, B. J. A saturation Mutagenesis approach to understanding PTEN lipid phosphatase activity and genotype-phenotype relationships. Am. J. Hum. Genet. 102, 943–955 (2018).

109. O’Roak, B. J. et al. Multiplex targeted sequencing identifies recurrently mutated genes in autism spectrum disorders. Science 338, 1619–1622 (2012).

