## Extended Data Figure 1 and 3 for "A scalable approach to resolving variants of uncertain significance"

ASPA_Grønbæk-Thygesen_2024_abundance

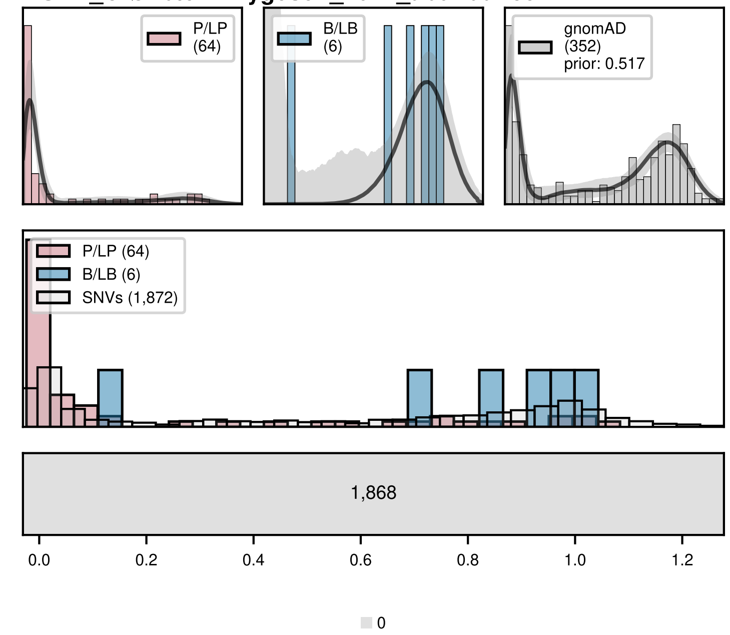


ASPA_Grønbæk-Thygesen_2024_toxicity

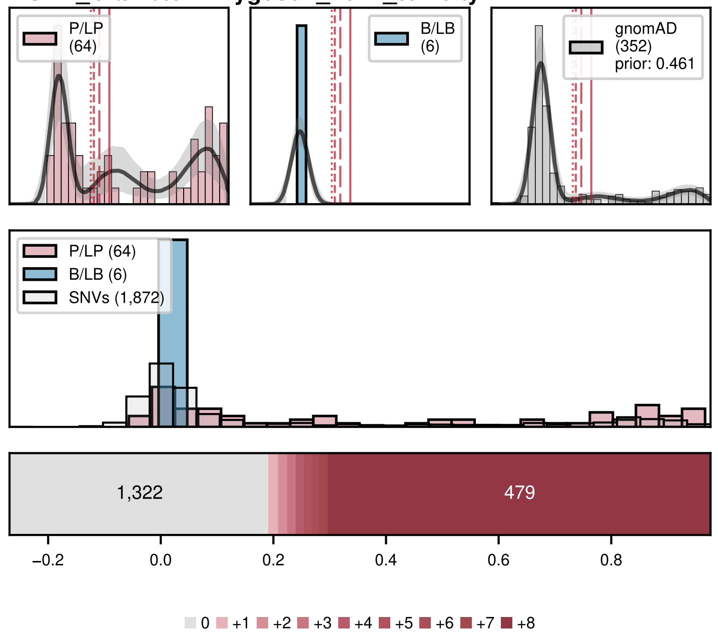


BAP1_Waters_2024

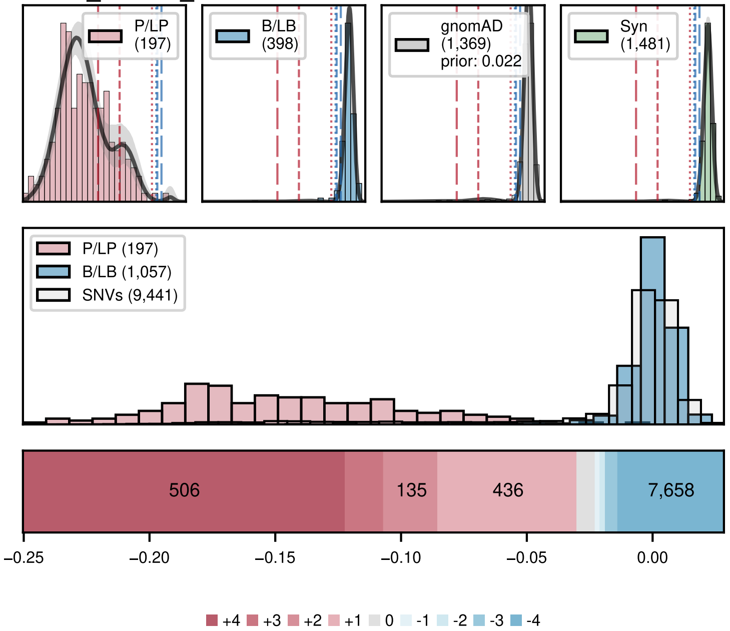


BARD1_IGVF

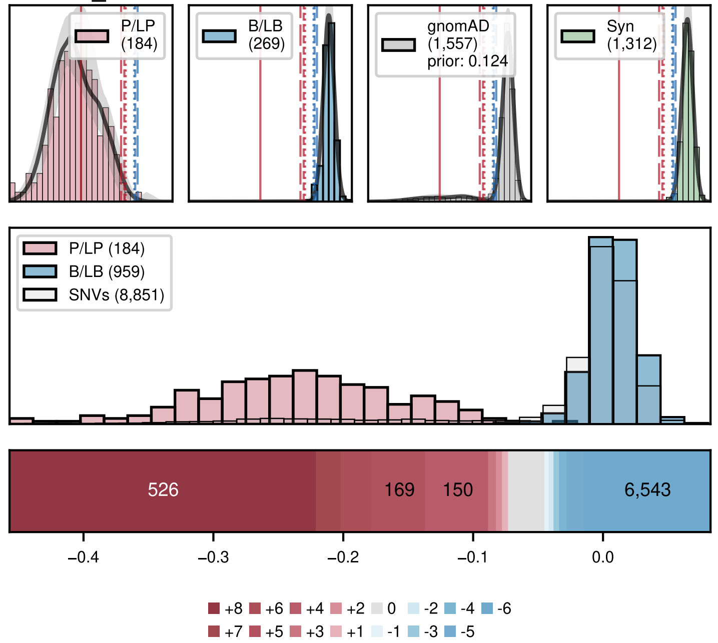


BRCA1_Adamovich_2022_Cisplatin_Resistance

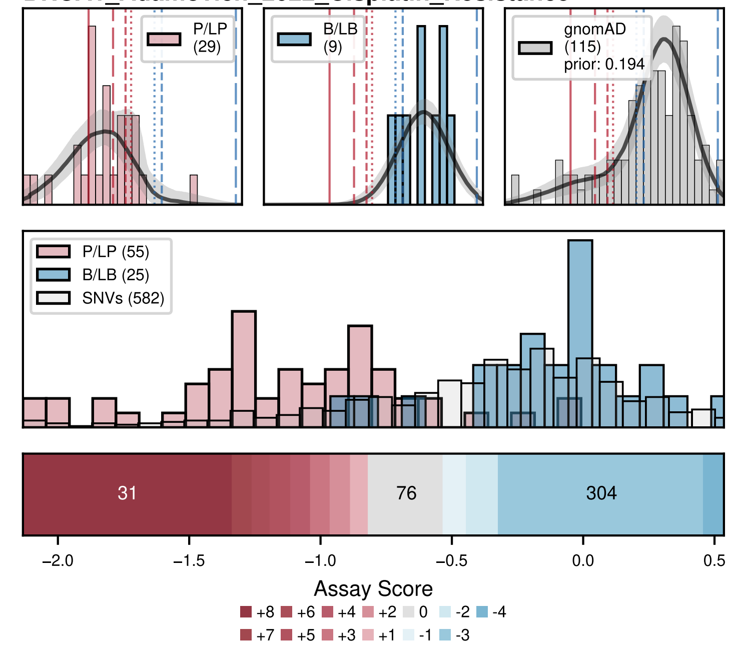


BRCA1_Adamovich_2022_HDR

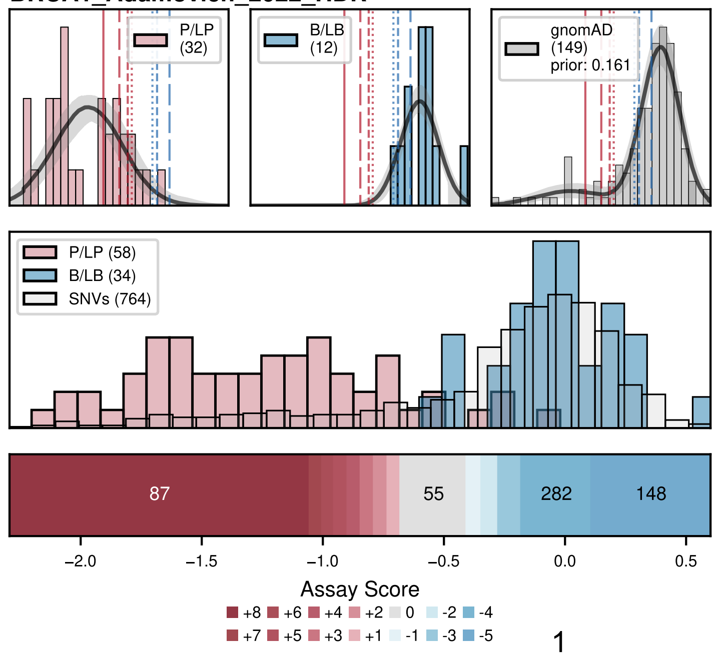

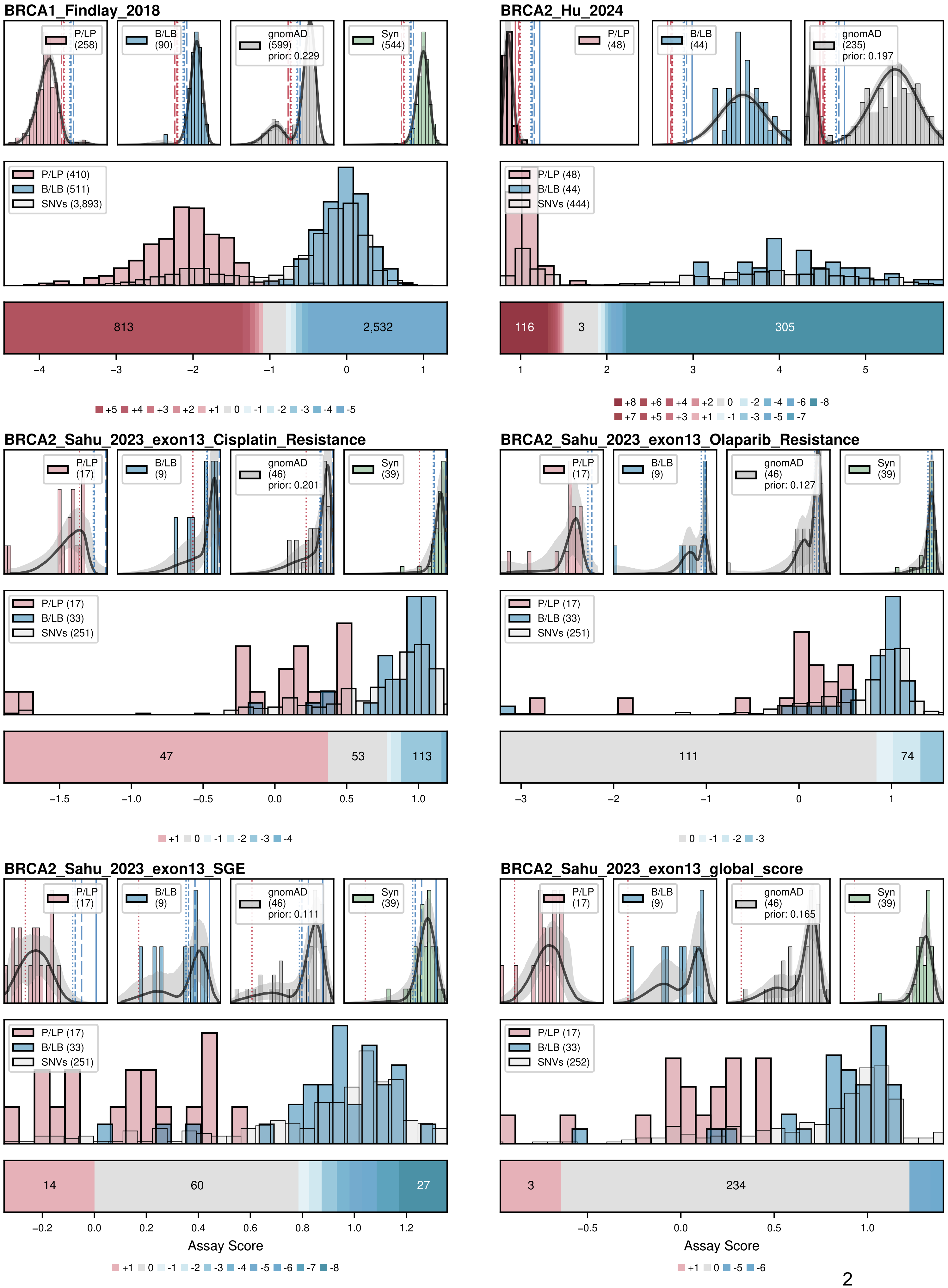

BRCA1_Findlay_2018
P/LP (258)
B/LB (90)
gnomAD (599) prior: 0.229
Syn (544)
P/LP (410)
B/LB (511)
SNVs (3,893)
813
2,532
−4 −3 −2 −1 0 1
+5 +4 +3 +2 +1 0 -1 -2 -3 -4 -5
BRCA2_Hu_2024
P/LP (48)
B/LB (44)
gnomAD (235) prior: 0.197
P/LP (48)
B/LB (44)
SNVs (444)
116
3
305
1 2 3 4 5
+8 +6 +4 +2 0 -2 -4 -6 -8
+7 +5 +3 +1 -1 -3 -5 -7
BRCA2_Sahu_2023_exon13_Cisplatin_Resistance
P/LP (17)
B/LB (9)
gnomAD (46) prior: 0.201
Syn (39)
P/LP (17)
B/LB (33)
SNVs (251)
47
53
113
−1.5 −1.0 −0.5 0.0 0.5 1.0
+1 0 -1 -2 -3 -4
BRCA2_Sahu_2023_exon13_Olaparib_Resistance
P/LP (17)
B/LB (9)
gnomAD (46) prior: 0.127
Syn (39)
P/LP (17)
B/LB (33)
SNVs (251)
111
74
−3 −2 −1 0 1
0 -1 -2 -3
BRCA2_Sahu_2023_exon13_SGE
P/LP (17)
B/LB (9)
gnomAD (46) prior: 0.111
Syn (39)
P/LP (17)
B/LB (33)
SNVs (251)
14
60
27
−0.2 0.0 0.2 0.4 0.6 0.8 1.0 1.2
Assay Score
+1 0 -1 -2 -3 -4 -5 -6 -7 -8
BRCA2_Sahu_2023_exon13_global_score
P/LP (17)
B/LB (9)
gnomAD (46) prior: 0.165
Syn (39)
P/LP (17)
B/LB (33)
SNVs (252)
3
234
−0.5 0.0 0.5 1.0
Assay Score
+1 0 -5 -6

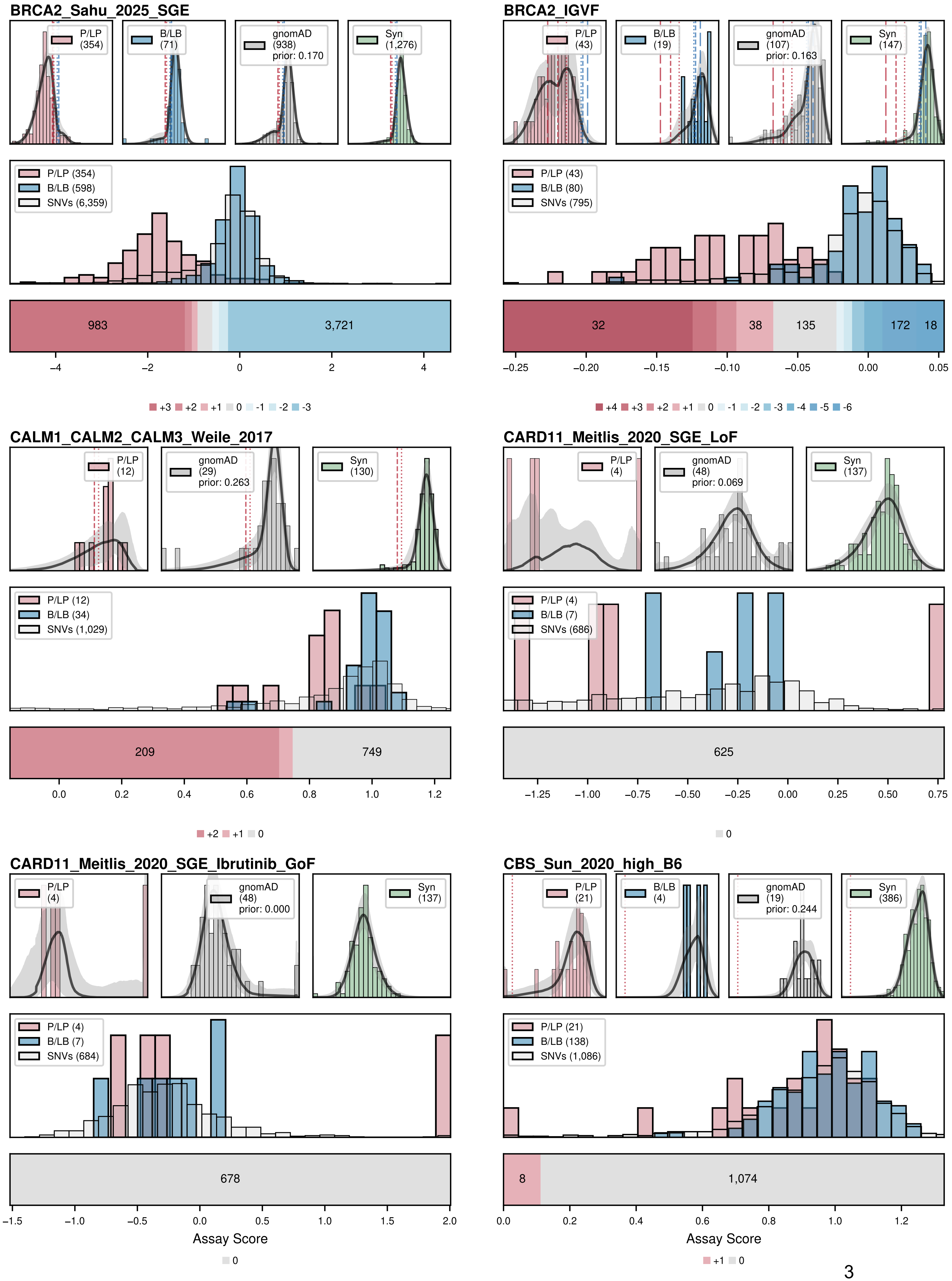

BRCA2_Sahu_2025_SGE
P/LP (354)
B/LB (71)
gnomAD (938) prior: 0.170
Syn (1,276)
P/LP (354)
B/LB (598)
SNVs (6,359)
983
3,721
+3 +2 +1 0 -1 -2 -3
BRCA2_IGVF
P/LP (43)
B/LB (19)
gnomAD (107) prior: 0.163
Syn (147)
P/LP (43)
B/LB (80)
SNVs (795)
32
38
135
172
18
+4 +3 +2 +1 0 -1 -2 -3 -4 -5 -6
CALM1_CALM2_CALM3_Weile_2017
P/LP (12)
gnomAD (29) prior: 0.263
Syn (130)
P/LP (12)
B/LB (34)
SNVs (1,029)
209
749
+2 +1 0
CARD11_Meitlis_2020_SGE_LoF
P/LP (4)
gnomAD (48) prior: 0.069
Syn (137)
P/LP (4)
B/LB (7)
SNVs (686)
625
0
CARD11_Meitlis_2020_SGE_Ibrutinib_GoF
P/LP (4)
gnomAD (48) prior: 0.000
Syn (137)
P/LP (4)
B/LB (7)
SNVs (684)
678
Assay Score
0
CBS_Sun_2020_high_B6
P/LP (21)
B/LB (4)
gnomAD (19) prior: 0.244
Syn (386)
P/LP (21)
B/LB (138)
SNVs (1,086)
8
1,074
Assay Score
+1 0

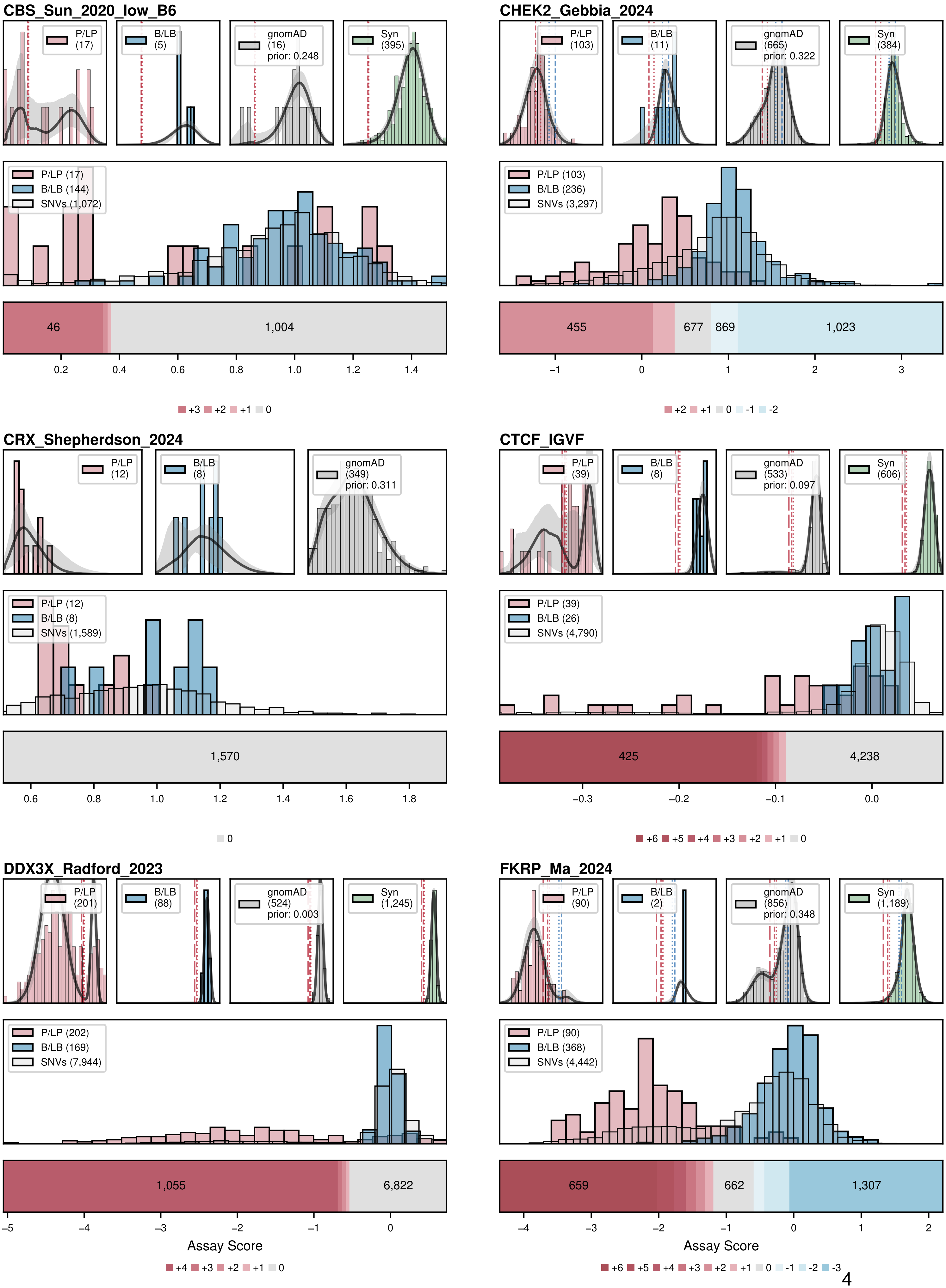
CBS_Sun_2020_low_B6
P/LP (17)
B/LB (5)
gnomAD (16) prior: 0.248
Syn (395)
P/LP (17)
B/LB (144)
SNVs (1,072)
46
1,004
0.2 0.4 0.6 0.8 1.0 1.2 1.4
+3 +2 +1 0
CHEK2_Gebbia_2024
P/LP (103)
B/LB (11)
gnomAD (665) prior: 0.322
Syn (384)
P/LP (103)
B/LB (236)
SNVs (3,297)
455
677
869
1,023
−1 0 1 2 3
+2 +1 0 -1 -2
CRX_Shepherdson_2024
P/LP (12)
B/LB (8)
gnomAD (349) prior: 0.311
P/LP (12)
B/LB (8)
SNVs (1,589)
1,570
0.6 0.8 1.0 1.2 1.4 1.6 1.8
0
CTCF_IGVF
P/LP (39)
B/LB (8)
gnomAD (533) prior: 0.097
Syn (606)
P/LP (39)
B/LB (26)
SNVs (4,790)
425
4,238
−0.3 −0.2 −0.1 0.0
+6 +5 +4 +3 +2 +1 0
DDX3X_Radford_2023
P/LP (201)
B/LB (88)
gnomAD (524) prior: 0.003
Syn (1,245)
P/LP (202)
B/LB (169)
SNVs (7,944)
1,055
6,822
−5 −4 −3 −2 −1 0
Assay Score
+4 +3 +2 +1 0
FKRP_Ma_2024
P/LP (90)
B/LB (2)
gnomAD (856) prior: 0.348
Syn (1,189)
P/LP (90)
B/LB (368)
SNVs (4,442)
659
662
1,307
−4 −3 −2 −1 0 1 2
Assay Score
+6 +5 +4 +3 +2 +1 0 -1 -2 -3

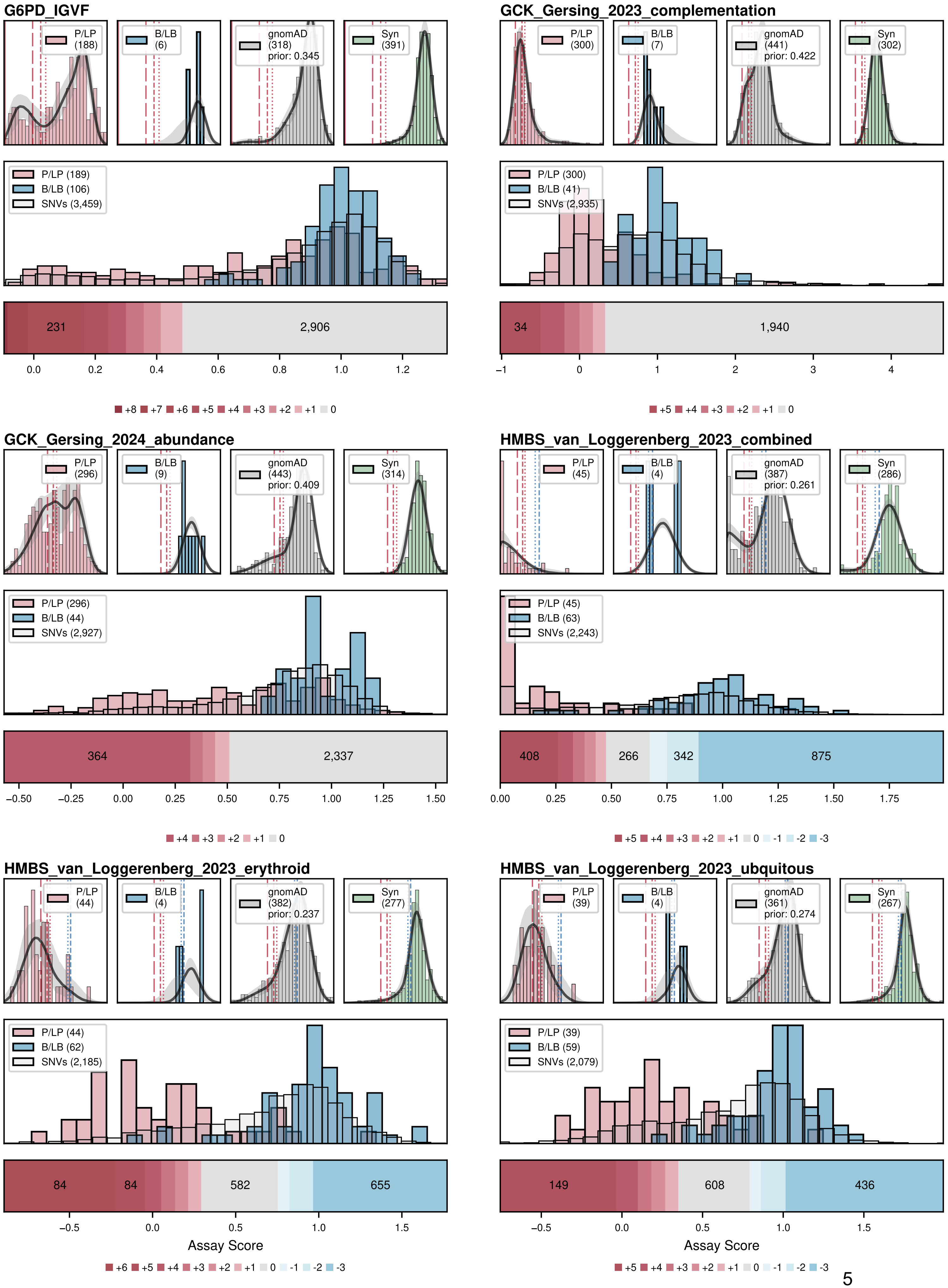
G6PD_IGVF
P/LP (188)
B/LB (6)
gnomAD (318) prior: 0.345
Syn (391)
P/LP (189)
B/LB (106)
SNVs (3,459)
231
2,906
0.0 0.2 0.4 0.6 0.8 1.0 1.2
+8 +7 +6 +5 +4 +3 +2 +1 0
GCK_Gersing_2023_complementation
P/LP (300)
B/LB (7)
gnomAD (441) prior: 0.422
Syn (302)
P/LP (300)
B/LB (41)
SNVs (2,935)
34
1,940
−1 0 1 2 3 4
+5 +4 +3 +2 +1 0
GCK_Gersing_2024_abundance
P/LP (296)
B/LB (9)
gnomAD (443) prior: 0.409
Syn (314)
P/LP (296)
B/LB (44)
SNVs (2,927)
364
2,337
−0.50 −0.25 0.00 0.25 0.50 0.75 1.00 1.25 1.50
+4 +3 +2 +1 0
HMBS_van_Loggerenberg_2023_combined
P/LP (45)
B/LB (4)
gnomAD (387) prior: 0.261
Syn (286)
P/LP (45)
B/LB (63)
SNVs (2,243)
408
266
342
875
0.00 0.25 0.50 0.75 1.00 1.25 1.50 1.75
+5 +4 +3 +2 +1 0 -1 -2 -3
HMBS_van_Loggerenberg_2023_erythroid
P/LP (44)
B/LB (4)
gnomAD (382) prior: 0.237
Syn (277)
P/LP (44)
B/LB (62)
SNVs (2,185)
84
84
582
655
−0.5 0.0 0.5 1.0 1.5
Assay Score
+6 +5 +4 +3 +2 +1 0 -1 -2 -3
HMBS_van_Loggerenberg_2023_ubquitous
P/LP (39)
B/LB (4)
gnomAD (361) prior: 0.274
Syn (267)
P/LP (39)
B/LB (59)
SNVs (2,079)
149
608
436
−0.5 0.0 0.5 1.0 1.5
Assay Score
+5 +4 +3 +2 +1 0 -1 -2 -3

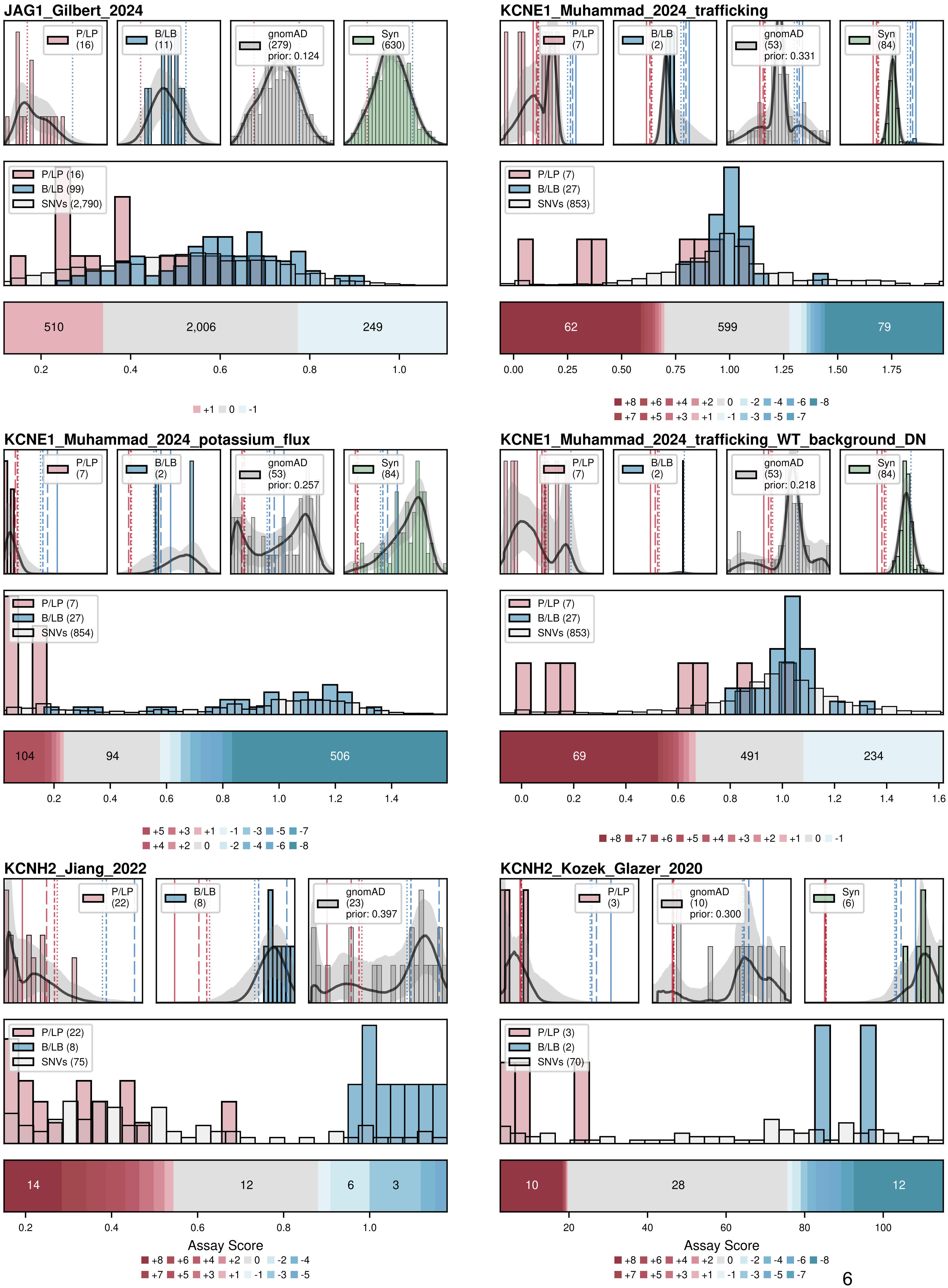
JAG1_Gilbert_2024
P/LP (16)
B/LB (11)
gnomAD (279) prior: 0.124
Syn (630)
P/LP (16)
B/LB (99)
SNVs (2,790)
510
2,006
249
+1 0 -1
KCNE1_Muhammad_2024_trafficking
P/LP (7)
B/LB (2)
gnomAD (53) prior: 0.331
Syn (84)
P/LP (7)
B/LB (27)
SNVs (853)
62
599
79
+8 +6 +4 +2 0 -2 -4 -6 -8
+7 +5 +3 +1 -1 -3 -5 -7
KCNE1_Muhammad_2024_potassium_flux
P/LP (7)
B/LB (2)
gnomAD (53) prior: 0.257
Syn (84)
P/LP (7)
B/LB (27)
SNVs (854)
104
94
506
+5 +3 +1 -1 -3 -5 -7
+4 +2 0 -2 -4 -6 -8
KCNE1_Muhammad_2024_trafficking_WT_background_DN
P/LP (7)
B/LB (2)
gnomAD (53) prior: 0.218
Syn (84)
P/LP (7)
B/LB (27)
SNVs (853)
69
491
234
+8 +7 +6 +5 +4 +3 +2 +1 0 -1
KCNH2_Jiang_2022
P/LP (22)
B/LB (8)
gnomAD (23) prior: 0.397
P/LP (22)
B/LB (8)
SNVs (75)
14
12
6
3
Assay Score
+8 +6 +4 +2 0 -2 -4
+7 +5 +3 +1 -1 -3 -5
KCNH2_Kozek_Glazer_2020
P/LP (3)
gnomAD (10) prior: 0.300
Syn (6)
P/LP (3)
B/LB (2)
SNVs (70)
10
28
12
Assay Score
+8 +6 +4 +2 0 -2 -4 -6 -8
+7 +5 +3 +1 -1 -3 -5 -7

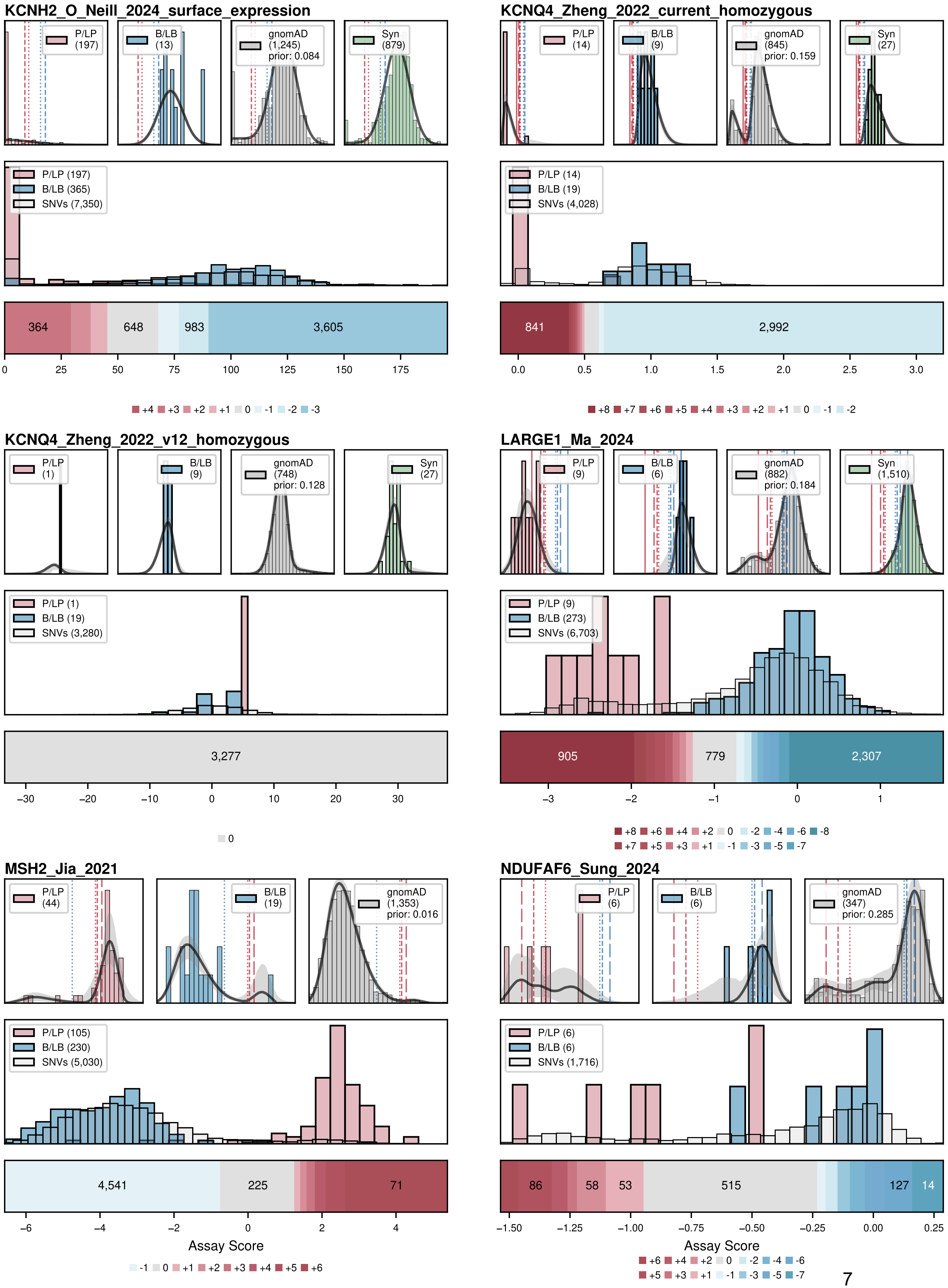
KCNH2_O_Neill_2024_surface_expression
P/LP (197)
B/LB (13)
gnomAD (1,245) prior: 0.084
Syn (879)
P/LP (197)
B/LB (365)
SNVs (7,350)
364
648
983
3,605
0 25 50 75 100 125 150 175
+4 +3 +2 +1 0 -1 -2 -3
KCNQ4_Zheng_2022_current_homozygous
P/LP (14)
B/LB (9)
gnomAD (845) prior: 0.159
Syn (27)
P/LP (14)
B/LB (19)
SNVs (4,028)
841
2,992
0.0 0.5 1.0 1.5 2.0 2.5 3.0
+8 +7 +6 +5 +4 +3 +2 +1 0 -1 -2
KCNQ4_Zheng_2022_v12_homozygous
P/LP (1)
B/LB (9)
gnomAD (748) prior: 0.128
Syn (27)
P/LP (1)
B/LB (19)
SNVs (3,280)
3,277
−30 −20 −10 0 10 20 30
0
LARGE1_Ma_2024
P/LP (9)
B/LB (6)
gnomAD (882) prior: 0.184
Syn (1,510)
P/LP (9)
B/LB (273)
SNVs (6,703)
905
779
2,307
−3 −2 −1 0 1
+8 +6 +4 +2 0 -2 -4 -6 -8
+7 +5 +3 +1 -1 -3 -5 -7
MSH2_Jia_2021
P/LP (44)
B/LB (19)
gnomAD (1,353) prior: 0.016
P/LP (105)
B/LB (230)
SNVs (5,030)
4,541
225
71
−6 −4 −2 0 2 4
Assay Score
-1 0 +1 +2 +3 +4 +5 +6
NDUFAF6_Sung_2024
P/LP (6)
B/LB (6)
gnomAD (347) prior: 0.285
P/LP (6)
B/LB (6)
SNVs (1,716)
86
58
53
515
127
14
−1.50 −1.25 −1.00 −0.75 −0.50 −0.25 0.00 0.25
Assay Score
+6 +4 +2 0 -2 -4 -6
+5 +3 +1 -1 -3 -5 -7

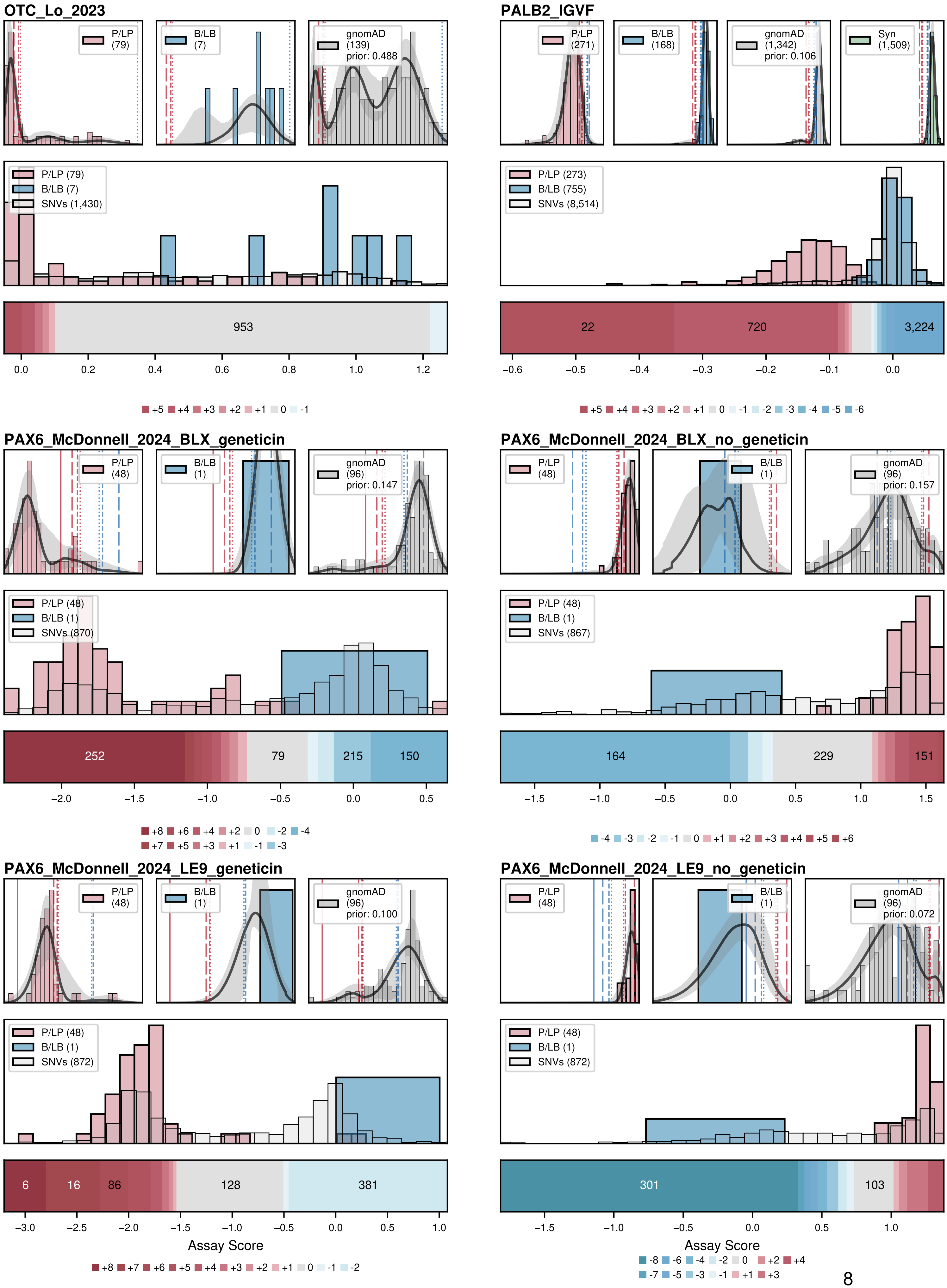
OTC_Lo_2023
P/LP (79)
B/LB (7)
gnomAD (139) prior: 0.488
P/LP (79)
B/LB (7)
SNVs (1,430)
953
0.0 0.2 0.4 0.6 0.8 1.0 1.2
+5 +4 +3 +2 +1 0 -1
PALB2_IGVF
P/LP (271)
B/LB (168)
gnomAD (1,342) prior: 0.106
Syn (1,509)
P/LP (273)
B/LB (755)
SNVs (8,514)
22
720
3,224
−0.6 −0.5 −0.4 −0.3 −0.2 −0.1 0.0
+5 +4 +3 +2 +1 0 -1 -2 -3 -4 -5 -6
PAX6_McDonnell_2024_BLX_geneticin
P/LP (48)
B/LB (1)
gnomAD (96) prior: 0.147
P/LP (48)
B/LB (1)
SNVs (870)
252
79
215
150
−2.0 −1.5 −1.0 −0.5 0.0 0.5
+8 +6 +4 +2 0 -2 -4
+7 +5 +3 +1 -1 -3
PAX6_McDonnell_2024_BLX_no_geneticin
P/LP (48)
B/LB (1)
gnomAD (96) prior: 0.157
P/LP (48)
B/LB (1)
SNVs (867)
164
229
151
−1.5 −1.0 −0.5 0.0 0.5 1.0 1.5
-4 -3 -2 -1 0 +1 +2 +3 +4 +5 +6
PAX6_McDonnell_2024_LE9_geneticin
P/LP (48)
B/LB (1)
gnomAD (96) prior: 0.100
P/LP (48)
B/LB (1)
SNVs (872)
6
16
86
128
381
−3.0 −2.5 −2.0 −1.5 −1.0 −0.5 0.0 0.5 1.0
Assay Score
+8 +7 +6 +5 +4 +3 +2 +1 0 -1 -2
PAX6_McDonnell_2024_LE9_no_geneticin
P/LP (48)
B/LB (1)
gnomAD (96) prior: 0.072
P/LP (48)
B/LB (1)
SNVs (872)
301
103
−1.5 −1.0 −0.5 0.0 0.5 1.0
Assay Score
-8 -6 -4 -2 0 +2 +4
-7 -5 -3 -1 +1 +3

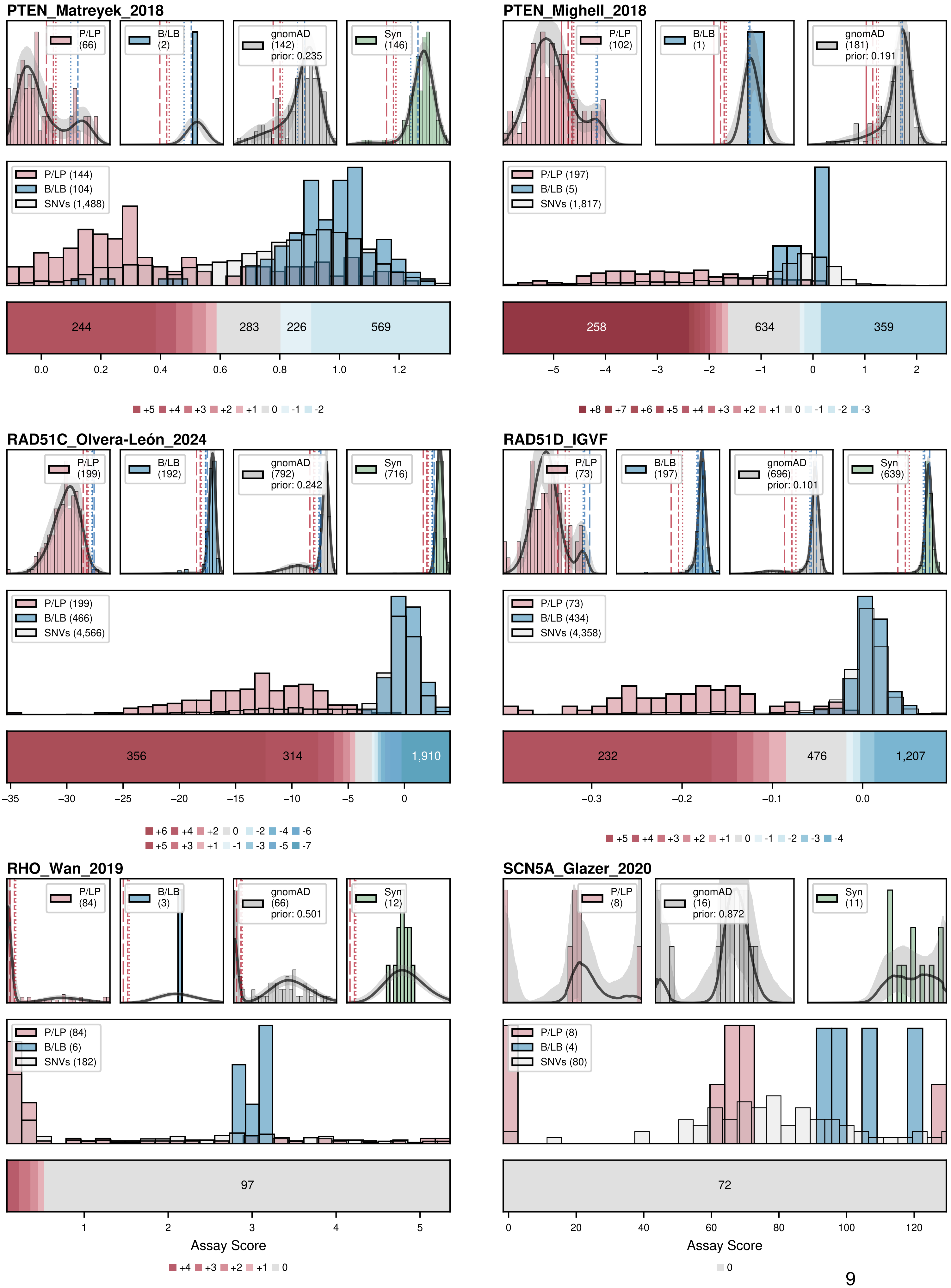
PTEN_Matreyek_2018
P/LP (66)
B/LB (2)
gnomAD (142) prior: 0.235
Syn (146)
P/LP (144)
B/LB (104)
SNVs (1,488)
244
283
226
569
0.0 0.2 0.4 0.6 0.8 1.0 1.2
+5 +4 +3 +2 +1 0 -1 -2
PTEN_Mighell_2018
P/LP (102)
B/LB (1)
gnomAD (181) prior: 0.191
P/LP (197)
B/LB (5)
SNVs (1,817)
258
634
359
−5 −4 −3 −2 −1 0 1 2
+8 +7 +6 +5 +4 +3 +2 +1 0 -1 -2 -3
RAD51C_Olvera-León_2024
P/LP (199)
B/LB (192)
gnomAD (792) prior: 0.242
Syn (716)
P/LP (199)
B/LB (466)
SNVs (4,566)
356
314
1,910
−35 −30 −25 −20 −15 −10 −5 0
+6 +4 +2 0 -2 -4 -6
+5 +3 +1 -1 -3 -5 -7
RAD51D_IGVF
P/LP (73)
B/LB (197)
gnomAD (696) prior: 0.101
Syn (639)
P/LP (73)
B/LB (434)
SNVs (4,358)
232
476
1,207
−0.3 −0.2 −0.1 0.0
+5 +4 +3 +2 +1 0 -1 -2 -3 -4
RHO_Wan_2019
P/LP (84)
B/LB (3)
gnomAD (66) prior: 0.501
Syn (12)
P/LP (84)
B/LB (6)
SNVs (182)
97
1 2 3 4 5
Assay Score
+4 +3 +2 +1 0
SCN5A_Glazer_2020
P/LP (8)
gnomAD (16) prior: 0.872
Syn (11)
P/LP (8)
B/LB (4)
SNVs (80)
72
0 20 40 60 80 100 120
Assay Score
0

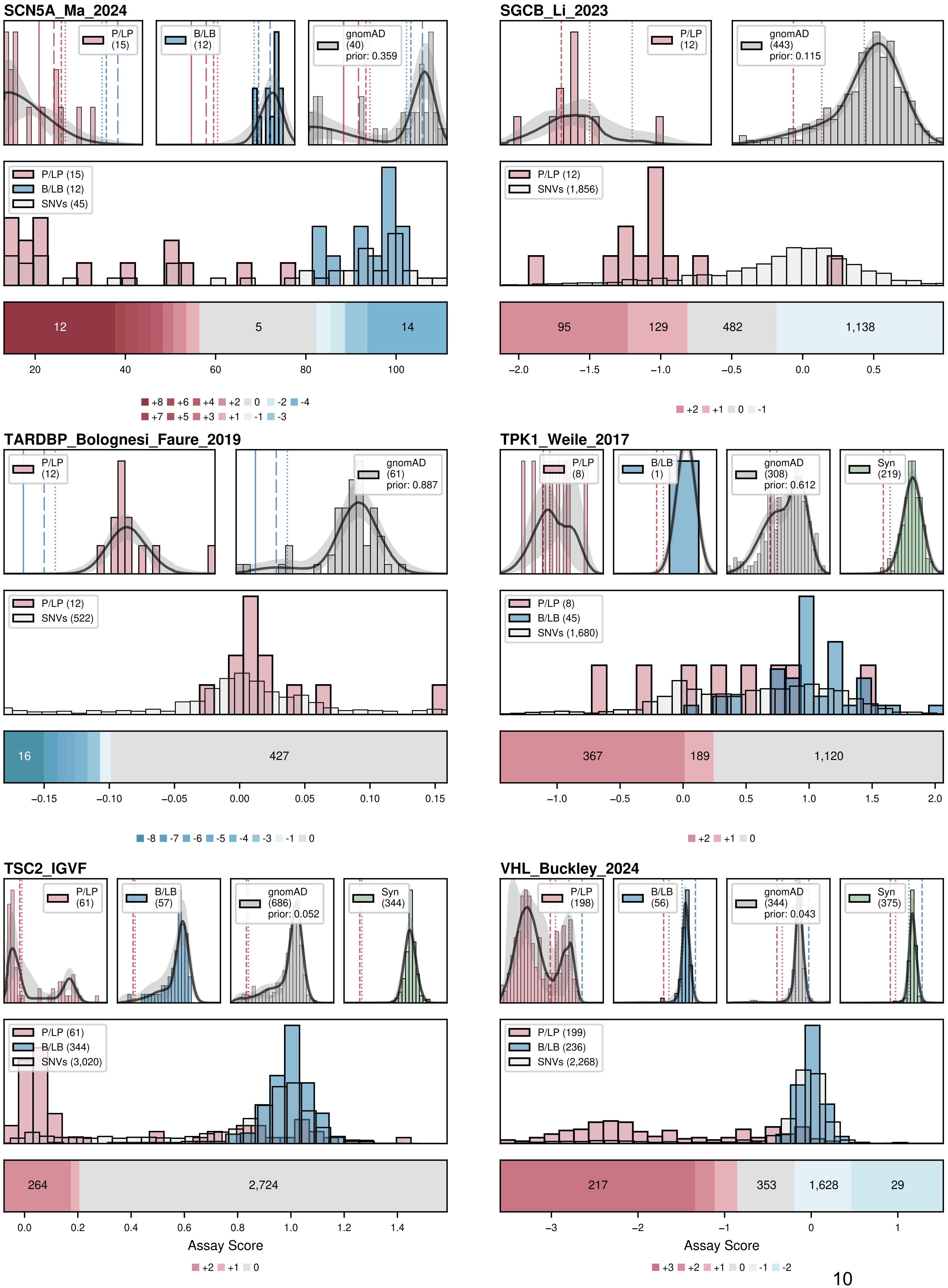
SCN5A_Ma_2024
P/LP (15)
B/LB (12)
gnomAD (40) prior: 0.359
P/LP (15)
B/LB (12)
SNVs (45)
12
5
14
20
40
60
80
100
+8 +6 +4 +2 0 -2 -4
+7 +5 +3 +1 -1 -3
SGCB_Li_2023
P/LP (12)
gnomAD (443) prior: 0.115
P/LP (12)
SNVs (1,856)
95
129
482
1,138
−2.0
−1.5
−1.0
−0.5
0.0
0.5
+2 +1 0 -1
TARDBP_Bolognesi_Faure_2019
P/LP (12)
gnomAD (61) prior: 0.887
P/LP (12)
SNVs (522)
16
427
−0.15
−0.10
−0.05
0.00
0.05
0.10
0.15
-8 -7 -6 -5 -4 -3 -1 0
TPK1_Weile_2017
P/LP (8)
B/LB (1)
gnomAD (308) prior: 0.612
Syn (219)
P/LP (8)
B/LB (45)
SNVs (1,680)
367
189
1,120
−1.0
−0.5
0.0
0.5
1.0
1.5
2.0
+2 +1 0
TSC2_IGVF
P/LP (61)
B/LB (57)
gnomAD (686) prior: 0.052
Syn (344)
P/LP (61)
B/LB (344)
SNVs (3,020)
264
2,724
0.0
0.2
0.4
0.6
0.8
1.0
1.2
1.4
Assay Score
+2 +1 0
VHL_Buckley_2024
P/LP (198)
B/LB (56)
gnomAD (344) prior: 0.043
Syn (375)
P/LP (199)
B/LB (236)
SNVs (2,268)
217
353
1,628
29
−3
−2
−1
0
1
Assay Score
+3 +2 +1 0 -1 -2

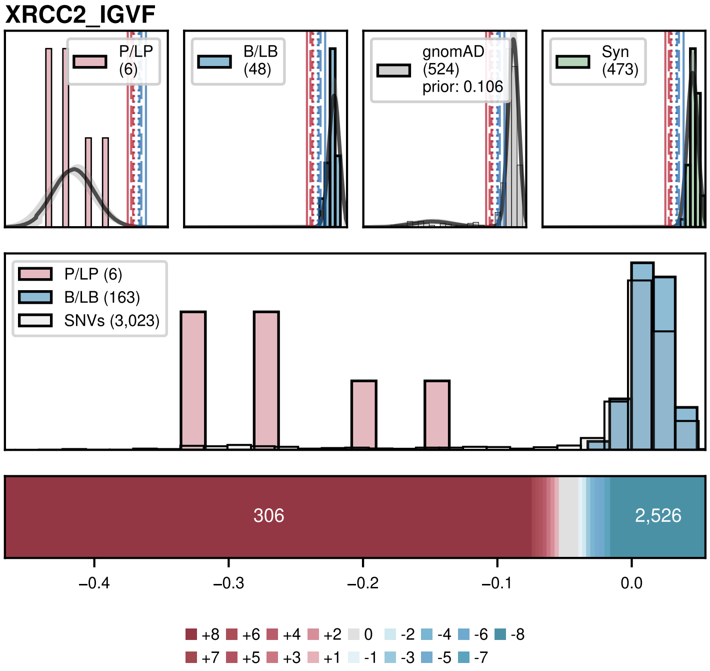


### Extended Data Figure 1: ExCALIBR model fits and assigned evidence for experimental datasets

ExCALIBR fits, variant score distributions, and calibrated evidence strengths are shown for 61 experimental datasets spanning 37 genes across 11 pages, with each dataset visualized in a single panel containing three components: top, ExCALIBR fits for pathogenic variants (red), benign variants (blue), gnomAD population variants (grey), and synonymous variants (green; exclusive with other variant classes when available), where ClinVar variants are from the December 2018 release for BRCA1, MSH2, TP53, and PTEN, and the January 2025 release for all other genes; middle, score distributions for pathogenic (red) and benign (blue) control variants from the January 2025 ClinVar release, overlaid with the distribution of all possible SNVs (light grey); bottom, score intervals corresponding to evidence strength assignments (up to 8 pathogenic or benign points) determined by ExCALIBR, with the number of SNVs assigned to each interval labeled within sufficiently large bins.

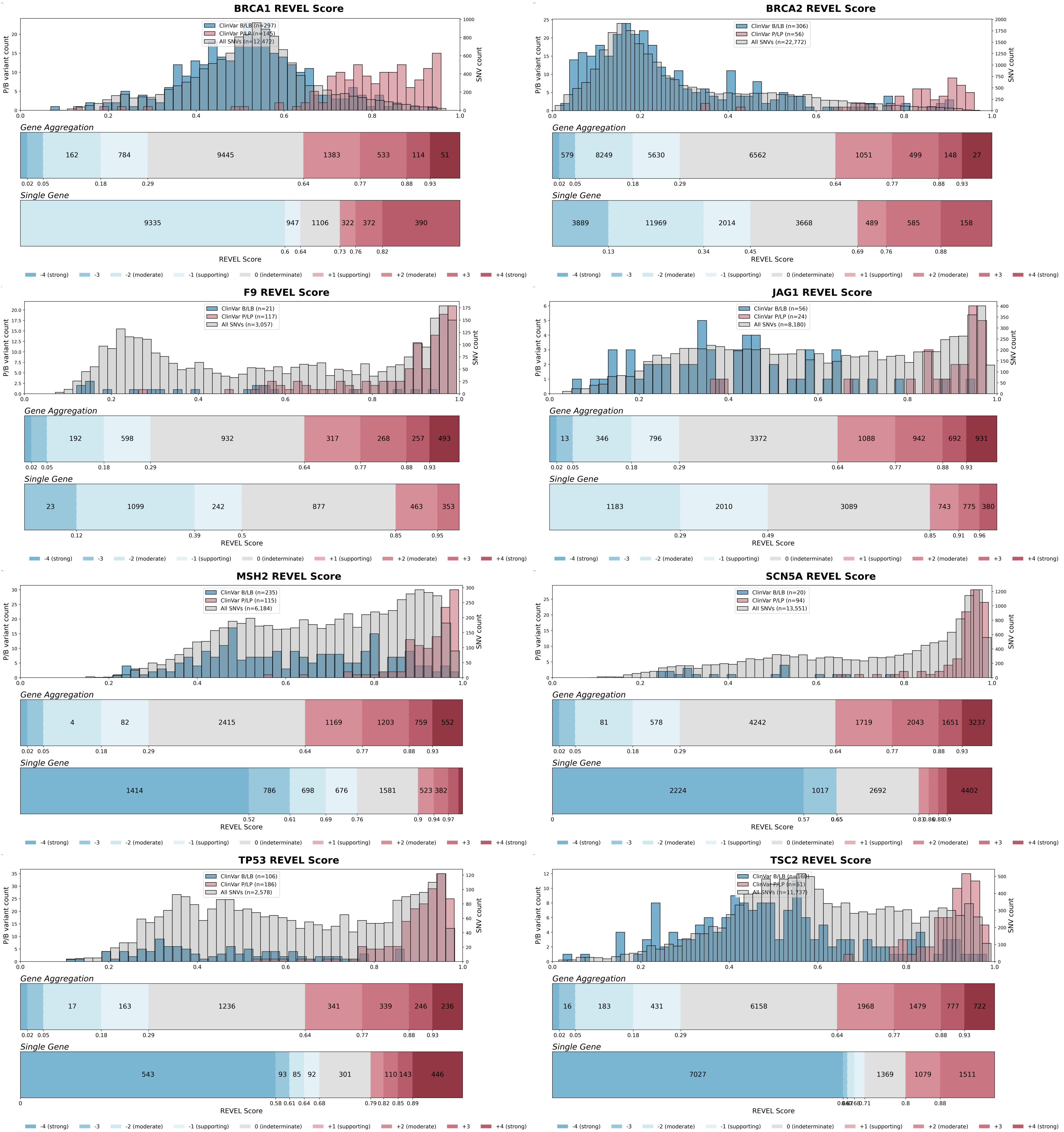
BRCA1 REVEL Score
ClinVar B/LB (n=297)
ClinVar P/LP (n=145)
All SNVs (n=12,472)
Gene Aggregation
162
784
9445
1383
533
114
51
Single Gene
9335
947
1106
322
372
390
BRCA2 REVEL Score
ClinVar B/LB (n=306)
ClinVar P/LP (n=56)
All SNVs (n=22,772)
Gene Aggregation
579
8249
5630
6562
1051
499
148
27
Single Gene
3889
11969
2014
3668
489
585
158
F9 REVEL Score
ClinVar B/LB (n=21)
ClinVar P/LP (n=117)
All SNVs (n=3,057)
Gene Aggregation
192
598
932
317
268
257
493
Single Gene
23
1099
242
877
463
353
JAG1 REVEL Score
ClinVar B/LB (n=56)
ClinVar P/LP (n=24)
All SNVs (n=8,180)
Gene Aggregation
13
346
796
3372
1088
942
692
931
Single Gene
1183
2010
3089
743
775
380
MSH2 REVEL Score
ClinVar B/LB (n=235)
ClinVar P/LP (n=115)
All SNVs (n=6,184)
Gene Aggregation
4
82
2415
1169
1203
759
552
Single Gene
1414
786
698
676
1581
523
382
SCN5A REVEL Score
ClinVar B/LB (n=20)
ClinVar P/LP (n=94)
All SNVs (n=13,551)
Gene Aggregation
81
578
4242
1719
2043
1651
3237
Single Gene
2224
1017
2692
4402
TP53 REVEL Score
ClinVar B/LB (n=106)
ClinVar P/LP (n=186)
All SNVs (n=2,578)
Gene Aggregation
17
163
1236
341
339
246
236
Single Gene
543
93
85
92
301
110
143
446
TSC2 REVEL Score
ClinVar B/LB (n=160)
ClinVar P/LP (n=61)
All SNVs (n=11,737)
Gene Aggregation
16
183
431
6158
1968
1479
777
722
Single Gene
7027
1369
1079
1511
P/B variant count
SNV count
REVEL Score
-4 (strong)
-3
-2 (moderate)
-1 (supporting)
0 (indeterminate)
+1 (supporting)
+2 (moderate)
+3
+4 (strong)

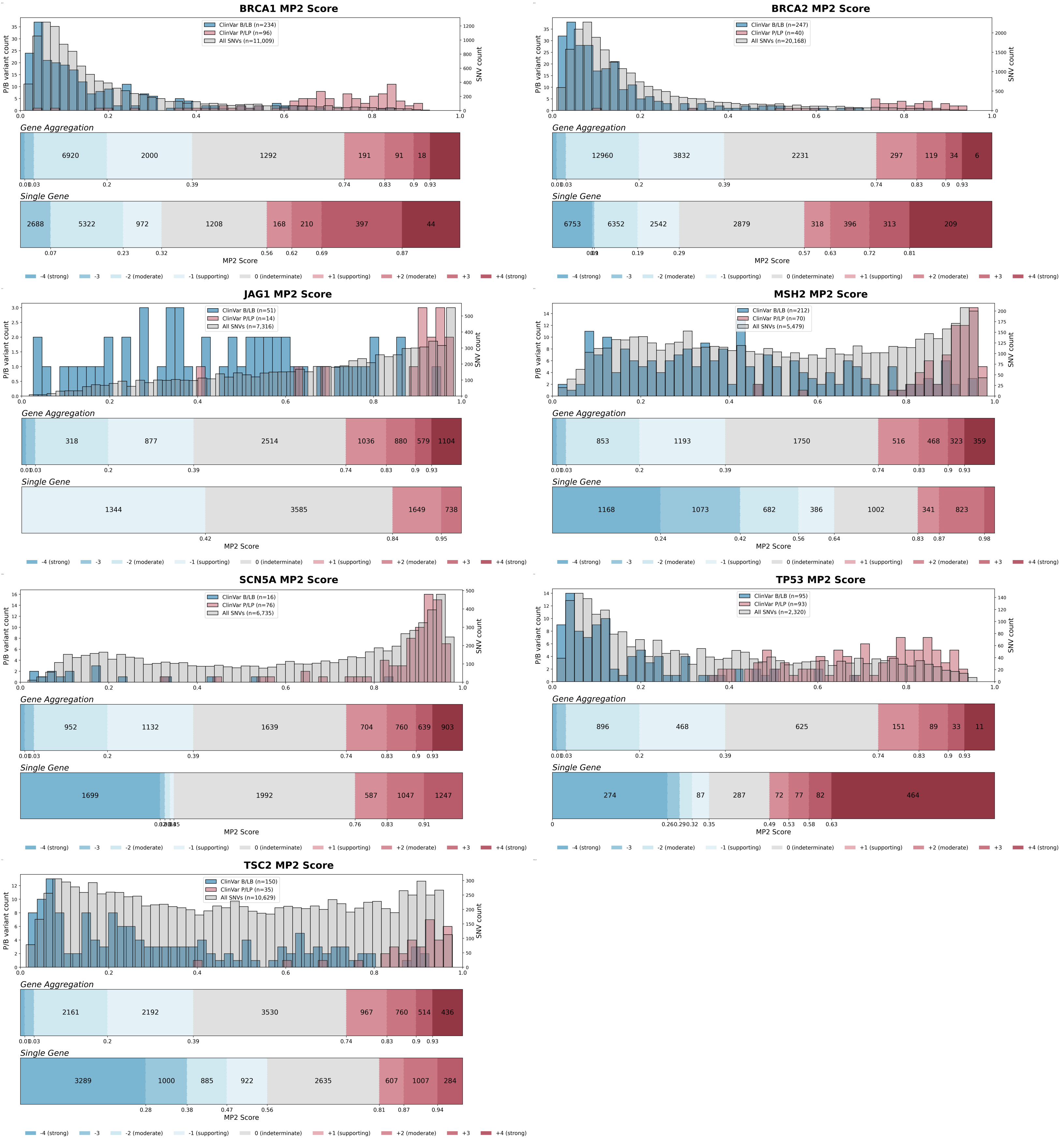
BRCA1 MP2 Score
ClinVar B/LB (n=234)
ClinVar P/LP (n=96)
All SNVs (n=11,009)
P/B variant count
SNV count
Gene Aggregation
6920
2000
1292
191
91
18
0.01
0.03
0.2
0.39
0.74
0.83
0.9
0.93
Single Gene
2688
5322
972
1208
168
210
397
44
0.07
0.23
0.32
0.56
0.62
0.69
0.87
MP2 Score
-4 (strong)
-3
-2 (moderate)
-1 (supporting)
0 (indeterminate)
+1 (supporting)
+2 (moderate)
+3
+4 (strong)
BRCA2 MP2 Score
ClinVar B/LB (n=247)
ClinVar P/LP (n=40)
All SNVs (n=20,168)
12960
3832
2231
297
119
34
6
6753
6352
2542
2879
318
396
313
209
0.19
0.29
0.57
0.63
0.72
0.81
JAG1 MP2 Score
ClinVar B/LB (n=51)
ClinVar P/LP (n=14)
All SNVs (n=7,316)
318
877
2514
1036
880
579
1104
1344
3585
1649
738
0.42
0.84
0.95
MSH2 MP2 Score
ClinVar B/LB (n=212)
ClinVar P/LP (n=70)
All SNVs (n=5,479)
853
1193
1750
516
468
323
359
1168
1073
682
386
1002
341
823
0.24
0.42
0.56
0.64
0.83
0.87
0.98
SCN5A MP2 Score
ClinVar B/LB (n=16)
ClinVar P/LP (n=76)
All SNVs (n=6,735)
952
1132
1639
704
760
639
903
1699
1992
587
1047
1247
0.76
0.83
0.91
TP53 MP2 Score
ClinVar B/LB (n=95)
ClinVar P/LP (n=93)
All SNVs (n=2,320)
896
468
625
151
89
33
11
274
87
287
72
77
82
464
0.26
0.29
0.32
0.35
0.49
0.53
0.58
0.63
TSC2 MP2 Score
ClinVar B/LB (n=150)
ClinVar P/LP (n=35)
All SNVs (n=10,629)
2161
2192
3530
967
760
514
436
3289
1000
885
922
2635
607
1007
284
0.28
0.38
0.47
0.56
0.81
0.87
0.94

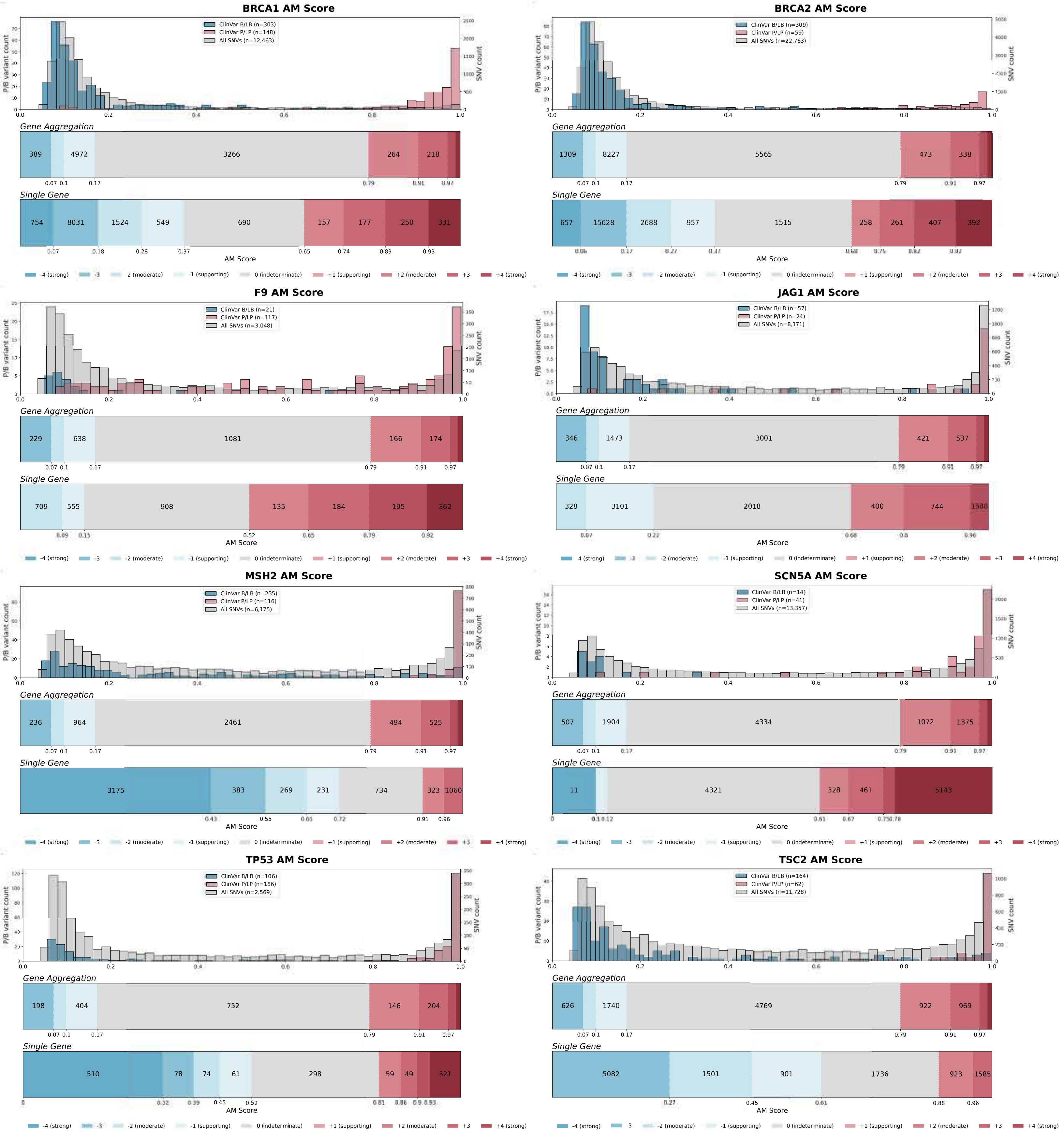
BRCA1 AM Score
ClinVar B/LB (n=303)
ClinVar P/LP (n=148)
All SNVs (n=12,463)
P/B variant count
SNV count
Gene Aggregation
389 4972 3266 264 218
0.07 0.1 0.17 0.79 0.91 0.97
Single Gene
754 8031 1524 549 690 157 177 250 331
0.07 0.18 0.28 0.37 0.65 0.74 0.83 0.93
AM Score
-4 (strong) -3 -2 (moderate) -1 (supporting) 0 (indeterminate) +1 (supporting) +2 (moderate) +3 +4 (strong)
BRCA2 AM Score
ClinVar B/LB (n=309)
ClinVar P/LP (n=59)
All SNVs (n=22,763)
Gene Aggregation
1309 8227 5565 473 338
0.07 0.1 0.17 0.79 0.91 0.97
Single Gene
657 15628 2688 957 1515 258 261 407 392
0.06 0.17 0.27 0.37 0.68 0.75 0.82 0.92
AM Score
F9 AM Score
ClinVar B/LB (n=21)
ClinVar P/LP (n=117)
All SNVs (n=3,048)
Gene Aggregation
229 638 1081 166 174
0.07 0.1 0.17 0.79 0.91 0.97
Single Gene
709 555 908 135 184 195 362
0.09 0.15 0.52 0.65 0.79 0.92
AM Score
JAG1 AM Score
ClinVar B/LB (n=57)
ClinVar P/LP (n=24)
All SNVs (n=8,171)
Gene Aggregation
346 1473 3001 421 537
0.07 0.1 0.17 0.79 0.91 0.97
Single Gene
328 3101 2018 400 744 1580
0.07 0.22 0.68 0.8 0.96
AM Score
MSH2 AM Score
ClinVar B/LB (n=235)
ClinVar P/LP (n=116)
All SNVs (n=6,175)
Gene Aggregation
236 964 2461 494 525
0.07 0.1 0.17 0.79 0.91 0.97
Single Gene
3175 383 269 231 734 323 1060
0.43 0.55 0.65 0.72 0.91 0.96
AM Score
SCN5A AM Score
ClinVar B/LB (n=14)
ClinVar P/LP (n=41)
All SNVs (n=13,357)
Gene Aggregation
507 1904 4334 1072 1375
0.07 0.1 0.17 0.79 0.91 0.97
Single Gene
11 4321 328 461 5143
0 0.1 0.12 0.61 0.67 0.75 0.78
AM Score
TP53 AM Score
ClinVar B/LB (n=106)
ClinVar P/LP (n=186)
All SNVs (n=2,569)
Gene Aggregation
198 404 752 146 204
0.07 0.1 0.17 0.79 0.91 0.97
Single Gene
510 78 74 61 298 59 49 521
0 0.32 0.39 0.45 0.52 0.81 0.86 0.9 0.93
AM Score
TSC2 AM Score
ClinVar B/LB (n=164)
ClinVar P/LP (n=62)
All SNVs (n=11,728)
Gene Aggregation
626 1740 4769 922 969
0.07 0.1 0.17 0.79 0.91 0.97
Single Gene
5082 1501 901 1736 923 1585
0.27 0.45 0.61 0.88 0.96
AM Score

### Extended Data Figure 3: Gene-specific predictor calibrations across 8 genes for REVEL, MutPred2, and AlphaMissense

Gene-specific calibrations for REVEL, MutPred2, and AlphaMissense, respectively, across 8 genes, each visualized with three components: top, pathogenic (red) and benign (blue) control missense variants from the ClinVar January 2025 release overlaid with all possible missense SNVs (light grey); middle, genome-wide aggregation score thresholds; bottom, gene-specific calibration score intervals, with both middle and bottom panels labeled by evidence strength (up to 4 pathogenic or benign points) and SNV counts shown within sufficiently large bins. Calibrations of MutPred2 scores for F9 are not shown because after excluding variants used in MutPred2 training, the remaining control variants were insufficient to support a robust gene-specific calibration.
